# Enhancing the hyaline cartilage regenerative potential of induced pluripotent stem cell-derived chondrocyte cell sheets

**DOI:** 10.64898/2026.09.25.754531

**Authors:** Takumi Takahashi, Ryoka Uchiyama, Eriko Toyoda, Miki Maehara, Yoshiko Takahashi, Shiho Wasai, Haruka Omura, Makoto Ogawa, Akihiro Yamashita, Masahiko Watanabe, Noriyuki Tsumaki, Masato Sato

## Abstract

**Introduction:** Autologous and allogeneic chondrocyte cell sheet transplantation has shown safety and potential efficacy for cartilage defects associated with knee osteoarthritis. Although polydactyly-derived chondrocyte (PDC) cell sheets have been used clinically for allogeneic transplantation, a scalable and reliable alternative cell source is needed. This study aimed to establish a proof-of-concept protocol for generating chondrocyte cell sheets from clinical-grade human induced pluripotent stem cells (iPS cells).

**Methods:** Cartilaginous tissue was generated from human iPS cells, and isolated chondrocytes were used to fabricate iPS cell-derived chondrocyte sheets. In Phase 1, sheets were produced in high-serum medium under normoxic, hypoxic, or hypoxic/high-seeding-density conditions and compared with PDC sheets using a three-dimensional in vitro model and xenogeneic orthotopic transplantation in an immunodeficient rat model. In Phase 2, sheets were produced in low-serum medium under hypoxic/high-seeding-density conditions and evaluated for cartilage-related gene expression, secreted factors, cell surface marker expression, and in vivo cartilage repair.

**Results:** In Phase 1, hypoxic/high-seeding-density culture enhanced in vitro chondrogenic properties. However, after transplantation, iPS cell-derived chondrocyte sheets formed fibrotic repair tissue, whereas PDC sheets generated hyaline-like cartilage. In Phase 2, low-serum culture increased expression of cartilage-related genes, including collagen type II alpha 1 (COL2A1) and SRY-box transcription factor 9 (SOX9), enhanced secretion of transforming growth factor beta 1 and melanoma inhibitory activity, and upregulated CD9, CD26, CD56, CD201, CD227, and GD2. Low-serum iPS cell-derived chondrocyte sheets regenerated hyaline cartilage at 4 weeks, and the repair tissue was maintained at 12 weeks without bone formation.

**Conclusions:** A robust fabrication protocol enabled human iPS cell-derived chondrocyte sheets to regenerate hyaline-like cartilage *in vivo*. These findings support their potential as a scalable cell source for clinical cartilage repair.

**HIGHLIGHTS:**

- Chondrocyte sheets were fabricated from clinical-grade human iPS cells.
- Hypoxia and high seeding density improved *in vitro* chondrogenicity but not *in vivo*.
- iPSC sheet fabricated under hypoxia, high seeding density, and low-serum regenerated hyaline-like cartilage *in vivo*.
- A protocol for iPSC sheet fabrication with translational potential was established.

## 1. INTRODUCTION

Osteoarthritis of the knee (OAK) is a progressive degenerative disease that has a significant impact on quality of life^1^. OAK is characterized as a disease of the whole joint, accompanied by the degeneration of articular cartilage, inflammation of the synovium, and formation of osteophytes, eventually leading to pain and disability^2^. Current treatment methods, such as hyaluronic acid injections, microfracture, osteochondral allograft transplantation, and autologous chondrocyte implantation, cannot completely reverse the progress of degeneration and fail to address the large cartilage defects found in OAK^3^. Thus, treatment methods to regenerate hyaline cartilage, the native type of cartilage found in articular cartilage, are actively being developed^4,5^.

Cell sheet technology^6,7^ has been applied to cartilage regeneration through the use of both autologous chondrocytes (ACs) from the adult patients’ knee joint^8^ and allogeneic chondrocytes from the surgical remains of juvenile patients with polydactyly^9^. For cell sheet fabrication, chondrocytes are enzymatically isolated from adult knee articular cartilage or juvenile polydactyly-derived cartilage and subsequently expanded in monolayer on temperature-responsive culture dishes. A first-in-human clinical study using AC sheets confirmed the safety of the treatment method and their potential to regenerate hyaline cartilage in patients with OAK^10^. Preclinical studies using both rat^11,12^ and rabbit^13,14^ xenogeneic transplantation models have further confirmed the efficacy of both AC sheets and polydactyly-derived chondrocyte (PDC) sheets to regenerate hyaline cartilage in the knee joint. With such evidence, a first-in-human clinical study using allogeneic PDC sheets was conducted, and the results demonstrated the safety of the allogeneic treatment method^15^.

However, several challenges remain: (1) the number of polydactyl donors is limited^16^; (2) only a limited amount of cartilage can be harvested; therefore, PDCs must be expanded through passaging, which may reduce the regenerative properties of PDC sheets^14,17^; and (3) donor selection is required to ensure safety and efficacy of PDC sheets^14^. Thus, an alternative allogeneic cell source that can address such issues is of significant interest.

One such cell source may be induced pluripotent stem (iPS) cells. In 2013, the iPS Cell Stock Project, located at the Center for iPS Cell Research and Application (CiRA), Kyoto University in Japan, was initiated to develop clinical-grade human iPS cell lines^18,19^. The iPS Cell Stock Project currently provides clinical-grade cell lines of human leukocyte antigen (HLA) homozygous iPS cells made from either peripheral blood mononuclear cells (PBMCs) or cord blood cells. As such, iPS cell lines that can be directly translated to the clinical setting provide a logical choice as a novel allogeneic cell source.

Several studies have reported on the successful fabrication of hyaline cartilage from iPS cells^20–22^. In particular, in 2015, Yamashita et al. reported a scaffold-less method for fabricating iPS cell-derived cartilaginous tissue (iPS-Cart) that exhibits hyaline cartilage properties^23^. The transplantation of iPS-Carts to rat and minipig osteochondral defect models showed good integration of iPS-Carts to cartilage defect areas and regeneration of hyaline cartilage in the knee joint^23^. iPS-Carts fabricated from iPS cells developed through the iPS Cell Stock Project are being used in a clinical study (jRCTa050190104). However, because iPS-Carts are small and spheroidal in shape, requiring structural anchors to patients’ surviving cartilage tissue, they are not immediately applicable to the large cartilage defects associated with OAK.

Accordingly, we proposed integrating the iPS-Cart and cell sheet technologies to develop iPS cell-derived chondrocyte (iPSC) sheets for the treatment of OAK. In a preliminary study, we attempted to fabricate iPSC sheets using cells isolated from iPS-Carts, using a protocol similar to that of fabricating AC sheets. This protocol involved fabricating iPSC sheets under normoxic conditions with high-serum (HS)^10^. However, when tested in a rabbit xenogeneic transplantation model, iPSC sheets failed to regenerate hyaline cartilage (Supplementary Fig. 1). This outcome highlighted the need for further optimization of the iPSC sheet fabrication process to enhance its cartilage regeneration potential.

Numerous methods have been investigated to maintain the chondrogenic phenotype of chondrocytes in monolayer culture. A comparative study of seeding densities in monolayer culture revealed that higher densities (∼2.5 × 10□ cells/cm²) better preserve chondrogenic phenotype of articular chondrocytes compared to lower densities (∼2.5 × 10³ cells/cm²)^24^. The application of hypoxic conditions during monolayer culture of bovine and human articular chondrocytes has been shown to enhance glycosaminoglycan production and upregulate chondrogenic expression^25,26^. Additionally, serum-free or serum-reduced approaches have been employed in monolayer culture^27–29^, with a recent study revealing that lipid scarcity plays an important role in enhanced chondrogenic commitment^30^. These studies demonstrate that manipulating culture conditions, including cell density, oxygen tension, and media composition, can significantly influence the maintenance of chondrogenic phenotype in monolayer culture, offering potential strategies to improve the fabrication of iPSC sheets.

The objective of this study was to establish proof-of-concept through (1) improving the properties of iPSC sheets *in vitro* and (2) demonstrating *in vivo* efficacy using xenogeneic orthotopic transplantation in a rat immunodeficient model^11^. To accelerate the translational potential of iPSC sheets, human iPS cells from the iPS Cell Stock Project^19^ were used to make iPS-Carts^23^, which in turn were used to fabricate iPSC sheets. The study encompassed two phases. We first hypothesized that hypoxic culture and high seeding densities would improve the regenerative properties of iPSC sheets. In Phase 1, iPSC sheets were fabricated using HS media as used in previous clinical studies and cultured in hypoxia and/or high seeding density conditions. The regenerative properties of iPSC sheets were compared to PDC sheets through xenogeneic transplantations. Based on the Phase 1 results, to further improve the regenerative properties of iPSC sheets, in Phase 2, it was hypothesized that a culture medium reduced in serum, in combination with hypoxia and high seeding density, would be beneficial for maintaining the chondrogenic properties of chondrocytes obtained from iPS-Carts. The regenerative properties of iPSC sheets fabricated under HS or low-serum (LS) conditions were characterized through *in vitro* and *in vivo* evaluations. The results support the development of iPSC sheets toward cartilage repair in patients with OAK.

## 2. RESULTS

### 2.1 Phase 1: Hypoxia and high seeding density improve the chondrogenic potential of iPSC sheets

To improve the regenerative properties of iPSC sheets, the effects of hypoxia and high seeding densities were evaluated. In Phase 1, iPSC sheets were fabricated under three conditions: (1) normoxia, seeded at 1 × 10^4^ cells/cm^2^ (iPSC N1), (2) hypoxia, seeded at 1 × 10^4^ cells/cm^2^ (iPSC H1), and (3) hypoxia seeded at 5 × 10^4^ cells/cm^2^ (iPSC H5). The PDC sheets were fabricated according to previously established protocols from three donors (i.e., PDC A, PDC B, and PDC C). The *in vitro* properties of iPSC sheets were compared to those of PDC sheets. Macroscopic (Fig. 1a, Supplementary Fig. 2) and histological (Fig. 1b, Supplementary Fig. 3) images revealed similar characteristics between PDC sheets and iPSC sheets. Immunohistochemistry showed positive staining for aggrecan (ACAN), collagen type I (COL1), and fibronectin (FN), and negative staining for collagen type II (COL2), indicating a dedifferentiated state during monolayer culture (Fig. 1b, Supplementary Fig. 3). PDC sheets of donors A, B, and C contained an average of 2.9, 2.7, and 3.7 × 10^6^ cells per sheet with an average thickness of 10.3, 11.9, and 17.1 μm, respectively (Fig. 1c, Supplementary Fig. 3). Compared to iPSC H1 sheets, iPSC N1 sheets contained a significantly higher number of cells (3.0 × 10^6^ cells vs 1.9 × 10^6^ cells). Compared to iPSC H1 sheets, iPSC H5 sheets had a significantly higher number of cells (2.2 × 10^6^ cells vs 1.9 × 10^6^ cells) (Fig. 1c). The thickness was highest for iPSC H5 sheets (29.1 μm), followed by iPSC N1 sheets (19.4 μm) and iPSC H1 sheets (13.2 μm) (Fig. 1c, Supplementary Fig. 3).

**Fig. 1.**
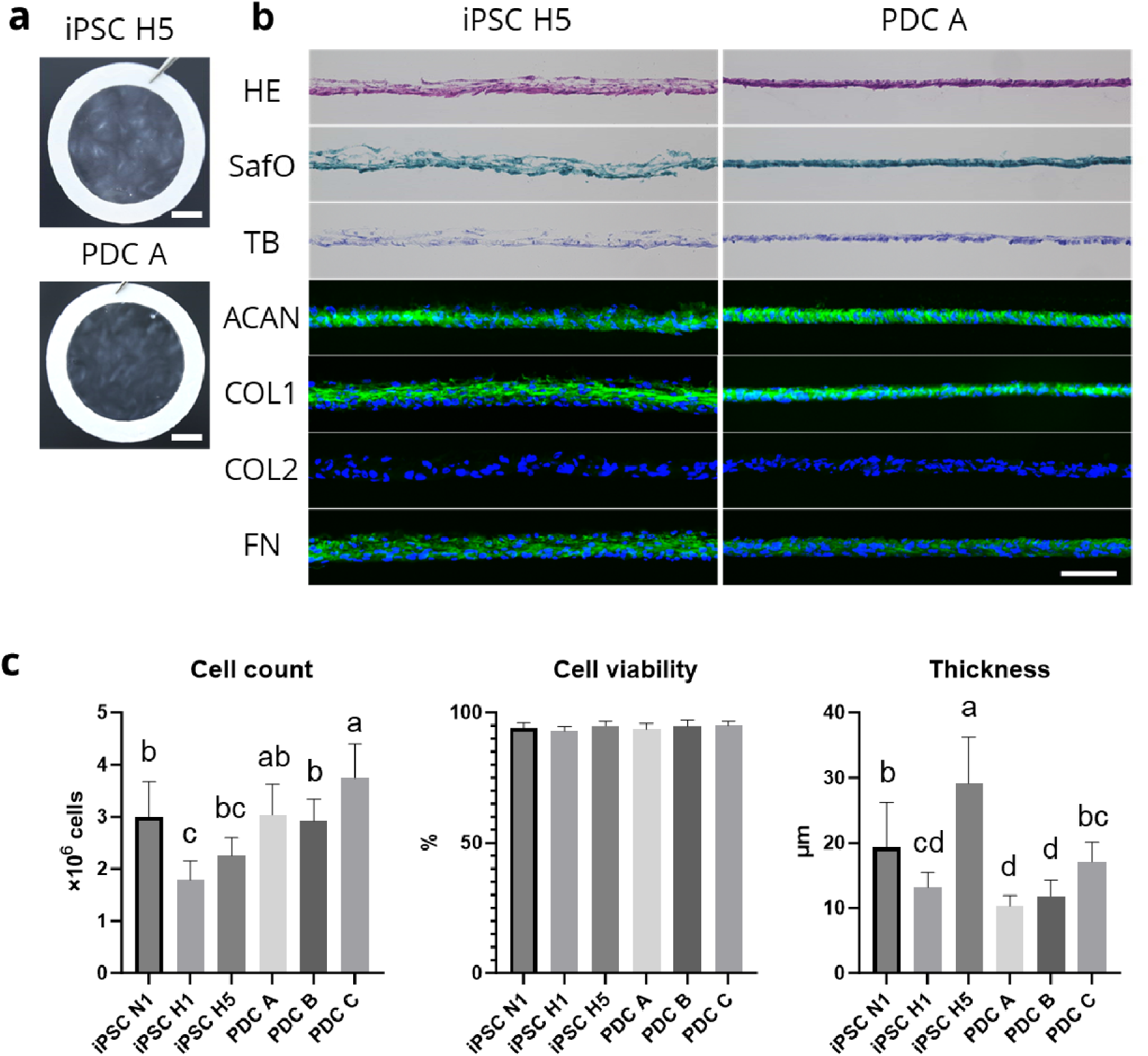
Structural properties of iPSC sheets and PDC sheets. (**a**) Macroscopic and (**b**) histological analyses of iPSC sheets and PDC sheets show that both sheets exhibit thin sheet structures that can be picked up using a PVDF membrane and express ACAN, COL1, and FN, indicating a dedifferentiated chondrocyte phenotype. (**c**) Cell count, cell viability, and thickness indicate a high density of viable cells present in the sheets, with iPSC H5 sheets having the highest thickness. Lower-case letters of the connecting letters report show that groups not sharing the same letter are significantly different. Scale bar = (**a**) 3 mm and (**b**) 100 µm. PVDF, polyvinylidene difluoride; HE, hematoxylin and eosin; SafO, safranin O; TB, toluidine blue; ACAN, aggrecan; COL1, collagen type I; COL2, collagen type II; FN, fibronectin.

The secretion levels of transforming growth factor β1 (TGF-β1) were higher for iPSC sheets, while those of melanoma inhibitory activity (MIA), a secretory protein used as an indicator of chondrogenic differentiation^31,14^, did not differ significantly between iPSC sheets and PDC sheets (Fig. 2a). A three-dimensional (3D) culture of chondrocyte sheets was used to predict their chondrogenicity. Gene expression analysis of 3D cultured sheets at 4 weeks revealed a significantly higher expression of *COL2A1* for iPSC H5 compared to iPSC N1, while the expression of *COL2A1* varied among PDC sheets of different donors (Fig. 2b).

**Fig. 2.**
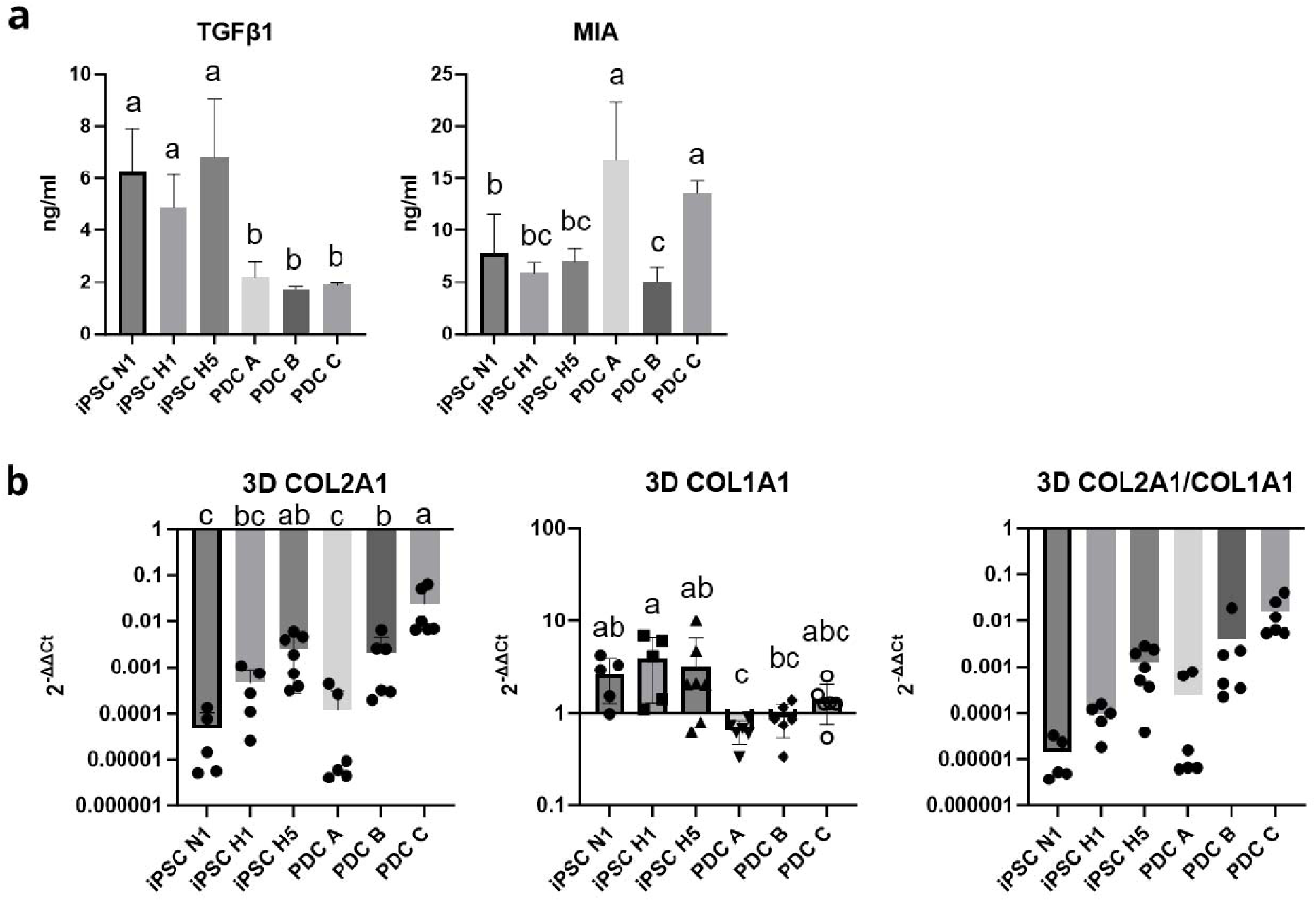
iPSC sheets form fibrotic tissue *in vitro*. Analyses of humoral factors and chondrogenic potential of iPSC sheets and PDC sheets. (**a**) Compared to PDC sheets, iPSC sheets produced significantly higher levels of TGF-β1, while PDC A and PDC C produced significantly higher levels of MIA compared to other sheets. (**b**) Chondrogenic potential was evaluated through gene expression analysis after 3D culture of the sheets. Combination of hypoxia and high seeding density significantly increased expression of *COL2A1* as well as improving *COL2A1* to *COL1A1* ratios. Lower-case letters of the connecting letters report show that groups not sharing the same letter are significantly different. TGF-β1 = transforming growth factor beta 1; MIA = melanoma inhibitory activity.

Using a previously established xenogeneic orthotopic transplantation model^11^, iPSC sheets fabricated under the three different conditions and PDC sheets from the three donors were evaluated *in vivo*. Histological evaluation showed that compared to the no-treatment (NT) group, in the iPSC sheet groups, staining for safranin O (SafO) and COL2 was minimal, while repair tissue stained strongly for COL1 (Figs. 3 and 4, Supplementary Figs. 4-15). COL2 and COL1 staining were used as an indicator of hyaline-like cartilage and fibrocartilage, respectively. Immunohistochemical expression for human-specific vimentin (hVim) indicated good engraftment of human cells, confirming that the repair tissue was formed by the transplanted human sheets. PDC sheets for all three donors demonstrated defect filling with repair tissue that stained strongly for SafO, toluidine blue (TB), and COL2, while the surfaces of the repair tissue stained for COL1, indicating mostly a hyaline-like cartilaginous repair tissue. Staining for hVim also indicated that human cells successfully engrafted to the recipient tissue. Histological scoring revealed that PDC sheets for all donors had significantly higher scores compared to the NT group, while iPSC sheets did not show a significant improvement compared to the NT group (Fig. 5).

**Fig. 3.**
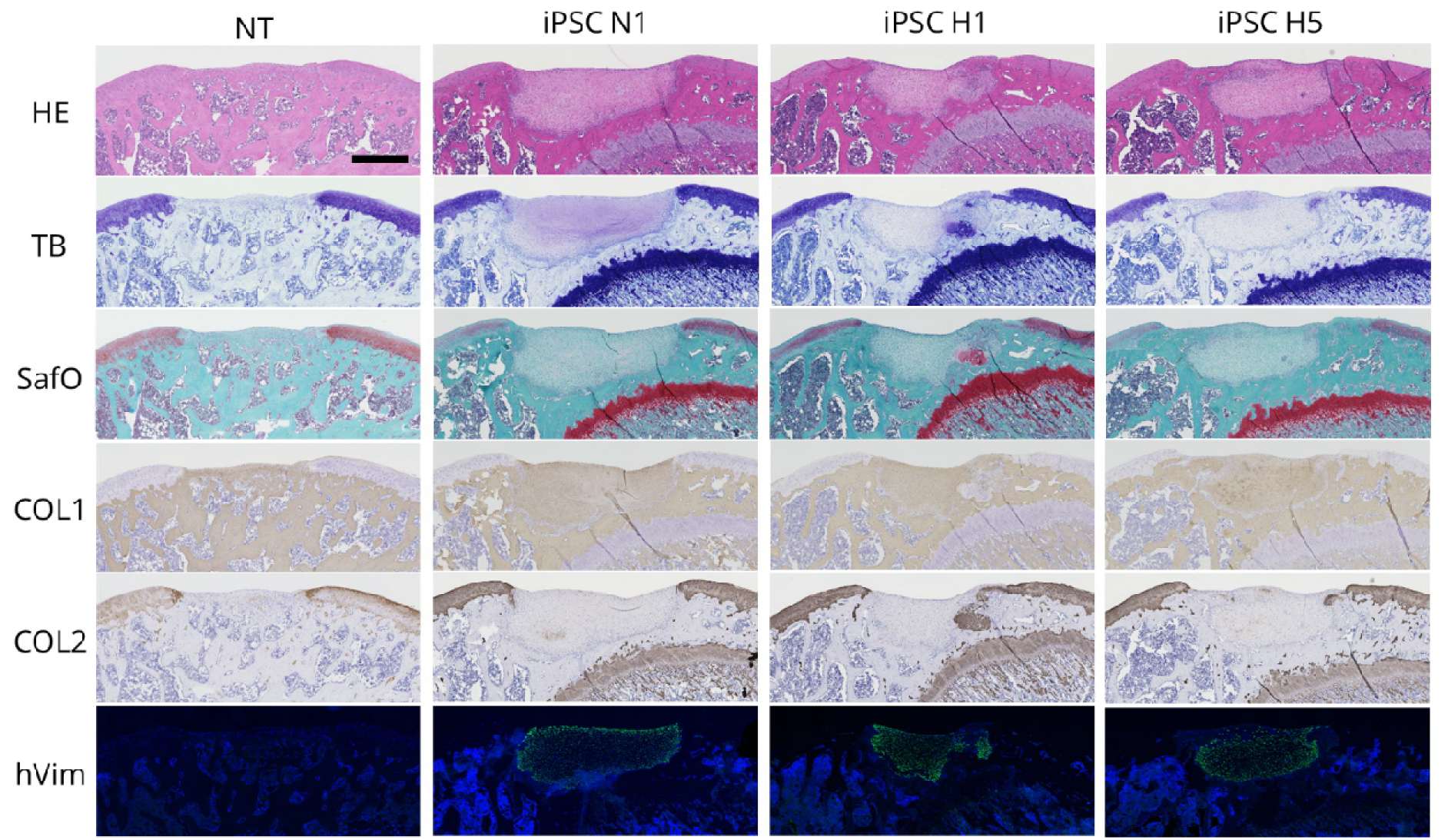
Histological evaluation at 4 weeks after transplantation of iPSC sheets in a xenogeneic orthotopic transplantation model using athymic nude rats. The NT group showed bone-like formation in the defect area with slightly lower staining of TB, SafO, and COL2. The transplantation of iPSC N1, H1, and H5 sheets demonstrated the formation of repair tissue that stained strongly for COL1 indicating a non-hyaline, fibrotic cartilage-like tissue formation. Staining for hVim showed that the transplanted cells filled the defect area and remained up to 4 weeks after transplantation. n = 6 per group. Scale bar = 500 µm. NT, no-treatment; HE, hematoxylin and eosin; SafO, safranin O; TB, toluidine blue; COL1, collagen type I; COL2, collagen type II; hVim, human-specific vimentin.

**Fig. 4.**
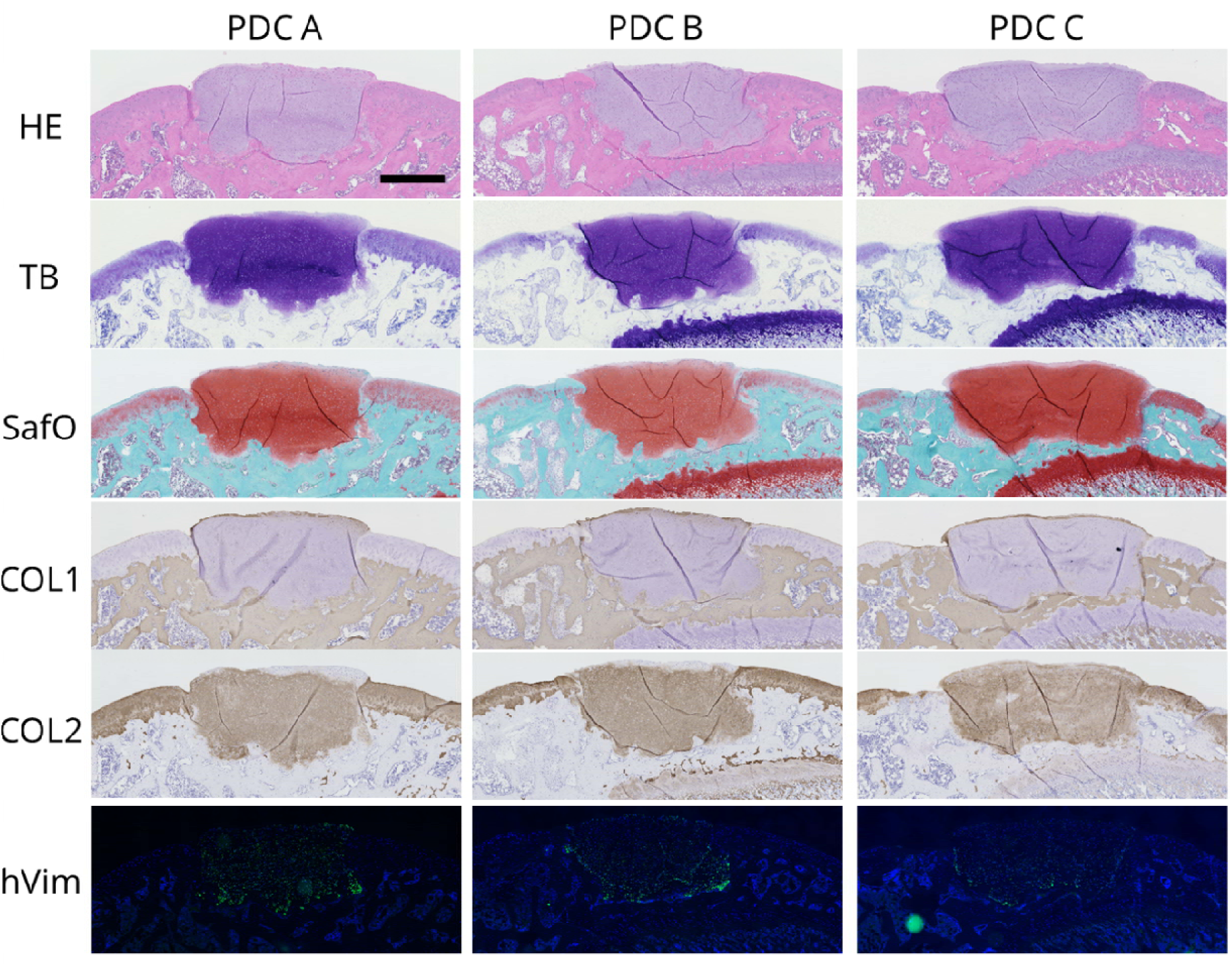
Histological evaluation at 4 weeks after transplantation of PDC sheets in a xenogeneic orthotopic transplantation model using athymic nude rats. The repair tissue of all three donors stained strongly for TB, SafO, and COL2, suggesting repair with hyaline-like cartilage tissue. Staining for COL1 to indicate fibrocartilage was limited to mainly the repair tissue surfaces. Staining for hVim indicated that the transplanted cells filled parts of the defect area and remained 4 weeks after transplantation. n = 6 per group. Scale bar = 500 µm. HE, hematoxylin and eosin; SafO, safranin O; TB, toluidine blue; COL1, collagen type I; COL2, collagen type II; hVim, human-specific vimentin.

**Fig. 5.**
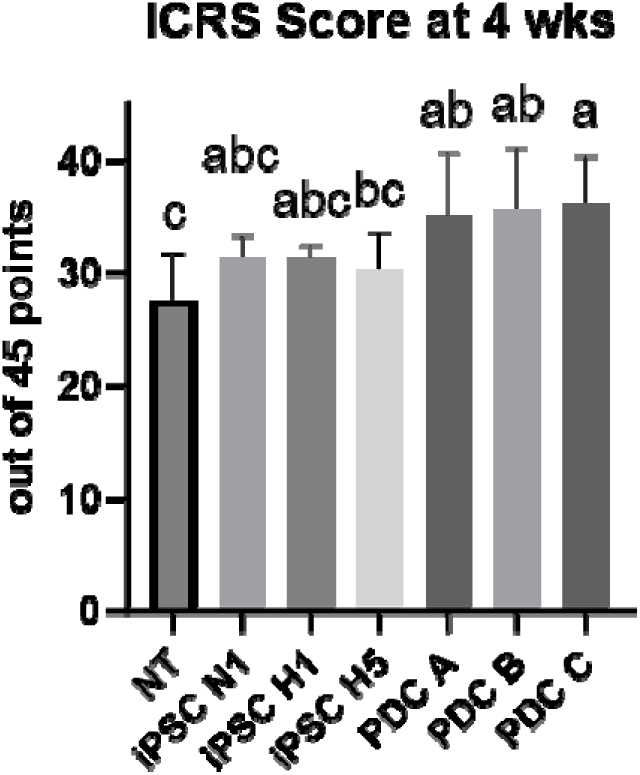
Comparison of repair tissue characteristics based on International Cartilage Regeneration and Joint Preservation Society (ICRS) scores at 4 weeks after transplantation of iPSC sheets and PDC sheets in a xenogeneic orthotopic transplantation model using athymic nude rats. Compared with the no-treatment (NT) group, only the PDC sheets had significantly higher ICRS scores. While iPSC sheets filled defect areas and formed repair tissue, the fibrotic nature of the repair tissue led to lower ICRS scores. Lower-case letters of the connecting letters report show that groups not sharing the same letter are significantly different.

### 2.2 Phase 2: Low-serum medium further improves the chondrogenic potential of iPSC sheets

Based on *in vivo* results in Phase 1, further modification to protocols for iPSC sheet fabrication was warranted. It was hypothesized that a culture medium reduced in serum would be beneficial for maintaining the chondrogenic properties of chondrocytes obtained from iPS-Carts. Thus, in combination with hypoxia and high seeding density condition used in Phase 1, low-serum (LS) medium was used to fabricate iPSC LS sheets. The iPSC LS sheets were compared to iPSC sheets fabricated using the HS medium, i.e., iPSC HS sheets.

The microscopic images of cultured chondrocytes observed on day 14 showed a polygonal shape for chondrocytes of iPSC LS sheets, while a fibroblastic shape was observed for chondrocytes of iPSC HS sheets (Supplementary Fig. 16). Both types of iPSC sheets were able to be detached using polyvinylidene difluoride (PVDF) membrane rings without tearing or shrinking (Fig. 6a, Supplementary Fig. 2). Histological analysis revealed that both types of iPSC sheets exhibited metachromasia with TB staining, while both were negative for SafO staining. Immunohistological analysis revealed that both types of iPSC sheets were positive for ACAN, COL1, and FN, while both were negative for COL2 (Fig. 6b). iPSC HS sheets contained a significantly higher (*p* = 0.0064) number of cells per sheet (3.2 ± 0.8 × 10^6^ cells) than iPSC LS sheets (1.4 ± 0.2 × 10^6^ cells) (Fig. 6c). Both types of iPSC sheets had high cell viability (*p* = 0.57; HS 93.7 ± 1.7% vs. LS 94.2 ± 1.7%; Fig. 6c). The thickness of iPSC HS sheets (21.6 ± 3.3 μm) was significantly greater (*p* = 0.00093) compared with iPSC LS sheets (16.0 ± 2.7 μm) (Fig. 6c). iPSC LS sheets produced a significantly higher level of TGF-β1 (*p* = 0.0013; HS 7.7 ± 1.7 ng vs. LS 27.6 ± 8.6 ng) and MIA (*p* = 0.00017; HS 23.2 ± 8.6 ng vs. LS 80.0 ± 15.1 ng) per sheet during 72 h (Fig. 7a). Furthermore, in 3D culture, no significant difference was detected for the expression of *COL2A1*, but the expression of *COL1A1* was significantly higher for iPSC LS sheets (Fig. 7b).

**Fig. 6.**
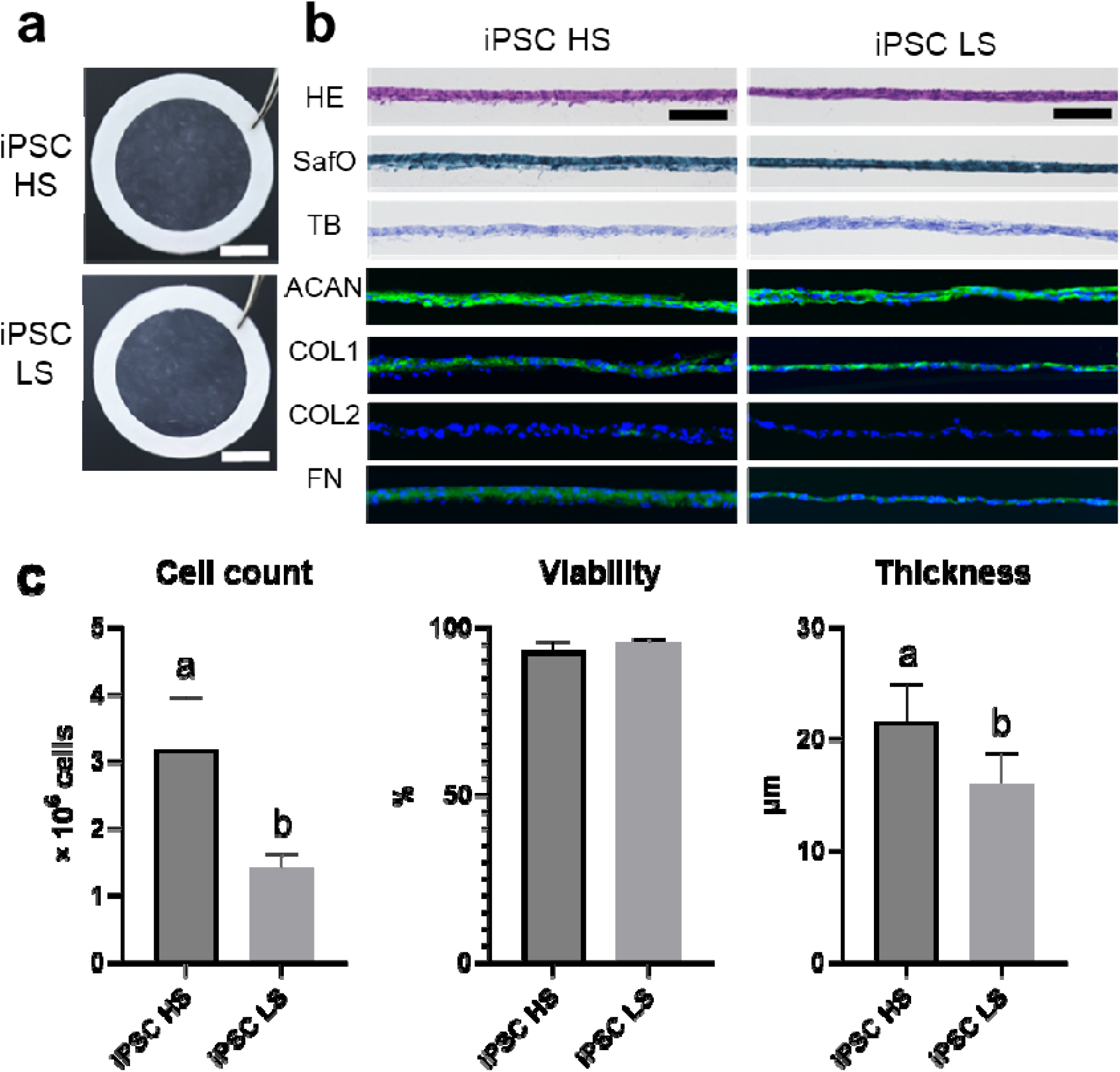
Structural properties of iPSC HS sheets and iPSC LS sheets. (**a**) Macroscopic and (**b**) histological analyses of iPSC HS sheets and LS sheets show that both sheets exhibit thin sheet structures that can be picked up using a PVDF membrane and express ACAN, COL1, and FN, indicating a dedifferentiated chondrocyte phenotype. (**c**) iPSC HS sheets compared to iPSC LS sheets had significantly higher cell count as well as thickness. Lower-case letters of the connecting letters report show that groups not sharing the same letter are significantly different. Scale bar = (**a**) 5 mm and (**b**)100 µm. PVDF, polyvinylidene difluoride; HE, hematoxylin and eosin; SafO, safranin O; TB, toluidine blue; ACAN, aggrecan; COL1, collagen type I; COL2, collagen type II; FN, fibronectin.

**Fig. 7.**
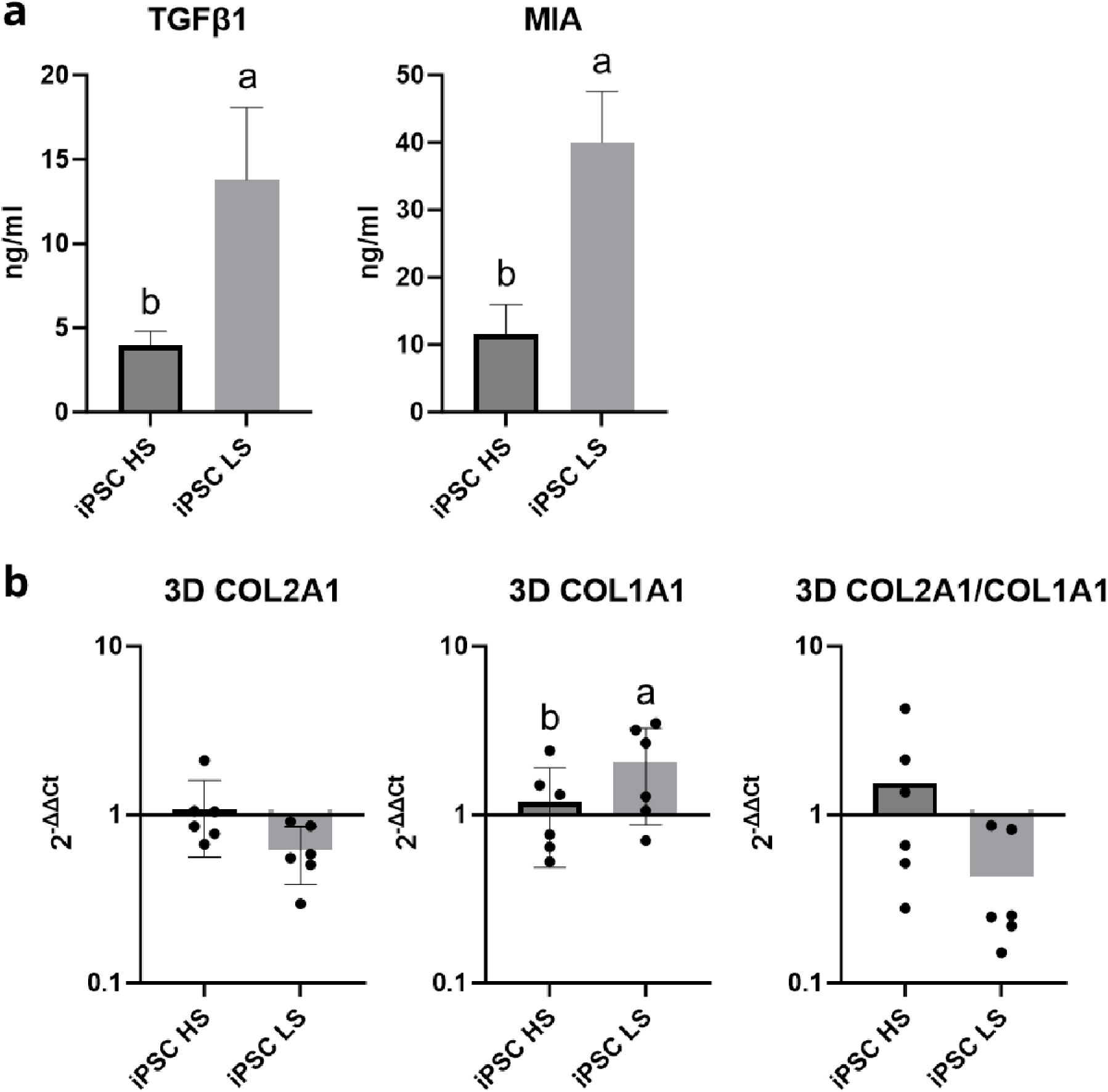
Analyses of humoral factors and chondrogenic potential of iPSC HS sheets and LS sheets. (**a**) iPSC LS sheets compared with iPSC HS sheets produced significantly higher levels of TGF-β1 and MIA. (**b**) Chondrogenic potential was evaluated through gene expression analysis after 3D culture of sheets. iPSC LS sheets exhibited significantly higher expression of *COL1A1*, but no significant differences were detected for *COL2A1* or *COL2A1* to *COL1A1* ratios. Lower-case letters of the connecting letters report show that groups not sharing the same letter are significantly different. TGF-β1, transforming growth factor beta 1; MIA, melanoma inhibitory activity.

To further probe the differences between iPSC HS and LS sheets, gene expression and cell surface marker analyses were performed. Gene expression analysis revealed that iPSC LS sheets had a significantly higher expression of *CHI3L1*, *CHI3L2*, *COL2A1*, *COL10A1*, *MIA*, *MMP13*, *MMP3*, *RUNX2*, and *SOX9* genes, and had a significantly lower expression of *DKK1* and *TGF-β1* genes. No significant differences were observed for *ACAN*, *COL1A1*, *COMP*, *ESM1*, and *GREM1* genes (Fig. 8). Comprehensive analysis of the cell surface marker profile using BD Human Lyoplate Screening Panel (BD Biosciences, Franklin Lakes, NJ, USA) (Supplementary Table 1) revealed that mesenchymal stem cell (MSC)-positive markers CD73, CD90, and CD105, were expressed at > 95% positivity by both types of sheets. MSC-negative markers CD14, CD19, CD45, and HLA-DR were expressed at < 2% positivity, while CD34 expression was slightly above 2% for iPSC HS sheets at 3.5% and iPSC LS sheets at 8.7%. Markers including CD9, CD13, CD29, CD44, CD81, CD140b, CD146 CD164, CD165, and CD166 were all expressed at > 90% positivity.

**Fig. 8.**
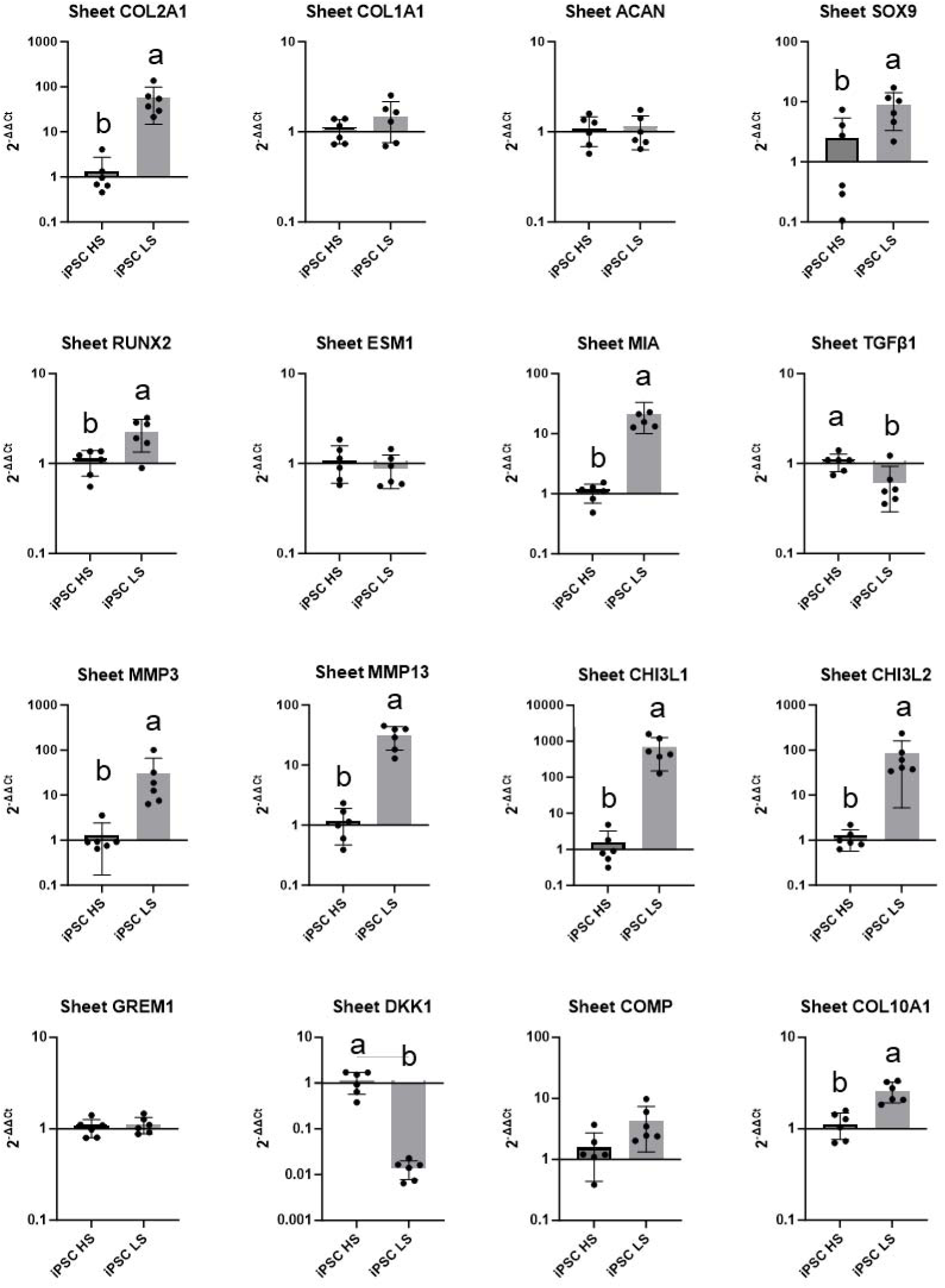
In vitro comparison between iPSC HS sheets and iPSC LS sheets. Gene expression analysis of iPSC HS sheets and LS sheets. Compared to iPSC HS sheets, iPSC LS sheets had significantly higher expression of key chondrogenic markers, such as *COL2A1*, *SOX9*, *MIA*, *CHI3L1*, and *CHI3L2*. Other hypertrophic and catabolic markers, such as *RUNX2*, *MMP13*, and *COL10A1*, were also found to be slightly higher for iPSC LS sheets compared to iPSC HS sheets. Lower-case letters of the connecting letters report show that groups not sharing the same letter are significantly different.

Single stain analysis (Table 1) was used to confirm differences in cell surface marker expression measured through both positivity and normalized median fluorescence intensity (nMFI). Notably, for iPSC LS sheets compared to iPSC HS sheets, a greater than 30% increase in positivity was observed for CD10, CD71, CD95, CD201, and CD227, while a > 30% decrease was observed for CD146 and β2-microglobulin. For others, significant increases in positivity were observed for markers such as CD9, CD26, CD56, SSEA-4, and GD2, and significant decreases for markers such as CD49a, CD49f, CD140b, and CD164. Significant decreases in nMFI were observed for CD44 and HLA-ABC.

**Table 1.**
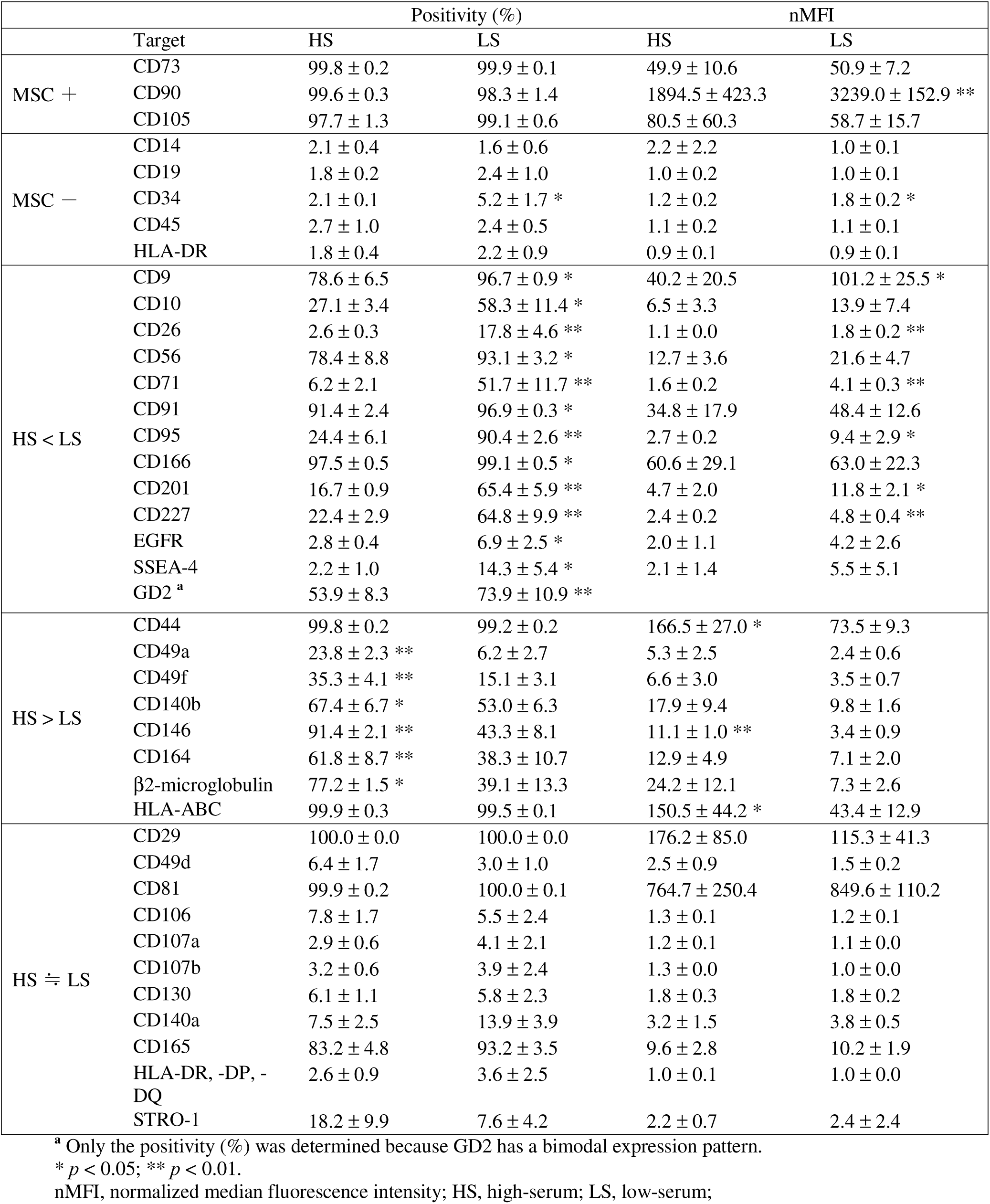
Results of flow cytometry analysis of the cells contained in iPSC sheets.

|  | Target | Positivity (%) |  | nMFI |  |
| --- | --- | --- | --- | --- | --- |
|  |  | HS | LS | HS | LS |
| MSC + | CD73 | 99.8 ± 0.2 | 99.9 ± 0.1 | 49.9 ± 10.6 | 50.9 ± 7.2 |
|  | CD90 | 99.6 ± 0.3 | 98.3 ± 1.4 | 1894.5 ± 423.3 | 3239.0 ± 152.9 ** |
|  | CD105 | 97.7 ± 1.3 | 99.1 ± 0.6 | 80.5 ± 60.3 | 58.7 ± 15.7 |
| MSC — | CD14 | 2.1 ± 0.4 | 1.6 ± 0.6 | 2.2 ± 2.2 | 1.0 ± 0.1 |
|  | CD19 | 1.8 ± 0.2 | 2.4 ± 1.0 | 1.0 ± 0.2 | 1.0 ± 0.1 |
|  | CD34 | 2.1 ± 0.1 | 5.2 ± 1.7 * | 1.2 ± 0.2 | 1.8 ± 0.2 * |
|  | CD45 | 2.7 ± 1.0 | 2.4 ± 0.5 | 1.1 ± 0.2 | 1.1 ± 0.1 |
|  | HLA-DR | 1.8 ± 0.4 | 2.2 ± 0.9 | 0.9 ± 0.1 | 0.9 ± 0.1 |
| HS < LS | CD9 | 78.6 ± 6.5 | 96.7 ± 0.9 * | 40.2 ± 20.5 | 101.2 ± 25.5 * |
|  | CD10 | 27.1 ± 3.4 | 58.3 ± 11.4 * | 6.5 ± 3.3 | 13.9 ± 7.4 |
|  | CD26 | 2.6 ± 0.3 | 17.8 ± 4.6 ** | 1.1 ± 0.0 | 1.8 ± 0.2 ** |
|  | CD56 | 78.4 ± 8.8 | 93.1 ± 3.2 * | 12.7 ± 3.6 | 21.6 ± 4.7 |
|  | CD71 | 6.2 ± 2.1 | 51.7 ± 11.7 ** | 1.6 ± 0.2 | 4.1 ± 0.3 ** |
|  | CD91 | 91.4 ± 2.4 | 96.9 ± 0.3 * | 34.8 ± 17.9 | 48.4 ± 12.6 |
|  | CD95 | 24.4 ± 6.1 | 90.4 ± 2.6 ** | 2.7 ± 0.2 | 9.4 ± 2.9 * |
|  | CD166 | 97.5 ± 0.5 | 99.1 ± 0.5 * | 60.6 ± 29.1 | 63.0 ± 22.3 |
|  | CD201 | 16.7 ± 0.9 | 65.4 ± 5.9 ** | 4.7 ± 2.0 | 11.8 ± 2.1 * |
|  | CD227 | 22.4 ± 2.9 | 64.8 ± 9.9 ** | 2.4 ± 0.2 | 4.8 ± 0.4 ** |
|  | EGFR | 2.8 ± 0.4 | 6.9 ± 2.5 * | 2.0 ± 1.1 | 4.2 ± 2.6 |
|  | SSEA-4 | 2.2 ± 1.0 | 14.3 ± 5.4 * | 2.1 ± 1.4 | 5.5 ± 5.1 |
|  | GD2 <sup>a</sup> | 53.9 ± 8.3 | 73.9 ± 10.9 ** |  |  |
| HS > LS | CD44 | 99.8 ± 0.2 | 99.2 ± 0.2 | 166.5 ± 27.0 * | 73.5 ± 9.3 |
|  | CD49a | 23.8 ± 2.3 ** | 6.2 ± 2.7 | 5.3 ± 2.5 | 2.4 ± 0.6 |
|  | CD49f | 35.3 ± 4.1 ** | 15.1 ± 3.1 | 6.6 ± 3.0 | 3.5 ± 0.7 |
|  | CD140b | 67.4 ± 6.7 * | 53.0 ± 6.3 | 17.9 ± 9.4 | 9.8 ± 1.6 |
|  | CD146 | 91.4 ± 2.1 ** | 43.3 ± 8.1 | 11.1 ± 1.0 ** | 3.4 ± 0.9 |
|  | CD164 | 61.8 ± 8.7 ** | 38.3 ± 10.7 | 12.9 ± 4.9 | 7.1 ± 2.0 |
|  | β2-microglobulin | 77.2 ± 1.5 * | 39.1 ± 13.3 | 24.2 ± 12.1 | 7.3 ± 2.6 |
|  | HLA-ABC | 99.9 ± 0.3 | 99.5 ± 0.1 | 150.5 ± 44.2 * | 43.4 ± 12.9 |
| HS ≐ LS | CD29 | 100.0 ± 0.0 | 100.0 ± 0.0 | 176.2 ± 85.0 | 115.3 ± 41.3 |
|  | CD49d | 6.4 ± 1.7 | 3.0 ± 1.0 | 2.5 ± 0.9 | 1.5 ± 0.2 |
|  | CD81 | 99.9 ± 0.2 | 100.0 ± 0.1 | 764.7 ± 250.4 | 849.6 ± 110.2 |
|  | CD106 | 7.8 ± 1.7 | 5.5 ± 2.4 | 1.3 ± 0.1 | 1.2 ± 0.1 |
|  | CD107a | 2.9 ± 0.6 | 4.1 ± 2.1 | 1.2 ± 0.1 | 1.1 ± 0.0 |
|  | CD107b | 3.2 ± 0.6 | 3.9 ± 2.4 | 1.3 ± 0.0 | 1.0 ± 0.0 |
|  | CD130 | 6.1 ± 1.1 | 5.8 ± 2.3 | 1.8 ± 0.3 | 1.8 ± 0.2 |
|  | CD140a | 7.5 ± 2.5 | 13.9 ± 3.9 | 3.2 ± 1.5 | 3.8 ± 0.5 |
|  | CD165 | 83.2 ± 4.8 | 93.2 ± 3.5 | 9.6 ± 2.8 | 10.2 ± 1.9 |
|  | HLA-DR, -DP, -DQ | 2.6 ± 0.9 | 3.6 ± 2.5 | 1.0 ± 0.1 | 1.0 ± 0.0 |
|  | STRO-1 | 18.2 ± 9.9 | 7.6 ± 4.2 | 2.2 ± 0.7 | 2.4 ± 2.4 |

<sup>a</sup> Only the positivity (%) was determined because GD2 has a bimodal expression pattern.\* $p < 0.05$ ; \*\* $p < 0.01$ .
nMFI, normalized median fluorescence intensity; HS, high-serum; LS, low-serum;

To evaluate the regenerative potential of iPSC LS sheets, the rat xenogeneic orthotopic transplantation model was used. The iPSC sheets fabricated under HS and LS conditions were transplanted, and repair tissue was evaluated at 4 weeks and 12 weeks after transplantation. No complications were observed during or following the transplantation surgery. No significant differences were detected among groups for total body weight throughout the experiment (Supplementary Fig. 29a). Weight distribution ratios were used to evaluate recovery from pain after the transplantation surgery (Supplementary Fig. 29b). Although no significant differences were detected among the groups, the HS and LS groups had improved weight distribution ratios at days 11 and 14 compared with the NT group, suggesting their faster recovery from pain.

Histological analyses were performed at 4 and 12 weeks after the transplantation. At 4 weeks, the defects of the NT group were not filled or only partially filled, while both transplantation groups showed filling of the defect with white cartilaginous tissue (Fig. 9a).

**Fig. 9.**
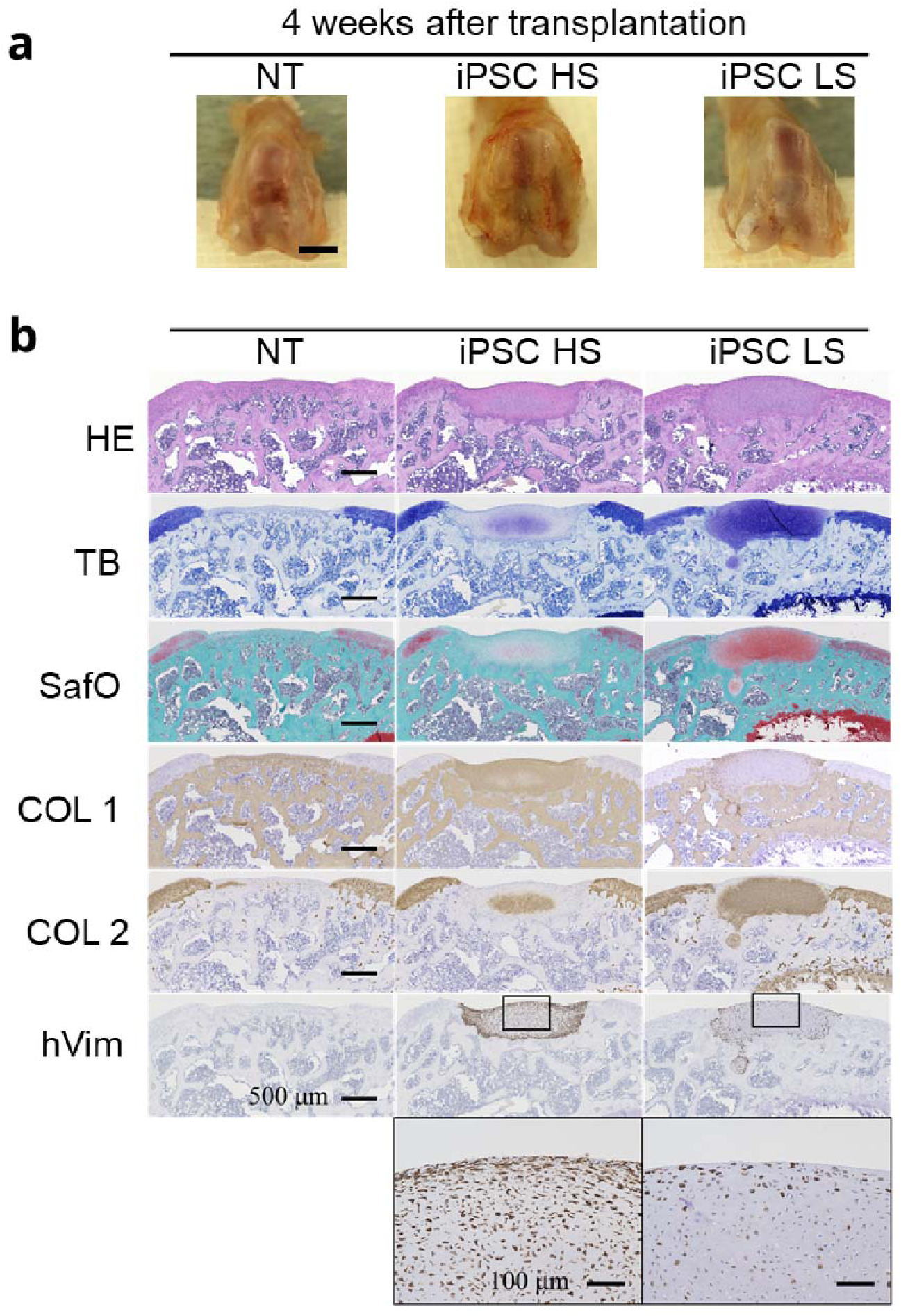
Histological analysis of regenerated cartilage in the defect areas at 4 weeks after transplantation. (**a**) Macroscopic image of the defect area and the regenerated cartilage. In the NT group, the defects were left unfilled or filled with limited amounts of white tissue. In both iPSC sheet transplantation groups, the defects were filled with white tissue. Scale bar = 1 mm. (**b**) Histological and immunohistochemical analyses of the regenerated cartilage. In the NT group, SafO staining and metachromasia with TB staining were very limited. Immunostaining of COL1 revealed strong staining within the defect area, while immunostaining of COL2 was very limited. In the iPSC HS group, slight SafO staining and slight metachromasia with TB staining were observed. Immunostaining of COL1 revealed strong staining of the repair tissue, while immunostaining of COL2 was limited. In the iPSC LS group, strong SafO staining and strong metachromasia with TB staining were observed. Immunostaining of COL1 was limited to the surface areas of the regenerated cartilage, while immunostaining of COL2 was strong for the repair tissue. Immunostaining for hVim revealed that a majority of repair tissue was formed by the transplanted cells. Vimentin staining was less dense in the iPSC LS group than in the iPSC HS group, and the cells in the iPSC LS group were observed with lacunae surrounding the cells. Scale bars = 500 μm except magnified images of hVim with scale bars = 100 μm. n = 12 per group. NT, no-treatment; HE, hematoxylin and eosin; SafO, safranin O; TB, toluidine blue; COL1, collagen type I; COL2, collagen type II; hVim, human-specific vimentin.

Histological analysis at 4 weeks revealed strong hyaline-like cartilage regeneration only in the LS group (Fig. 9b, Supplementary Fig. 17-28). In the NT group, the regeneration of cartilage was minimal with very little SafO staining or metachromasia with TB staining. The small areas of regenerated cartilage exhibited immunostaining for COL1 indicating fibrocartilage, while immunostaining for COL2 indicating hyaline-like cartilage was minimal. In the HS group, while slight staining for SafO and metachromasia with TB staining, and slight immunostaining for COL2 were observed, the majority of the tissue exhibited immunostaining for COL1. In the LS group, strong metachromasia with TB staining and strong SafO staining, and strong immunostaining for COL2 were observed, while immunostaining for COL1 was limited to the surface areas of the regenerated cartilage. Immunostaining for hVim revealed that in the transplantation groups, the defect was filled mainly with transplanted human cells. In addition, round chondrocytes with lacunae formation were observed in the LS group, whereas spindle-shaped chondrocytes without lacunae formation were observed in the HS group with larger areas of hVim immunostaining.

Similar results were observed at 12 weeks after transplantation. In the NT group, the surrounding areas of the defects had increased cartilage loss (Fig. 10a). Histological analysis at 12 weeks also revealed strong hyaline-like cartilage regeneration in the LS group with no formation of bone-like structures or significantly altered cartilage structures (Fig. 10b). Immunostaining for hVim at 12 weeks revealed comparable results to those at 4 weeks.

**Fig. 10.**
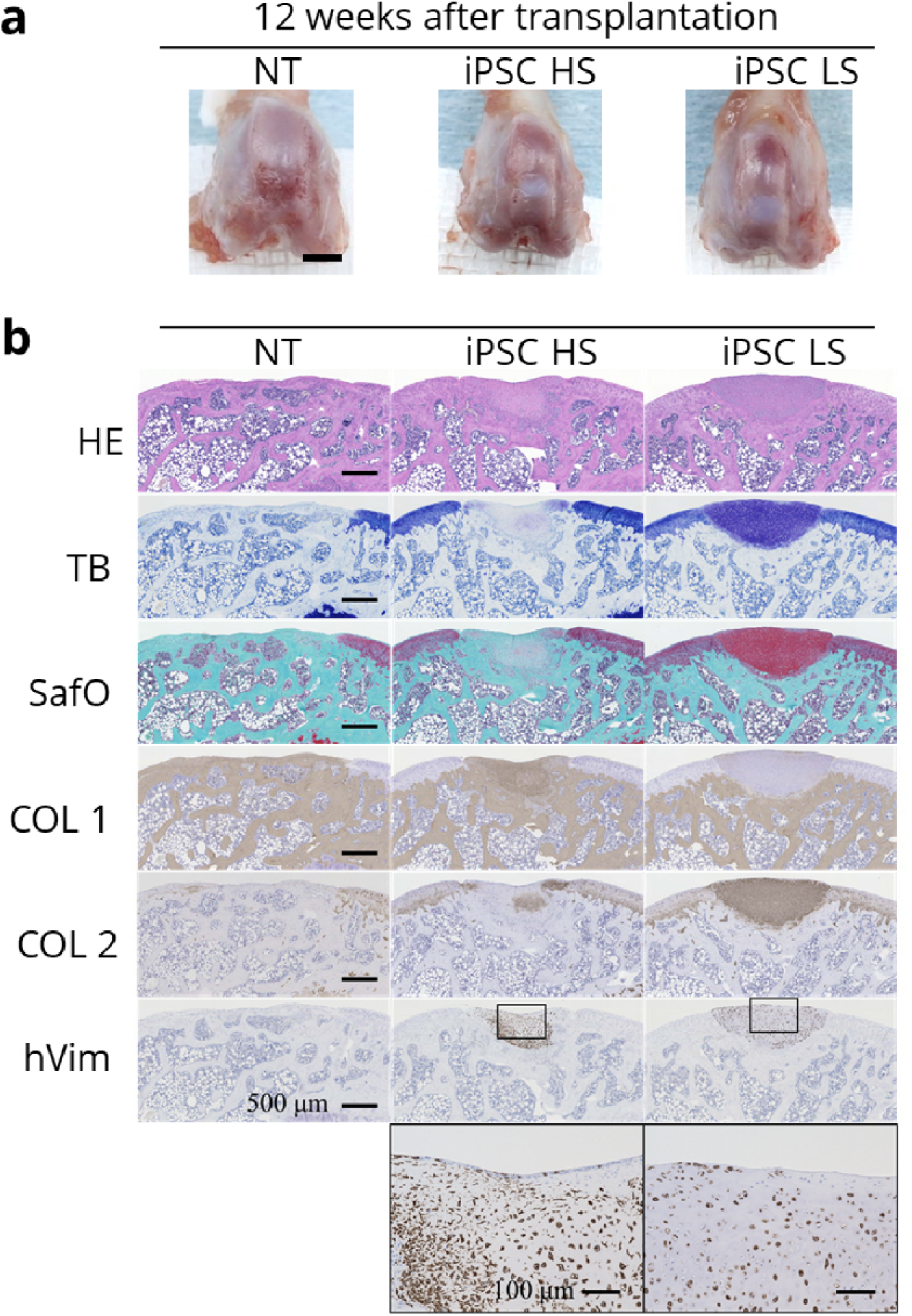
Histological analysis of regenerated cartilage in the defect areas at 12 weeks after transplantation. (**a**) Macroscopic image of the defect area and the repair tissue. In the NT group, the defects were left unfilled or filled with limited amounts of white tissue. In both iPSC sheet transplantation groups, the defects were filled with white tissue. Scale bar = 1 mm. (**b**) Histological and immunohistochemical analyses of the regenerated cartilage. In the NT group, SafO staining and metachromasia with TB staining were very limited. Immunostaining of COL1 revealed strong staining within the defect area, while immunostaining of COL2 was very limited. In the iPSC HS group, slight SafO staining and slight metachromasia with TB staining were observed. Immunostaining of COL1 revealed strong staining of the repair tissue, while immunostaining of COL2 was limited. In the iPSC LS group, strong SafO staining and strong metachromasia with TB staining were observed. Immunostaining of COL1 was limited to the surface areas of the repair tissue, and immunostaining of COL2 was strong. Immunostaining for hVim revealed that regenerated cartilage was formed mainly by the transplanted cells. Vimentin staining was less dense in the iPSC LP group than in the iPSC HS group, and the cells in the iPSC LS group were observed with lacunae surrounding the cells. Scale bars = 500 μm except magnified images of hVim with scale bars = 100 μm. n = 6 per group. NT, no-treatment; HE, hematoxylin and eosin; SafO, safranin O; TB, toluidine blue; COL1, collagen type I; COL2, collagen type II; hVim, human-specific vimentin.

Histological scores for each item are summarized in Supplementary Table 2. Compared with the NT group, the LS group had significantly higher histological total scores at both 4 weeks (*p* = 0.0187) and 12 weeks (*p* = 0.0372) (Fig. 11). Compared with the NT group, the HS group had higher total scores at both 4 and 12 weeks, although the difference was not significant. In the LS group, the greatest improvements were observed in various categories, such as basal integration and matrix staining at 4 weeks, and matrix staining and degree of defect filling at 12 weeks.

**Fig. 11.**
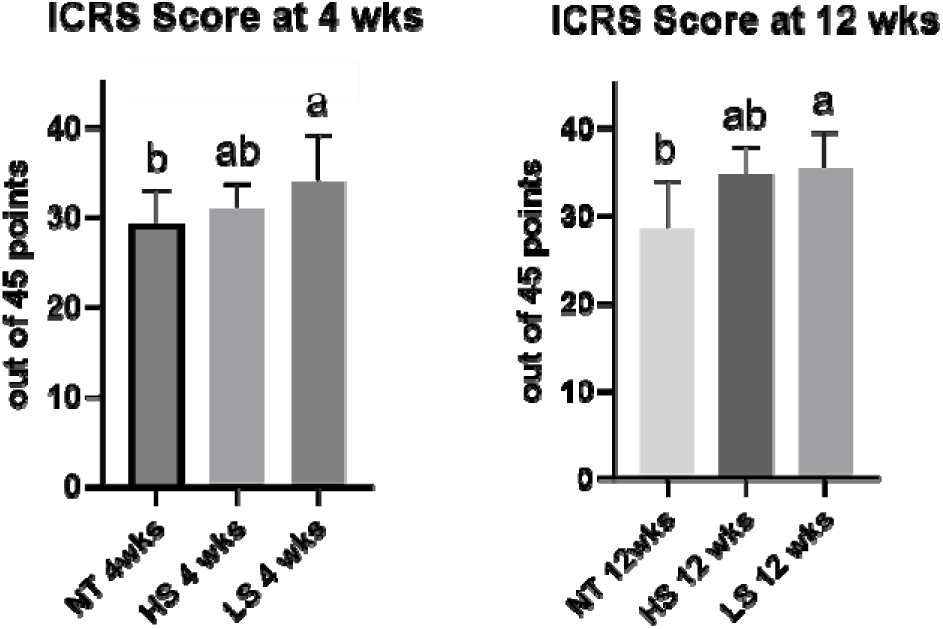
Histological scoring was performed using a modified version of the ICRS scoring system, where total higher histological scores indicate better repair cartilage quality. At both 4 weeks (n = 12 per group) and 12 weeks (n = 6 per group), the iPSC LS group had significantly higher scores compared with the no-treatment (NT) group. Lower-case letters of the connecting letters report show that groups not sharing the same letter are significantly different.

## 3. DISCUSSION

The observation that xenogeneic transplantation of iPSC LS sheets has the potential to regenerate hyaline-like cartilage in immunodeficient rats at both 4 and 12 weeks after transplantation confirms our hypothesis that a low-serum medium with hypoxic culture and high seeding density improves the *in vivo* efficacy of iPSC sheets. These results establish a proof-of-concept for iPSC sheets fabricated from clinically relevant iPS cells in hyaline cartilage regeneration.

Histological analyses of iPSC sheets *in vitro* did not particularly correlate with the potential for hyaline cartilage regeneration *in vivo*. Histological analysis confirmed that iPSC sheets across all culture conditions showed similar properties to previously reported AC and PDC sheets^9,11–13^. Further analysis showed that iPSC LS sheets contained significantly fewer cells and had reduced thickness, suggesting that chondrocytes reach confluence earlier in LS media. Despite exhibiting a dedifferentiated phenotype *in vitro*, our *in vivo* results indicate that iPSC LS sheets have the potential to regenerate hyaline-like cartilage, while iPSC HS sheets primarily produce tissue with fibrotic characteristics. The formation of lacunae by chondrocytes in the regenerated cartilage from iPSC LS sheets indicated a more mature articular cartilage phenotype. Although the specific mechanisms leading to formation of fibrocartilage versus hyaline-like cartilage remain to be investigated, the combination of LS media and hypoxic conditions likely influenced the efficacy of iPSC sheets. These findings warrant the further investigation of the mechanisms involved in the regenerative potential of these sheets.

Paracrine effects have been suggested as a key mode-of-action of chondrocyte cell sheets, warranting the investigation of secretory factors^15,32^. Interestingly, in iPSC LS sheets, the secretion levels of TGF-β1 and MIA increased by more than 3.6-fold and 3.4-fold, respectively, even with fewer numbers of cells per sheet. However, gene expression of TGF-β1 was significantly lower for iPSC LS sheets, possibly due to an autocrine effect. Both TGF-β1 and MIA are known anabolic factors of cartilage^33,34^, produced by chondrocyte cell sheets as reported previously^9,32^. Attempts to increase TGF-β1 production have been achieved in clinical trials of gene therapy using polydactyly-derived chondrocytes transduced with a retrovirus carrying TGF-β1 cDNA to treat joint inflammation in patients with OAK^35^. Higher secretion of TGF-β1, MIA, and other anabolic factors may contribute to the improved repair with hyaline-like cartilage and reduced inflammatory response.

In our search for additional *in vivo* efficacy markers, we conducted gene expression analysis and a comprehensive analysis of cell surface markers. iPSC LS sheets exhibited higher gene expression levels of factors related to cartilage formation, including *COL2A1*, *MIA*, *MMP3*, *MMP13*, and *SOX9*, suggesting that compared with iPSC HS sheets, iPSC LS sheets may be more active in remodeling of the extracellular matrix^36–38^. iPSC LS sheets also exhibited small increases in *RUNX2* and *COL10A1* expressions, but bone formation was not observed at 12 weeks *in vivo*. We found that the expression profile of iPSC sheets largely aligns with the criteria proposed by the International Society for Cellular Therapy to define human MSCs^39^, with the exception of a small CD34-positive population detected in the iPSC LS sheets. Compared with iPSC HS sheets, iPSC LS sheets exhibited a significantly higher expression of cell surface markers including CD9, CD10, CD26, CD201, CD227, and SSEA-4, which are also highly expressed (> 80%) in PDC sheets. Notably, HLA-DR, -DP, and -DQ were not expressed by iPSC sheets, suggesting low immunogenicity^12^. Furthermore, iPSC LS sheets showed significantly higher expression levels of CD56 and GD2, which have low expression levels (< 20%) in PDC sheets. CD56, a cell surface glycoprotein important for cell-cell adhesion, has been used to isolate distinct MSC populations from human bone marrow^40,41^. Studies have indicated that deficiency of CD56 expression by MSCs accelerates chondrocyte hypertrophy in an experimental OA model^42^. GD2 has been identified as a potential marker for MSCs isolated from human bone marrow^43^ and has also been used to isolate progenitor cell populations in the nucleus pulposus^44,45^. While gene expression and cell surface markers may not be directly linked to chondrogenic potential, they can serve as valuable *in vivo* efficacy markers and will be investigated in future studies. Additional studies comparing iPSC sheets with PDC sheets as well as AC sheets will provide greater insight into potential efficacy markers for the regenerative potential of chondrocyte cell sheets.

Several limitations exist in this study that warrant further investigations. One limitation is that only one donor of iPS cells was used. This approach was taken because iPS cells from donor QHJI are one of the best-studied cell lines and have been used in clinical studies to treat exudative age-related macular degeneration and Parkinson’s disease^46,47^. Further studies with other cell lines are needed to expand the range of iPS cells that can be used. Another limitation is that the exact contents of the LS medium are proprietary, including the origin of the serum. The possibility that supplements to the LS medium may contribute to the regenerative effects of iPSC LS sheets remains to be investigated.

While different jurisdictions impose different sets of criteria towards clinical application, additional studies are necessary to establish safety and potential efficacy of iPSC sheets. The rat xenogeneic transplantation model, a cost-effective model in optimizing the fabrication of human iPSC sheets, has limitations in the translational pathway. One limitation is that the osteochondral defect model is an acute model that does not completely reflect the chronic inflammation and catabolic environment of OAK that affect the entire joint, including both articular cartilage and synovial tissue. Another limitation is that the athymic rat is immunocompromised and does not reflect an allogeneic response. In assessing the dynamic changes during the regenerative response, regenerated tissue and the synovial pathology should be further characterized for biomechanical, biochemical, tribological properties and for inflammatory responses, such as macrophage phenotypes, at different time points to better understand the repair process. In addition, long-term studies in large animal models are necessary in most jurisdictions to obtain approval for clinical use^48,49^, warranting future studies that use an analogous cellular product for preclinical testing^49,50^. Further comparative studies of iPSC sheets and PDC sheets may also be needed to support the rationale for clinical translation.

Safety is of utmost importance in cell-based therapies using iPS cells. Our current approach, utilizing both iPS cells and iPS-Carts that are well characterized for safety, accelerates this translational process, although the final human product, i.e., iPSC sheets, will need to undergo additional testing such as immunogenicity, tumorigenicity, toxicity, and purity testing^49^. It is nevertheless encouraging that iPS cells from the iPS Cell Stock Project that are used in this study are well characterized and tested for safety^19,51^. Other preclinical studies have demonstrated the safety of cell-based therapies derived from donor QHJI^52^. Additionally, QHJI01s04 used in this study is one of two iPS cell lines that have registered Drug Master Files with the US FDA, aiding commercial development^53^. Also, iPS-Carts themselves have been shown to have low immunogenicity^54^ and low tumorigenicity^52^, while concerns for cells obtained from iPS-Carts to form bone^55–57^ and off-target populations^58,59^ have been raised. Additional evidence for safety is expected from an ongoing clinical study using iPS-Carts to treat knee cartilage defects (jRCTa050190104). Future safety studies for iPSC sheets, supported by safety evidence of iPS cells and iPS-Carts used, will support first-in-human studies using iPSC sheets.

## 4. CONCLUSIONS

In this study, we established proof-of-concept for iPSC sheets fabricated from clinically relevant iPS cells through a xenogeneic transplantation model using immunodeficient rats. iPSC sheets fabricated using a reduced-serum medium, hypoxic culture, and high seeding densities demonstrated their potential in regenerating hyaline-like cartilage *in vivo*. In the future, additional safety and large animal studies will be conducted to lay a foundation for the clinical application of iPSC sheets.

## 5. METHODS

The use of iPS cells from the iPS Cell Stock Project and their derivatives was approved by the Kyoto University Ethics Committee (Clinical #79) and Tokai University Medical Ethics Committee (18I-#46).

Animal studies were approved by the Institutional Animal Experiment Committee at Tokai University (#172046, #195015, #201034, #212037) and were performed in accordance with the guidelines of the Institutional Regulations for Animal Experiments and the Fundamental Guidelines for Proper Conduct of Animal Experiments and Related Activities in Academic Research Institutions under the jurisdiction of the Ministry of Education, Culture, Sports, Science, and Technology for animal handling and care.

### 5.1 Fabrication of iPS cell-derived chondrocyte sheets

Clinical-grade HLA homozygous iPS cells established from human adult PBMCs (QHJI 01s04) were used. The HLA haplotype of donor QHJI covers approximately 17% of the Japanese population and is the most common in Japan^18^. iPS cells were generated in a feeder-free, xeno-free culture system, and reprogramming genes were introduced by the electroporation of episomal plasmid vectors (pCE-OCT3/4, pCE-hSK, pCE-hUL, pCE-mp53DD and PCXB-EBNA1)^19,60^. Extensive testing for safety has been performed, and test results are publicly available^61^. For example, plasmid remnants were evaluated through qPCR and reported as below the limit of quantification. iPS-Carts were fabricated at CiRA through methods reported previously^23^ and six lots of iPS-Carts were used throughout the study. Briefly, iPS cells were differentiated toward the mesoendoderm lineage, followed by chondrogenic differentiation. The cells were transferred to 3D culture after 2 weeks and cultured in chondrogenic media for another 14 weeks. The iPS-Carts were then shipped to Tokai University in a specialized container with heat packs (Sanplatec Corp., Osaka, Japan) at 37 °C. The shipped iPS-Carts were placed in a humidified incubator at 37 °C in 5% CO_2_ and cultured for an additional 3–5 weeks prior to experiments. The chondrogenic medium was replaced every 3 to 4 days.

To fabricate iPSC sheets, iPS-Carts were digested in Dulbecco’s modified Eagle’s medium: F12 nutrient mixture (DMEM/F12; Thermo Fisher, Waltham, MA, USA) supplemented with 20% fetal bovine serum (FBS; SAFC Biosciences, Lenexa, KS, USA), 1% antibiotic, antimycotic solution (AB; Fujifilm, Osaka, Japan), and 167 μg/mL Liberase MNP-S GMP Grade (Roche, Basel, Switzerland) for 2.5–3.0 h at 37 °C in 5% CO_2_. The cell suspension was filtered through a 40-μm cell strainer (Greiner, Kremsmünster, Austria) and washed three times with DMEM/F12 medium supplemented with 20% FBS and 1% AB. The collected cells were directly seeded to temperature-responsive culture inserts (CellSeed, Tokyo, Japan) at 5 × 10^4^ cells/cm^2^ and subsequently cultured in a humidified incubator (Taitec, Koshigaya, Saitama, Japan) at 37 °C in 5% CO_2_ under 2% O_2_ (i.e., hypoxic) conditions. DMEM/F12 supplemented with 20% FBS and 1% AB (HS medium) was used to seed iPSC HS sheets. After day 3, 100 μg/mL ascorbic acid (AA; Wako Pure Chemical Industries, Osaka, Japan) was added to the HS medium and the medium was replaced every 3 days. Similarly, MesenPRO medium (Thermo Fisher), a proprietary formula containing 2% serum, was supplemented with 2 mM GlutaMAX Supplement (Thermo Fisher) and 1% AB (LS medium) to seed iPSC LS sheets. AA was not added to LS medium in this study. LS medium was replaced every other day according to the manufacturer’s instructions, as replacing the medium every 3 days significantly decreased proliferation in a preliminary experiment.

### 5.2 Cell count and viability of cell sheets

The iPSC sheets were washed in Dulbecco’s phosphate-buffered saline (DPBS; Thermo Fisher) and incubated in 2 mL accutase (BD Biosciences) at room temperature (RT) for 15 min. The sheets were collected and resuspended in HS medium containing 0.25 mg/mL Collagenase P (Roche) at 37 °C for 30 min. The collected cells were resuspended in HS medium, and trypan blue exclusion assay was performed to determine cell count and viability using a Countess II Automated Cell Counter (Thermo Fisher). The experiments were repeated with five lots (technical n = 2 or 3).

### 5.3 Histological and immunohistochemical analyses of cell sheets

Histological evaluation of cell sheets was performed using HE, TB, and SafO stains, and immunohistochemical evaluation was performed for COL1, COL2, ACAN, and FN using methods reported previously^13^. The primary and secondary antibodies used are listed in Supplementary Table 3. All images were obtained using a BZ-X810 fluorescence microscope (Keyence, Osaka, Japan).

### 5.4 Measurement of humoral factors in culture supernatants

After 2 weeks of culture, HS and LS media were replaced by DMEM/F12 medium supplemented with 1% FBS and 1% AB, and cultured in a humidified, normoxic incubator for 72 h at 37 °C in 5% CO_2_. A total of 2 mL of culture supernatants inside the temperature-responsive culture inserts was collected, centrifuged at 15,000 *g* for 10 min, and stored at –80 °C. Commercially available enzyme-linked immunosorbent assay kits were used to measure the concentrations of TGF-β1 (R&D Systems, Minneapolis, MN, USA) and MIA (Roche). Absorbance measurements were made using a Multiskan GO spectrophotometer (Thermo Fisher). The experiments were repeated with four lots (technical n = 2).

### 5.5 Comprehensive analysis of cell surface markers

The Human Cell Surface Marker Screening Panel (BD Biosciences), consisting of 242 monoclonal antibodies to cell surface markers, was used to identify differences between the two types of iPSC sheets. The experiment was conducted from a single lot of iPSC-Carts as a preliminary step. Briefly, 20 iPSC sheets per group were washed in DPBS, incubated in accutase for 5 min at RT, and then pooled in 50 mL tubes (TPP, Trasadingen, Switzerland). The sheets were suspended in 0.25 mg/mL of Collagenase P dissolved in HS culture medium for 30 min in a 37 °C water bath. The isolated cells were washed with DPBS and then filtered through 40-μm cell strainers. The cells were then suspended in wash buffer composed of DPBS with 0.2% FBS and 1 mM EDTA (Thermo Fisher). Cell counting was performed, and total suspension volume was adjusted to 1.0 × 10^5^ cells per 100 μL. Microtiter 96-well plates were prepared according to the manufacturer’s protocol with 20 μL of antibody and 100 μL of the cell suspension per well. The plates were incubated for 30 min on ice, and then the cells were washed. Next, the cells were suspended in corresponding secondary antibodies for 20 min on ice in the dark. The cells were washed and then incubated in 100 μL of Cytofix Fixation Buffer (BD Biosciences) for 15 min on ice. The cells were washed and then analyzed using a FACSVerse analyzer (BD Biosciences). At least 10,000 events were observed per sample. Flow cytometry data were analyzed using FlowJo Software (version 10.7.1; BD Biosciences). The gating strategy is illustrated in Supplementary Fig. 30. To determine the positivity of cells, gates were set for corresponding isotype controls to < 2% positivity. The nMFI was determined as the fold change between the MFIs of the target and the corresponding isotype control^62^.

### 5.6 Single stain analysis of selected cell surface markers

For subsequent analysis, a single stain protocol was used to evaluate selected cell surface markers with the addition of STRO-1, a frequently used MSC marker (Supplementary Table 4). The cells were isolated from iPSC sheets and analyzed using the same methods as the comprehensive analysis without the fixation step. After incubating the cells with antibodies at 4 °C for 30 min, the cells were washed and immediately analyzed with FACSVerse analyzer. The experiments were repeated with four lots.

### 5.7 Gene expression analysis

For gene expression analysis, fabricated iPSC sheets were each placed in 1 mL TRIzol reagent (Thermo Fisher) and dissociated using beads in a 2.0 mL hard tube (Bio Medical Science, Tokyo, Japan) for 3 min at 1500 rpm using a Shake Master NEO (Bio Medical Science). Total RNA was obtained using an RNeasy Mini Kit (Qiagen, Hilden, Germany) with the RNase-Free DNAse Set (Qiagen), according to the manufacturer’s instructions. RNA was converted into cDNA using the QuantiTect Reverse Transcription Kit (Qiagen), according to the manufacturer’s instructions. For real-time polymerase chain reaction (RT-PCR), 2.5 ng of cDNA was mixed with RNase-free water (Qiagen) in a total of 9.6 μL along with 0.2 μL of forward primer (final 500 nM; Eurofins Genomics K.K., Tokyo, Japan), 0.2 μL of reverse primer (final 500 nM), and 10.0 μL of PowerUp Syber Green Master Mix (Thermo Fisher) for a final volume of 20 μL. RT-PCR was performed using the QuantStudio 3 Real-Time PCR System (Thermo Fisher), according to the manufacturer’s instructions. Briefly, the reaction mixture was incubated at 50 °C for 2 min., 95 °C for 2 min., followed by 40 cycles of 95 °C for 15 s and 60 °C for 1 min. Dissociation curves were obtained by incubating the PCR product at 95 °C for 15 s, 60 °C for 1 min., and 95 °C for 15 s, to verify a single product. The –delta Ct was calculated using RPL13A as the housekeeping gene to compare the relative gene expression levels of iPSC sheets. A list of primers used for RT-PCR is provided in Supplementary Table 5. The experiments were repeated with six lots (technical n = 2 or 3).

### 5.8 Evaluation of chondrogenic potential

Chondrogenic potential was evaluated using a 3D culture method developed for chondrocyte cell sheets. Fabricated iPSC sheets were transferred to HydroCell 6-well dishes (CellSeed), a super-low cell-binding dish, and cultured in HS medium supplemented with 100 μg/mL AA, 0.35 mM L-proline (Sigma), 0.1 μM dexamethasone (Sigma), 1:500 insulin-transferrin-selenium-A (Thermo Fisher), and 10 ng/mL TGF-β1 (Peprotech, Cranbury, NJ, USA) for 7 days in a humidified, normoxic incubator at 37 °C in 5% CO_2_. The spontaneously formed 3D pellets were collected for gene expression analysis. Using the same protocol as for evaluating iPSC sheets, the gene expression levels of *COL2A1* and *COL1A1* were measured to estimate the ability of iPSC sheets to form hyaline-like cartilage tissue. The experiments were repeated with six lots and two or three technical replicates.

### 5.9 Evaluation through xenogeneic transplantation

A total of 72 and 52 in Phases 1 and 2, respectively, of 12-week-old male athymic nude rats (F334/Njcl-rnu/rnu; Clea Japan, Tokyo, Japan), weighing approximately 270 g, were used in the study. Two rats were housed per cage with access to water and food *ad libitum*. Prior to surgery, the rats were randomized by total body weight and allocated to three groups (no treatment [NT group], iPSC HS sheet transplantation [HS group], or iPSC LS sheet transplantation [LS group]), and evaluated at either 4 weeks (n = 12 per group) or 12 weeks (n = 6 per group).

Transplantation surgery was performed as reported previously^12^. An osteochondral defect (2 mm diameter, 1 mm depth) was created, and in the transplantation groups, one-half of the 4.2 cm^2^ cell sheet was transplanted to each defect so as to cover the entire defect and the surrounding areas. The rats were free to move around following surgery.

### 5.10 Evaluation of pain and discomfort posttransplantation

Total body weight was monitored posttransplantation and an incapacitance tester (Linton Instrumentation, Diss, Norfolk, England) was used to evaluate pain and discomfort, as reported previously^11^. Measurements were made on days 3, 7, 11, 14, 17, 21, 24, and 28 after transplantation. The average surgical limb weight distribution ratio (%) of the hind limbs was calculated from 10 repeated measurements of each animal as follows: operated limb weight distribution ratio (%) = operated limb load (g) / total limb load (g) × 100.

### 5.11 Histological evaluation of regenerated cartilage

Rats were euthanized humanely by administration of high-dose Nembutal at 4 and 12 weeks after transplantation. The operated knee was opened, and the distal portion of the femur was excised, processed, and embedded in paraffin wax, and then standard protocols were used for histological and immunohistochemical staining, as reported previously^11^. Hematoxylin and eosin (HE), TB, and SafO/Fast Green staining was performed to evaluate the cartilage filling the defect. Immunohistochemical staining was performed for COL1 and COL2.

The human-specific vimentin (hVim) immunostaining was modified slightly. Briefly, deparaffinized sections were treated with 0.01 M citric acid buffer for 10 min at 98 °C. The sections were washed in 0.01 M phosphate-buffered saline, blocked with normal equine serum (Vector Laboratories, Burlingame, CA, USA), and then treated with a hVim antibody overnight at 4 °C. The stained sections were washed and treated with ImmPRESS polymer anti-rabbit IgG reagent (Vector Laboratories) for 1 h at RT, and then washed again and immersed for 2 min in Tris-HCl buffer (pH 7.6) containing 0.05% diaminobenzidine and 0.0054% hydrogen peroxide, and then counterstained with HE. Primary antibodies used are summarized in Supplementary Table 3. All images were obtained using a BZ-X810 fluorescence microscope (Keyence).

### 5.12 Histological scoring of regenerated cartilage

A modified version of the International Cartilage Regeneration and Joint Preservation Society (ICRS) histological grading system was used to evaluate the regenerated cartilage, as reported previously^11,13,63^. The images obtained from SafO staining were evaluated by two trained scorers (H.O. and S.W.), who were blinded to the group identities. A description of the scoring items and breakdown of scores are provided in Supplementary Table 6.

### 5.13 Statistical analysis

All results are expressed as average and standard deviation. Prism 10 (GraphPad) was used to perform statistical analysis and generate plots. A paired *t* test was used to determine statistical significance between two groups. Each experiment was performed from a single lot of iPS-Cart to compare the two iPSC sheets. One-way analysis of variance (ANOVA) with Tukey’s test for *post hoc* analysis was used to identify significant differences among three or more groups. For analysis of FACS data, when cell positivity included extreme values close to 100%, only the nMFI values were used to determine statistical differences. ICRS histological scores among the groups at 4 and 12 weeks were evaluated using one-way ANOVA with Tukey’s test for *post hoc* analysis to identify significant differences between groups. For all analyses, *p* values < 0.05 were considered statistically significant. Connecting letters report was used to indicate significant differences, in which different letters indicate that a significant difference exists between groups.

## AUTHOR CONTRIBUTIONS

T.T.: Conception and design, financial support, collection and/or assembly of data, data analysis and interpretation, manuscript writing. U.R.: Collection and/or assembly of data, data analysis and interpretation. E.T.: Conception and design, financial support, data analysis and interpretation. M.M.: Collection and/or assembly of data, data analysis and interpretation. Y.T.: Collection and/or assembly of data, data analysis and interpretation. S.W.: Collection and/or assembly of data. H.O.: Collection and/or assembly of data. M.O.: Collection and/or assembly of data. A.Y.: Provision of study material or patients. M.W.: Administrative support. N.T.: Provision of study material or patients, financial support. M.S.: Conception and design, financial support, data analysis and interpretation, and final approval of manuscript.

## ACKNOWLEDGEMENTS

This research was funded by Japan Agency for Medical Research and Development (AMED) under grant numbers JP17bm0104001, JP17bm0304004 (to NT), and JP20bm0404035 (to MS), by Japan Society for the Promotion of Science (JSPS) KAKENHI grant numbers JP16K01396 (to ET) and JP19K20680 (to TT), and by 2017, 2018, 2019, and 2020 Tokai University School of Medicine Research Aids (to TT). Funders played no role in study design, data collection, analysis and interpretation of data, or the writing of this manuscript.

The authors remember Miho Morioka for her invaluable contributions that guided this work. The authors are grateful for Nobuyuki Shima’s critical experimental advice. The authors are grateful for Eri Okada, Saori Shirasuna, and Satoko Suzuki’s assistance with data collection. The authors are grateful for Kumiko Kitamura and Ayako Watanabe’s administrative assistance in managing grants. The authors are grateful to the Medical Science College Office, Tokai University, for their technical support in animal studies, histological analysis, and flow cytometric analysis.

## Data availability statement

The data obtained in this study are available upon reasonable request to the corresponding author (MS).

## ABBREVIATIONS

AC: autologous chondrocyte
ACAN: aggrecan
CiRA: Center for iPS Cell Research and Application
COL1: collagen type I
COL2: collagen type II
FN: fibronectin
HE: hematoxylin and eosin
HLA: human leukocyte antigen
HS: high-serum
hVim: human-specific vimentin
ICRS: International Cartilage Regeneration and Joint Preservation Society
iPS cell: induced pluripotent stem cell
iPSC: iPS cell-derived chondrocyte
iPS-Cart: iPS cell-derived cartilaginous tissue
LS: low-serum
MIA: melanoma inhibitory activity
MSC: mesenchymal stem cell
nMFI: normalized median fluorescence intensity
NT: no-treatment
OAK: osteoarthritis of the knee
PBMC: peripheral blood mononuclear cell
PDC: polydactyly-derived chondrocyte
PVDF: polyvinylidene difluoride
SafO: safranin O
TB: toluidine blue
TGF-β1: transforming growth factor beta 1

