## Supplementary Figures for "Enhancing the hyaline cartilage regenerative potential of induced pluripotent stem cell-derived chondrocyte cell sheets"

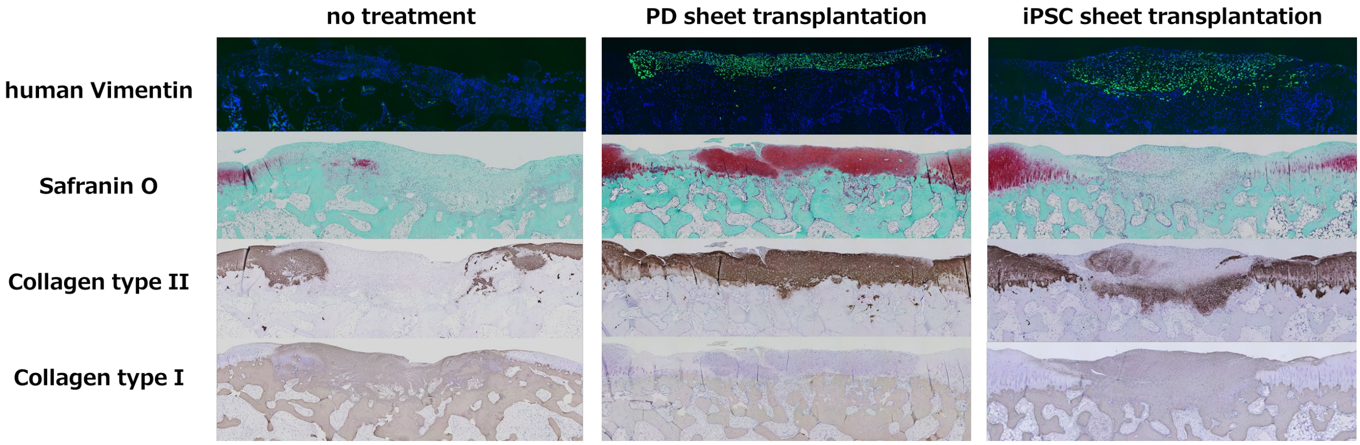


**Supplementary Figure 1.** In a preliminary study, a rabbit xenogeneic transplantation model was used to compare the *in vivo* efficacy of iPS cell-derived chondrocyte cell sheets (iPSC sheet) fabricated under the same conditions as polydactyly-derived chondrocyte cell sheets (PD sheets). Only the PD sheet transplantation group exhibited regeneration of hyaline cartilage as indicated by immunostaining of type II collagen. In the iPSC sheet transplantation group, the regenerated cartilage was positive for immunostaining of type I collagen indicating that the regenerated cartilage was mostly fibrocartilage. Immunostaining of human-specific vimentin revealed that the regenerated cartilage was composed of the transplanted human cells, confirming successful engraftment of transplanted cells in both groups.

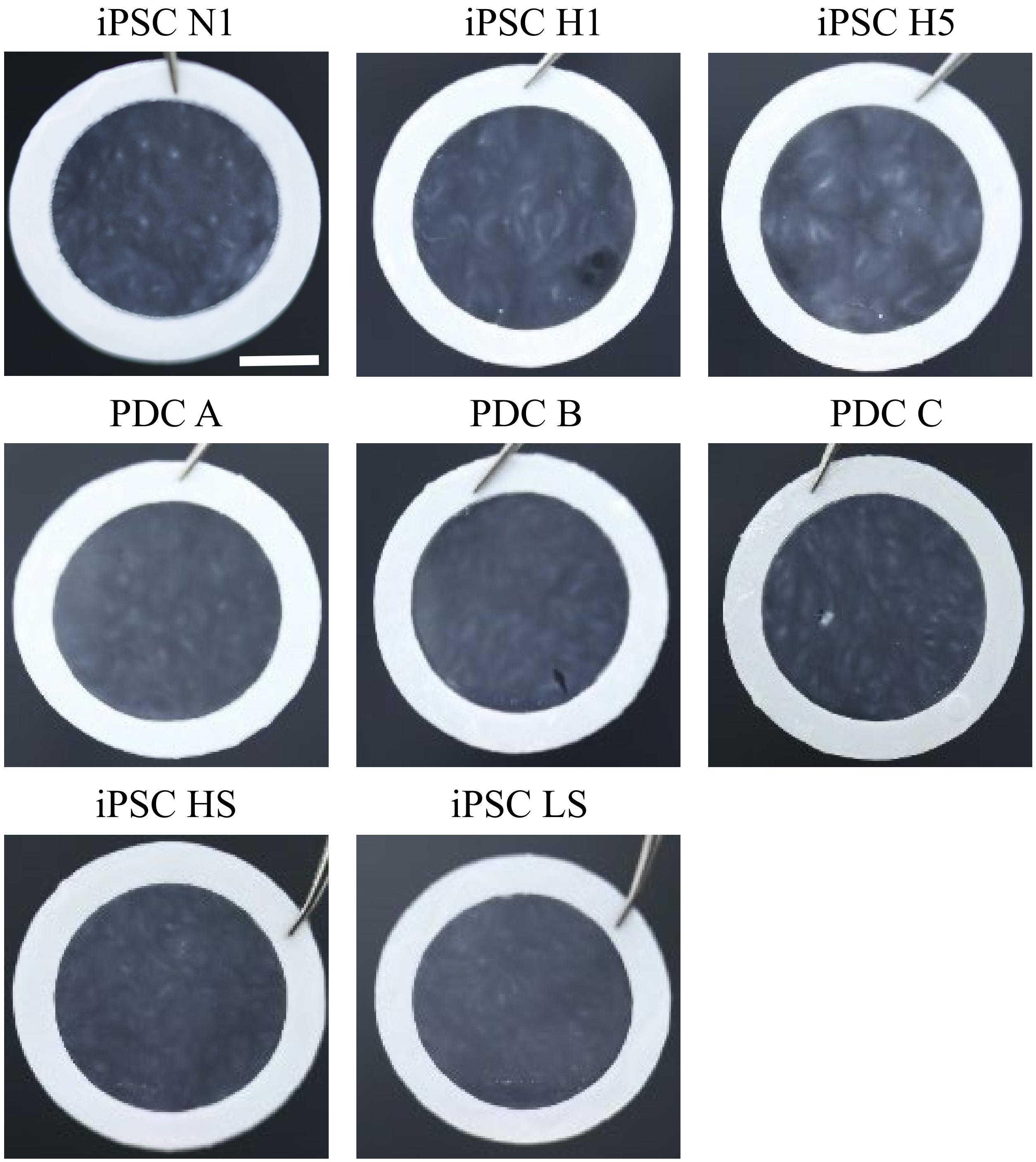


**Supplementary Figure 2. Macroscopic images of iPSC sheets and PDC sheets**. Both iPSC sheets and PDC sheets exhibit a thin sheet structure that can be picked up using PVDF membrane. Scale bar = 5 mm.

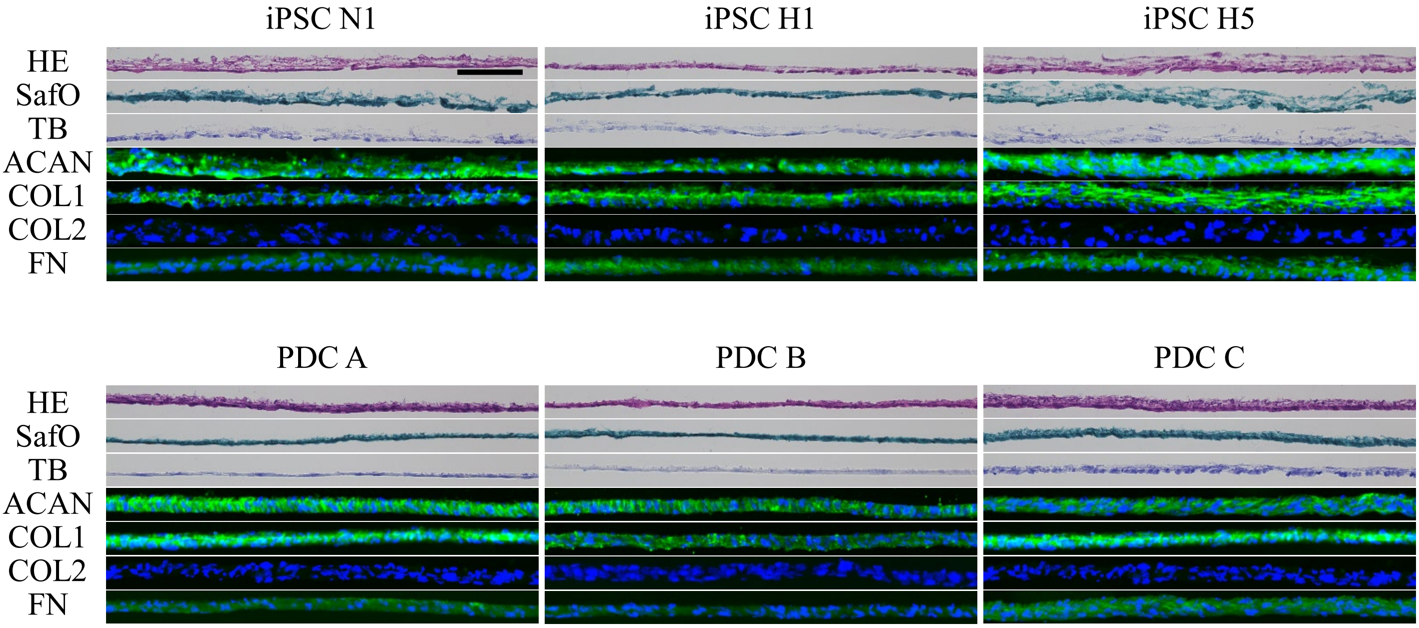


**Supplementary Figure 3. Histological analysis of iPSC sheets and PDC sheets**. Both iPSC sheets and PDC sheets express ACAN, COL1, and FN but not COL2, indicating a dedifferentiated chondrocyte phenotype. Scale bar = 100 μm.

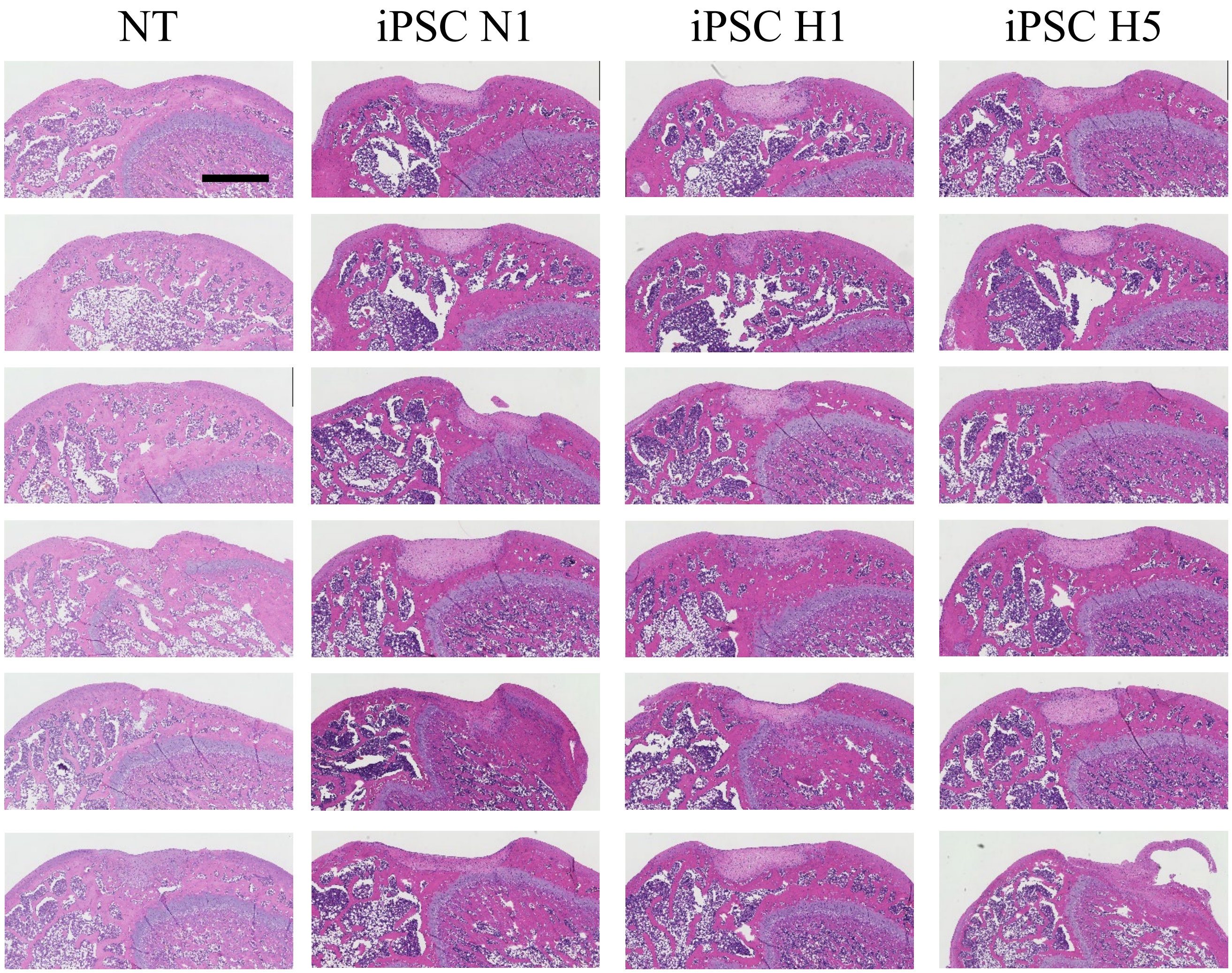


**Supplementary Figure 4. Histological evaluation at 4 weeks after transplantation of iPSC sheets in a xenogeneic orthotopic transplantation model using athymic nude rats.** Hematoxylin and eosin staining results for all animals. n = 6 per group. Scale bar = 500 μm.

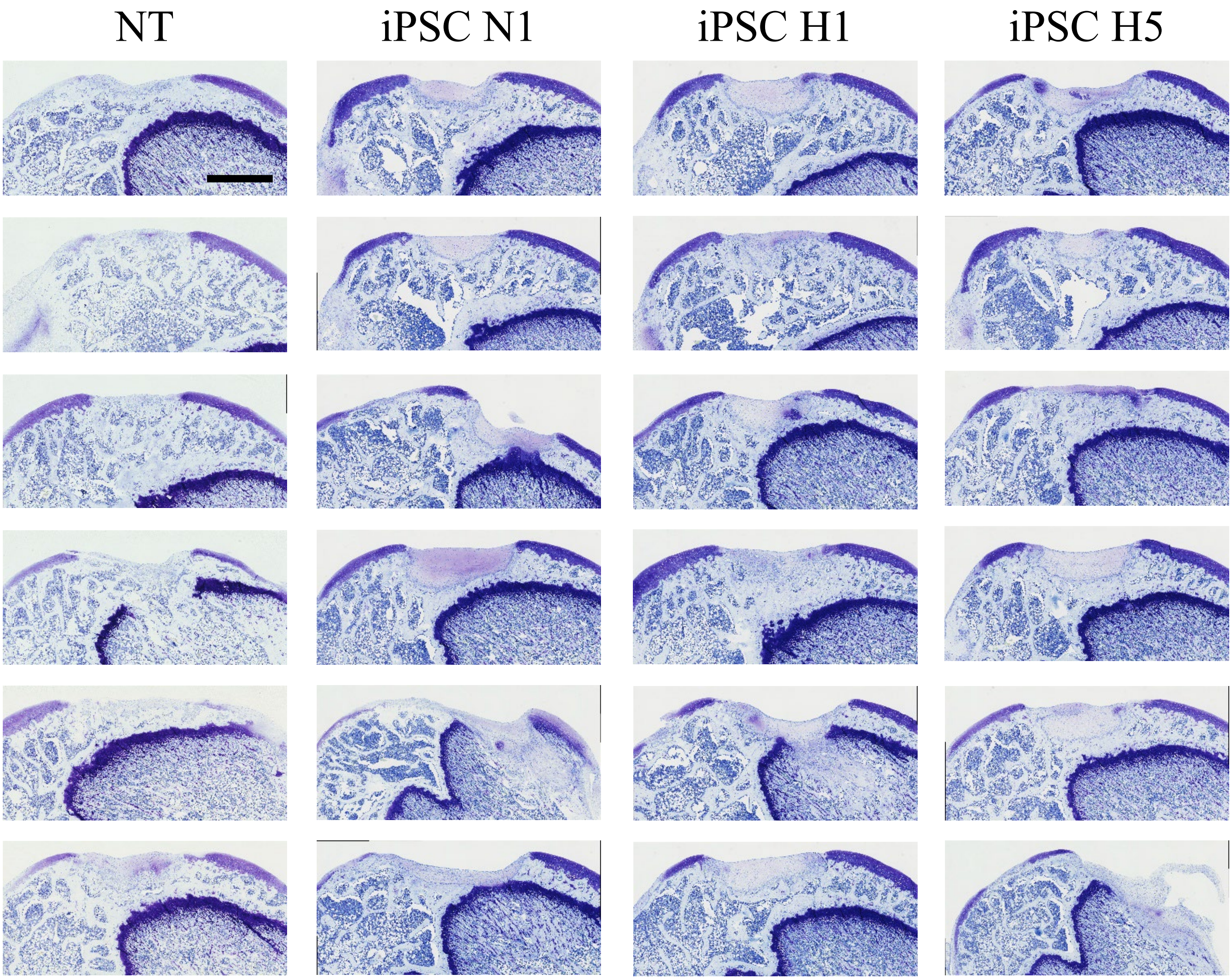


**Supplementary Figure 5. Histological evaluation at 4 weeks after transplantation of iPSC sheets in a xenogeneic orthotopic transplantation model using athymic nude rats.** Toluidine blue staining results for all animals. n = 6 per group. Scale bar = 500 μm.

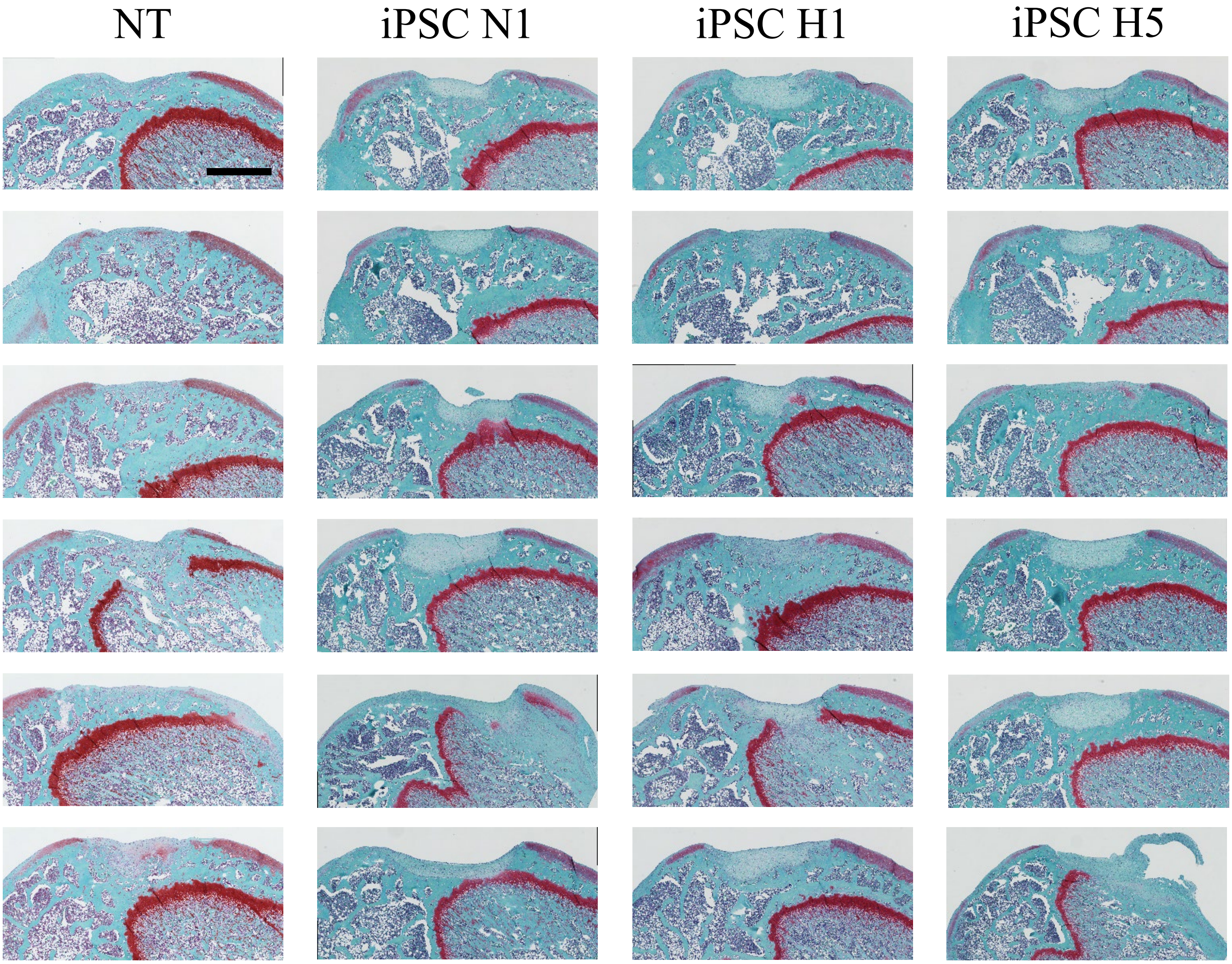


**Supplementary Figure 6. Histological evaluation at 4 weeks after transplantation of iPSC sheets in a xenogeneic orthotopic transplantation model using athymic nude rats.** Safranin O staining results for all animals. n = 6 per group. Scale bar = 500 µm.

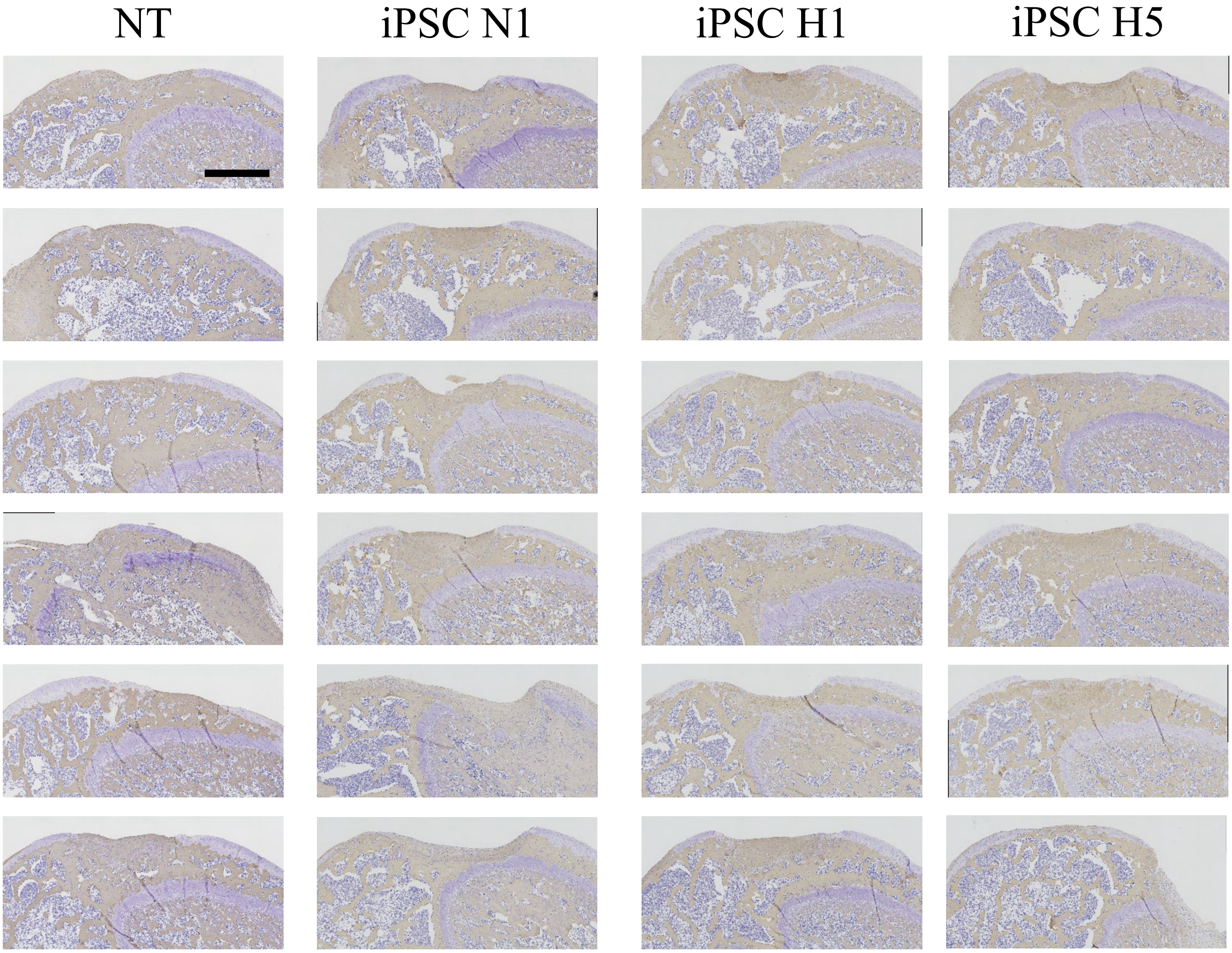


**Supplementary Figure 7. Histological evaluation at 4 weeks after transplantation of iPSC sheets in a xenogeneic orthotopic transplantation model using athymic nude rats.** Immunohistological staining for collagen type I for all animals. n = 6 per group. Scale bar = 500 µm.

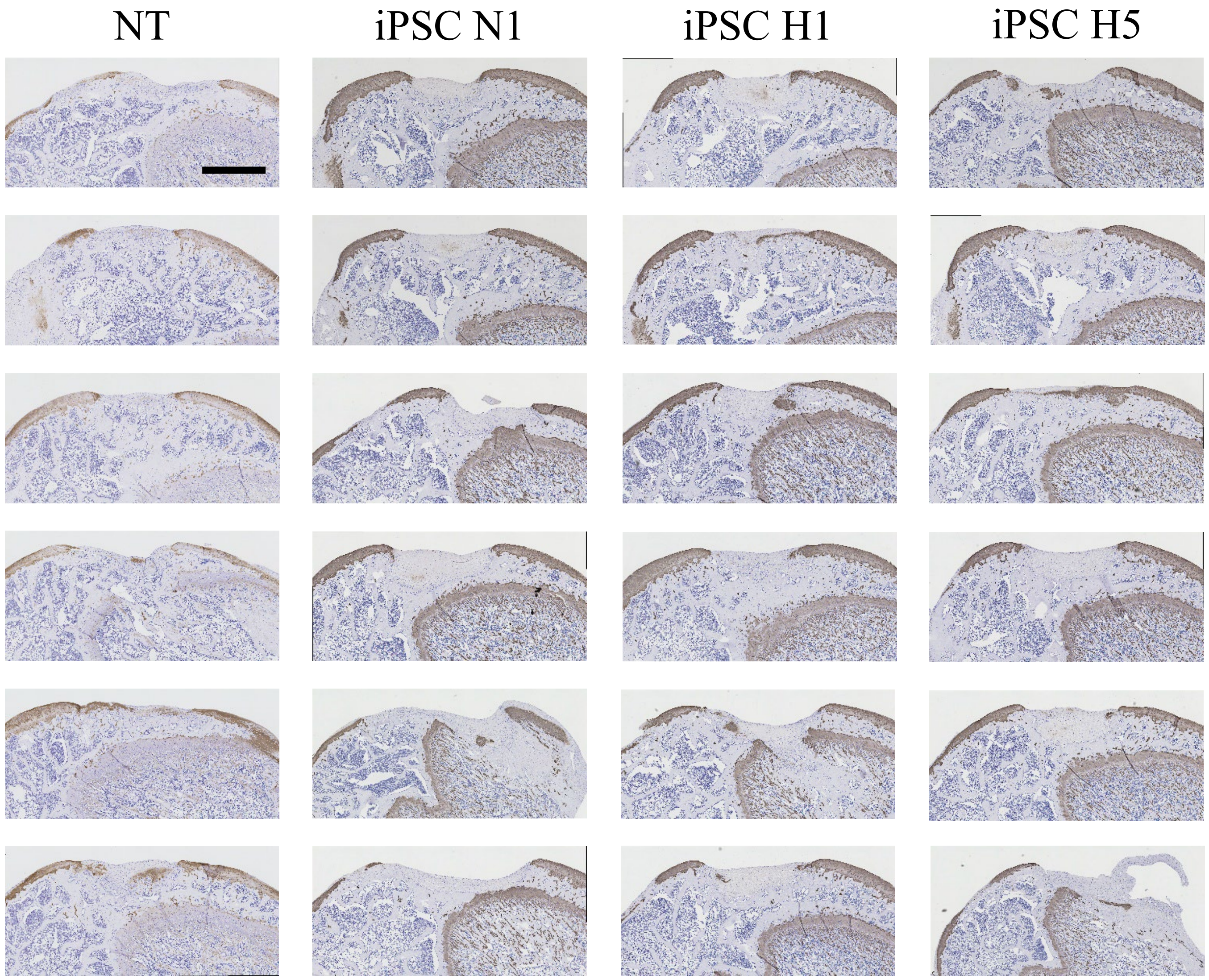


**Supplementary Figure 8. Histological evaluation at 4 weeks after transplantation of iPSC sheets in a xenogeneic orthotopic transplantation model using athymic nude rats.** Immunohistological staining for collagen type II for all animals. n = 6 per group. Scale bar = 500 µm.

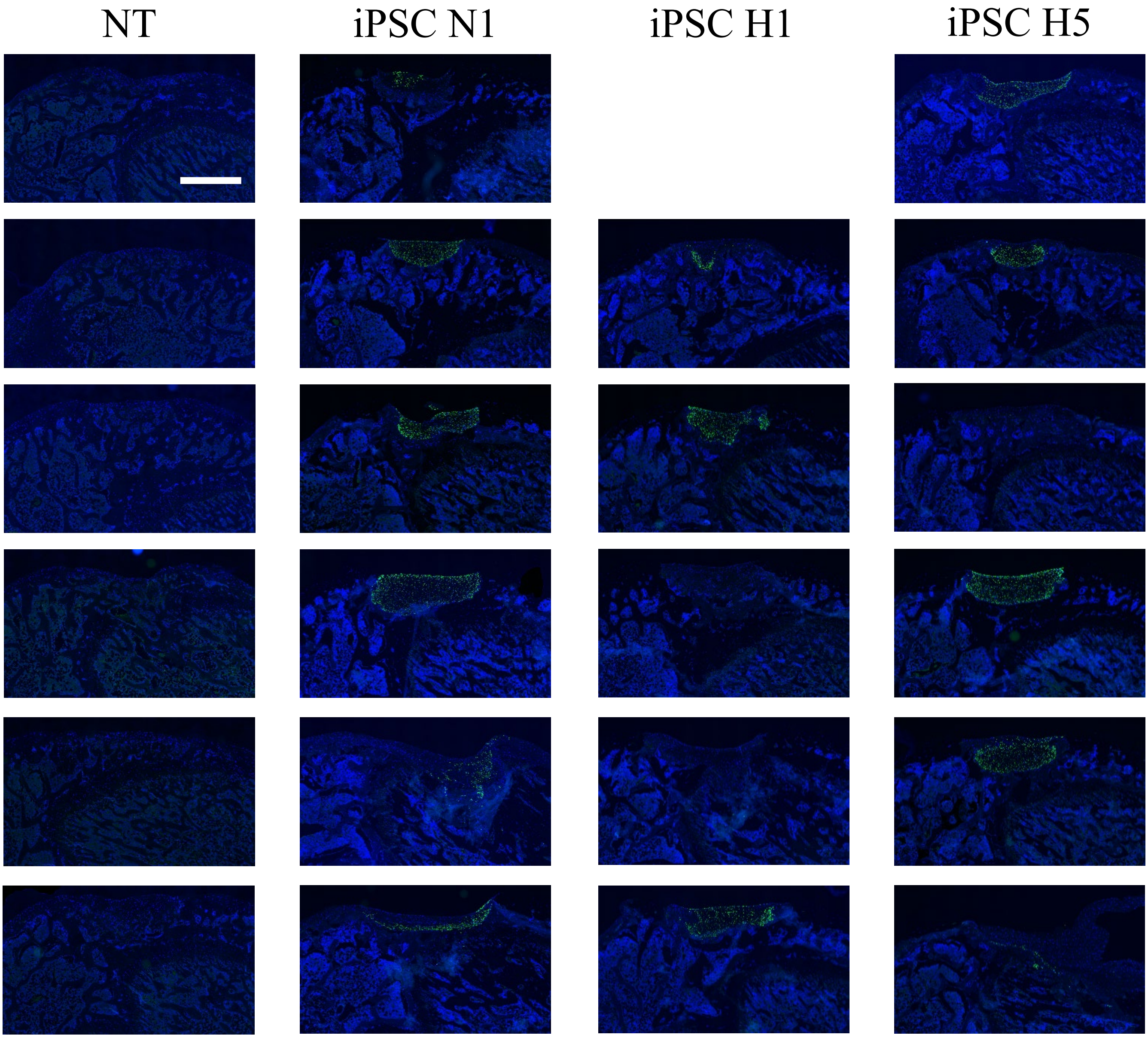


**Supplementary Figure 9. Histological evaluation at 4 weeks after transplantation of iPSC sheets in a xenogeneic orthotopic transplantation model using athymic nude rats.** Immunohistological staining for human-specific vimentin for all animals. n = 6 per group (one image was not obtained). Scale bar = 500 μm.

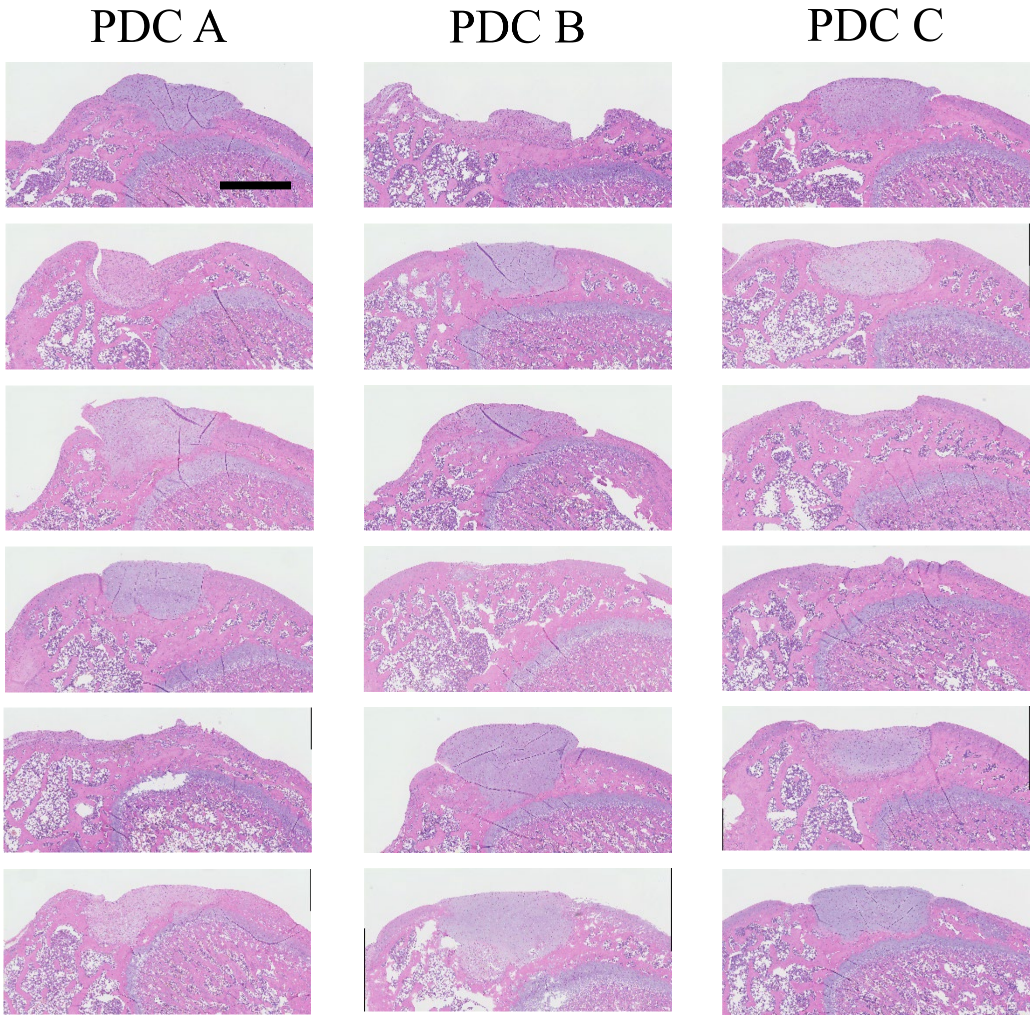


**Supplementary Figure 10. Histological evaluation at 4 weeks after transplantation of PDC sheets in a xenogeneic orthotopic transplantation model using athymic nude rats.** Hematoxylin and eosin staining results for all animals. n = 6 per group. Scale bar = 500 µm.

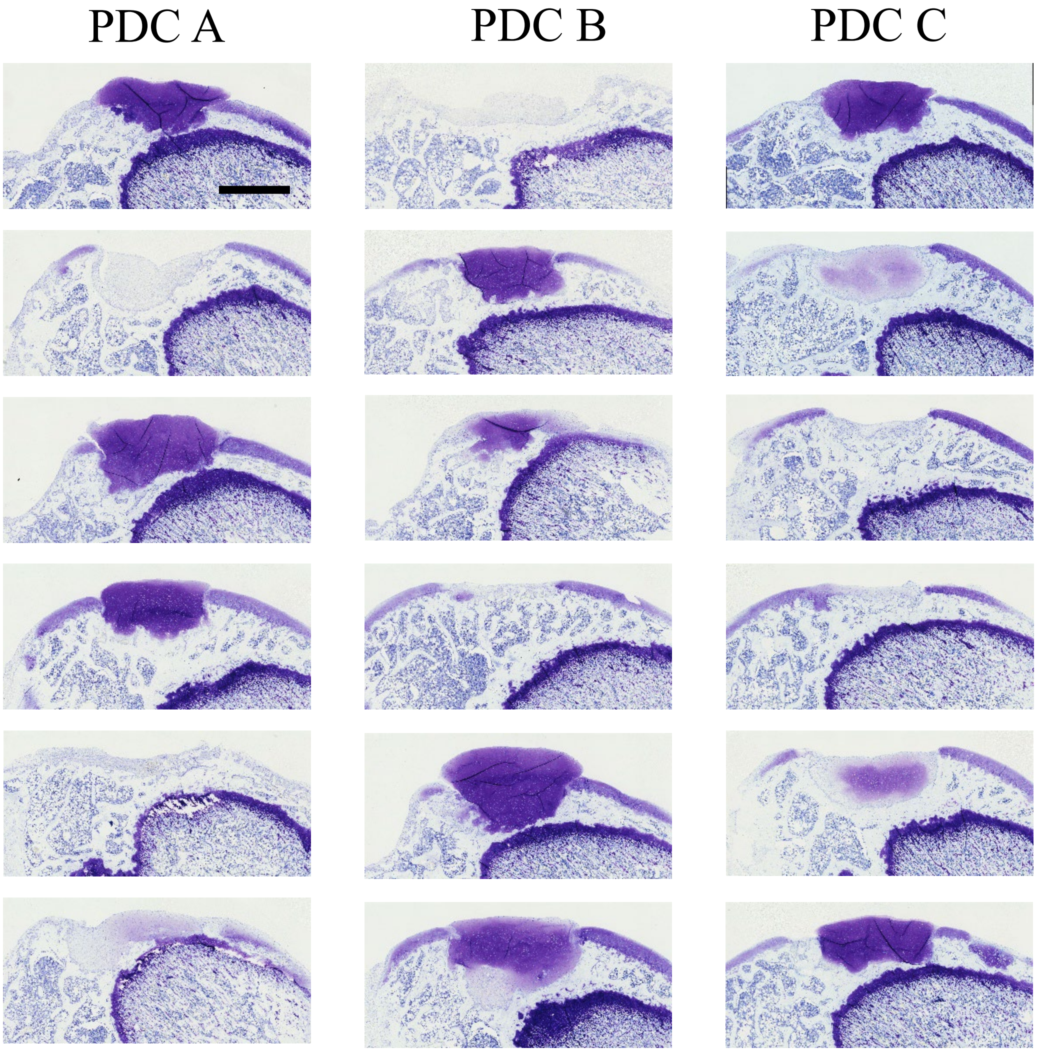


**Supplementary Figure 11. Histological evaluation at 4 weeks after transplantation of PDC sheets in a xenogeneic orthotopic transplantation model using athymic nude rats.** Toluidine blue staining results for all animals. n = 6 per group. Scale bar = 500 µm.

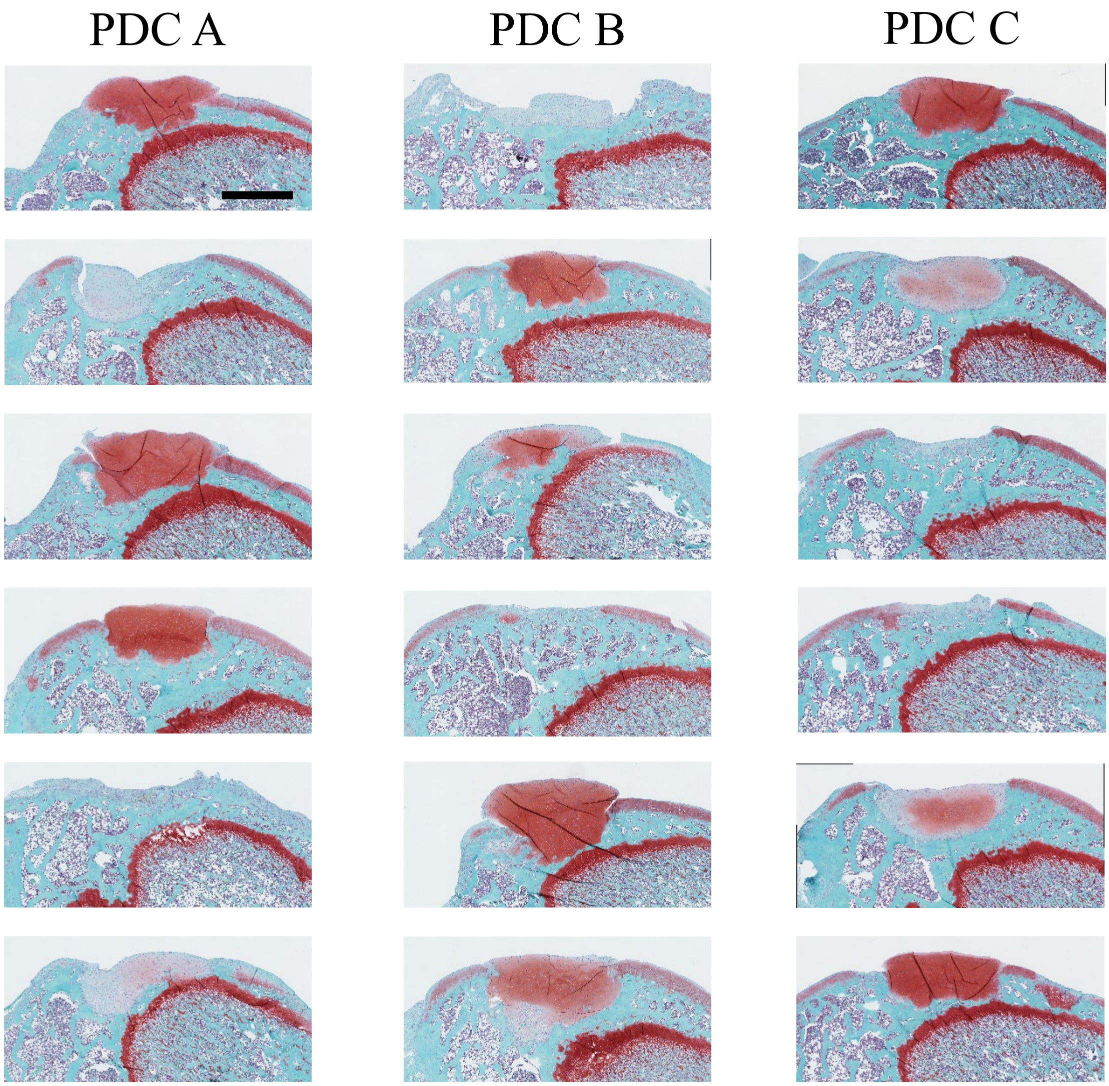


**Supplementary Figure 12. Histological evaluation at 4 weeks after transplantation of PDC sheets in a xenogeneic orthotopic transplantation model using athymic nude rats.** Safranin O staining results for all animals. n = 6 per group. Scale bar = 500 μm.

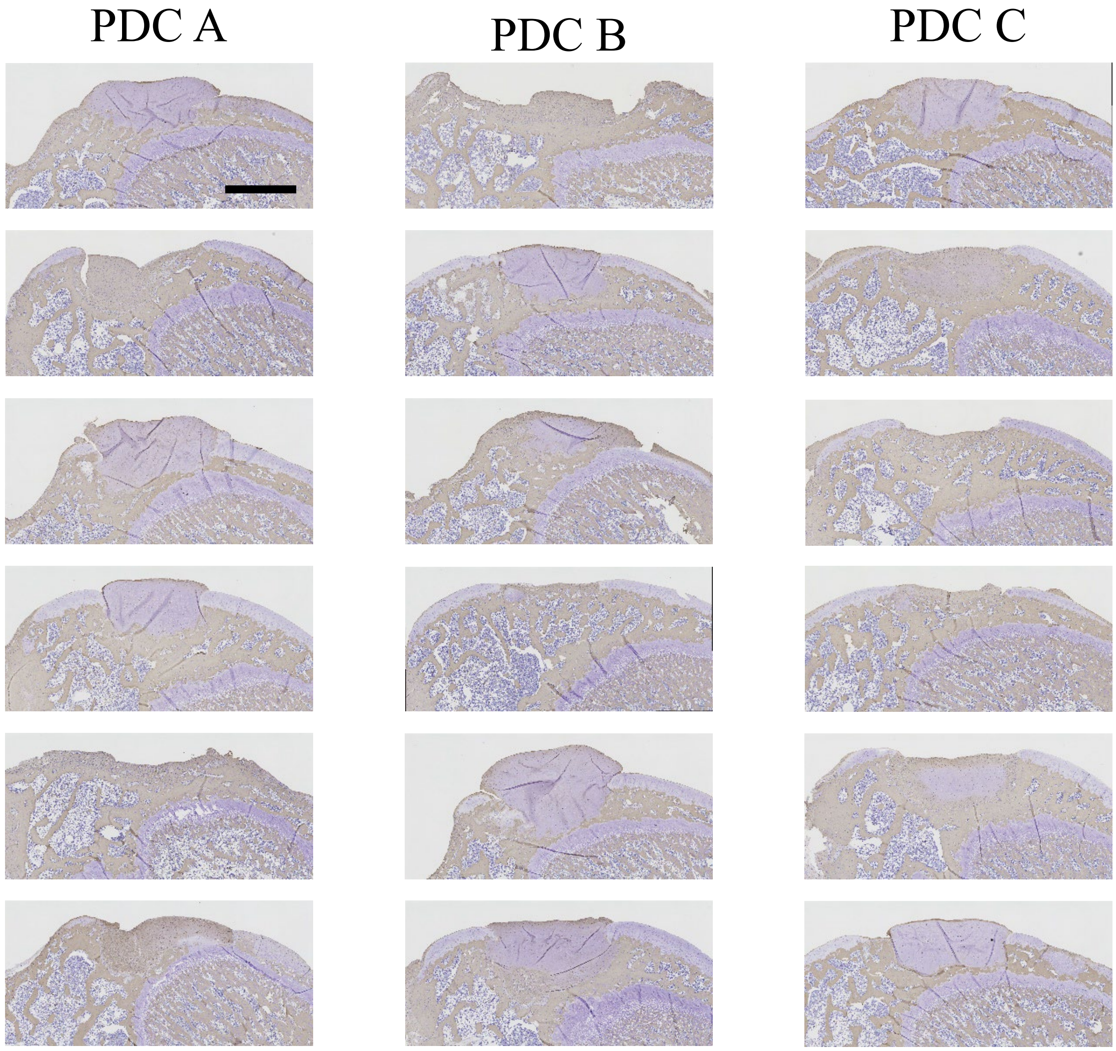


**Supplementary Figure 13. Histological evaluation at 4 weeks after transplantation of PDC sheets in a xenogeneic orthotopic transplantation model using athymic nude rats.** Immunohistological staining for collagen type I for all animals. n = 6 per group. Scale bar = 500 µm.

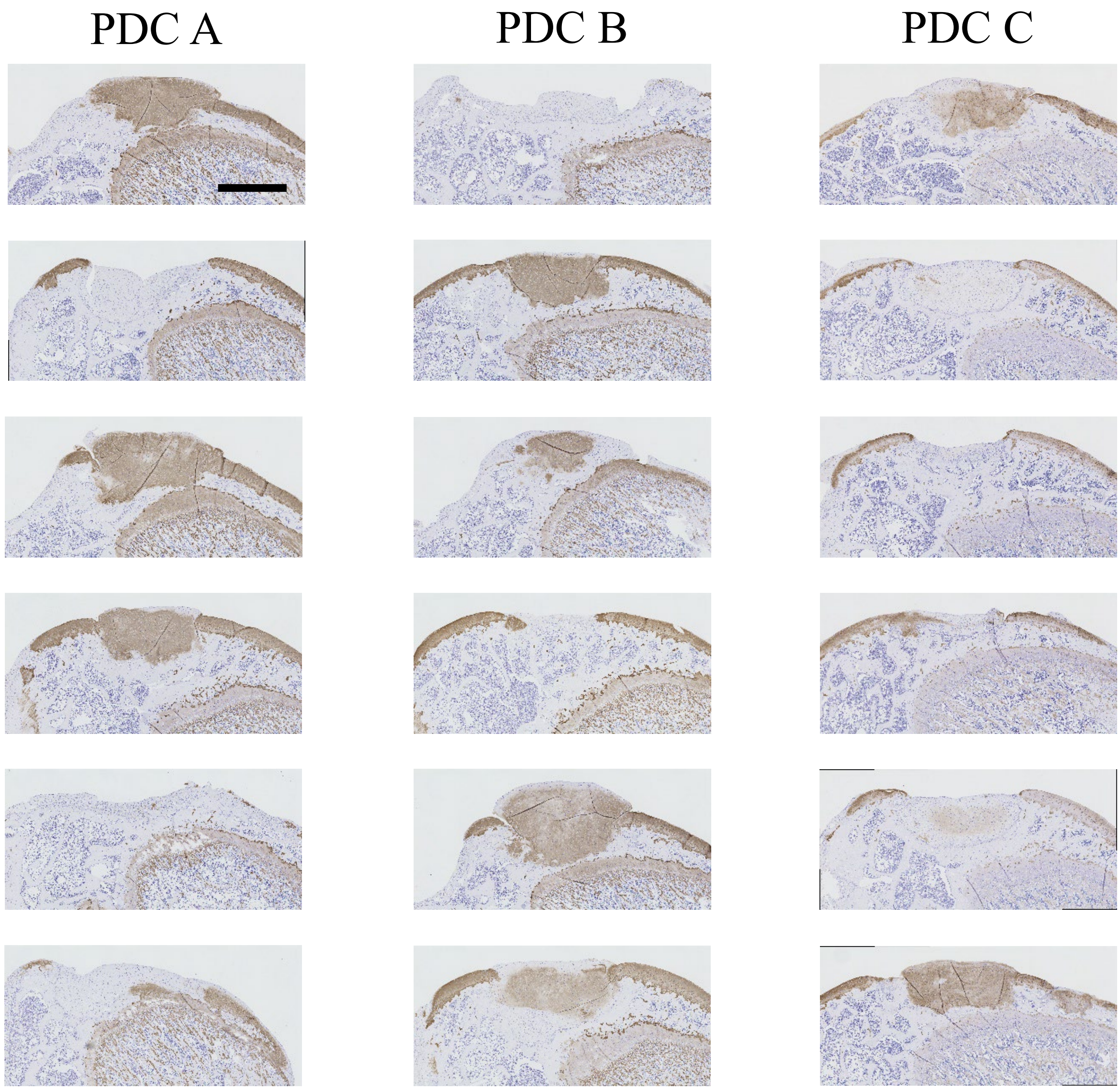


**Supplementary Figure 14. Histological evaluation at 4 weeks after transplantation of PDC sheets in a xenogeneic orthotopic transplantation model using athymic nude rats.** Immunohistological staining for collagen type II for all animals. n = 6 per group. Scale bar = 500 µm.

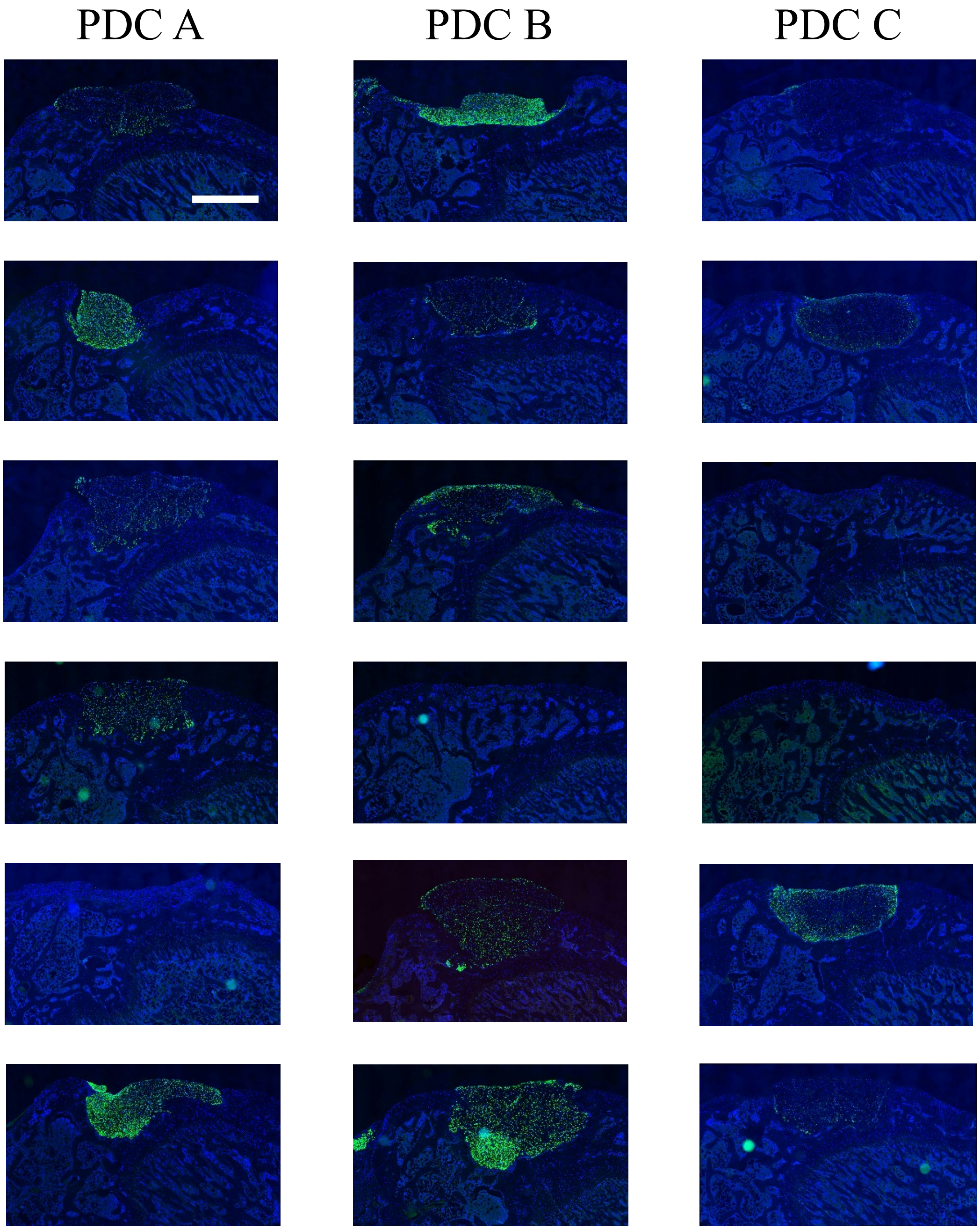


**Supplementary Figure 15. Histological evaluation at 4 weeks after transplantation of PDC sheets in a xenogeneic orthotopic transplantation model using athymic nude rats.** Immunohistological staining for human-specific vimentin for all animals. n = 6 per group. Scale bar = 500 µm.

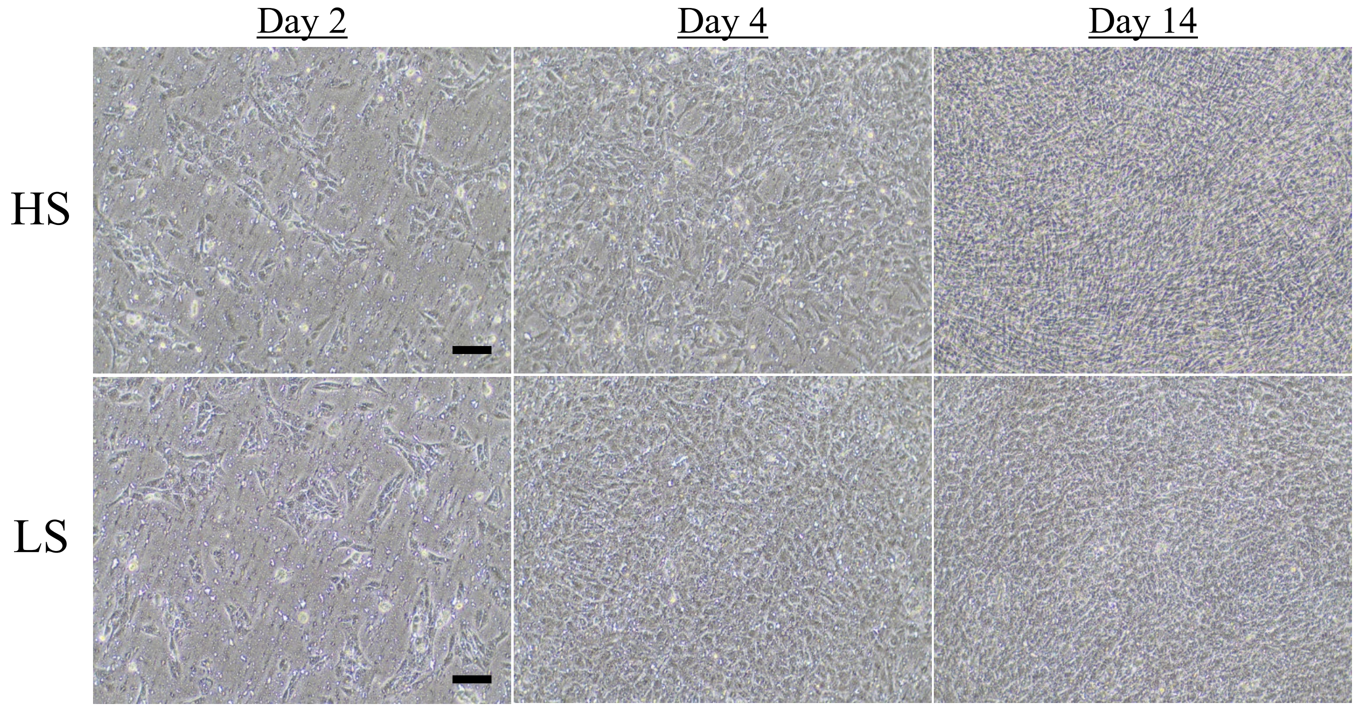


**Supplementary Figure 16.** Microscopic images (×10 magnification) of chondrocytes obtained from iPS-Carts and cultured in temperature-responsive culture inserts at days 2, 4, and 14. The chondrocytes observed at day 14 showed a polygonal shape for iPSC sheets fabricated under LS media, while a fibroblastic shape was observed for iPSC sheets fabricated under HS media. Scale bar = 100 µm. HS = high-serum; LS = low-serum.

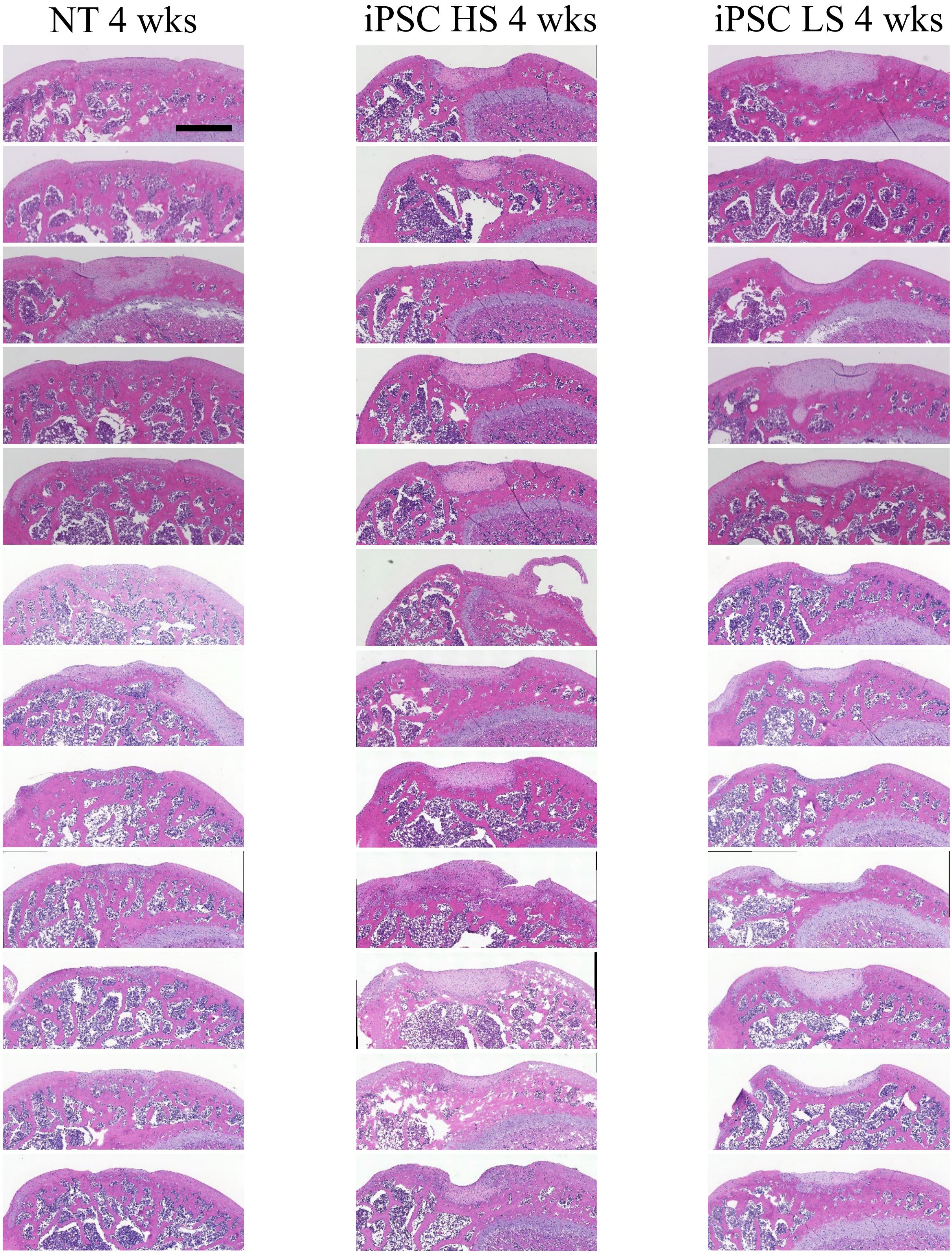


**Supplementary Figure 17. Histological evaluation at 4 weeks after transplantation of iPSC sheets in a xenogeneic orthotopic transplantation model using athymic nude rats.** Hematoxylin and eosin staining results for all animals. n = 12 per group. Scale bar = 500 µm.

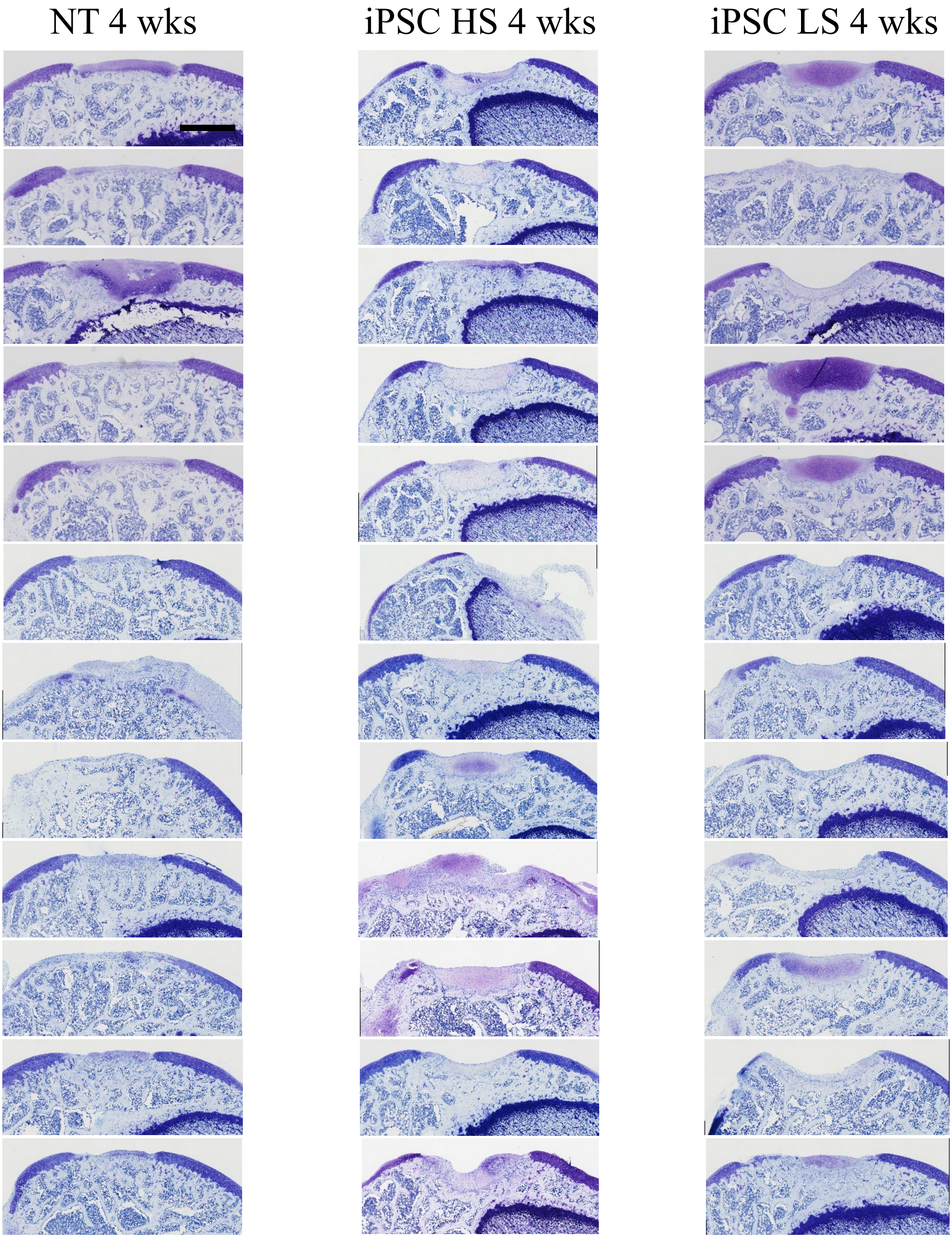


**Supplementary Figure 18. Histological evaluation at 4 weeks after transplantation of iPSC sheets in a xenogeneic orthotopic transplantation model using athymic nude rats.** Toluidine blue staining results for all animals. n = 12 per group. Scale bar = 500 µm.

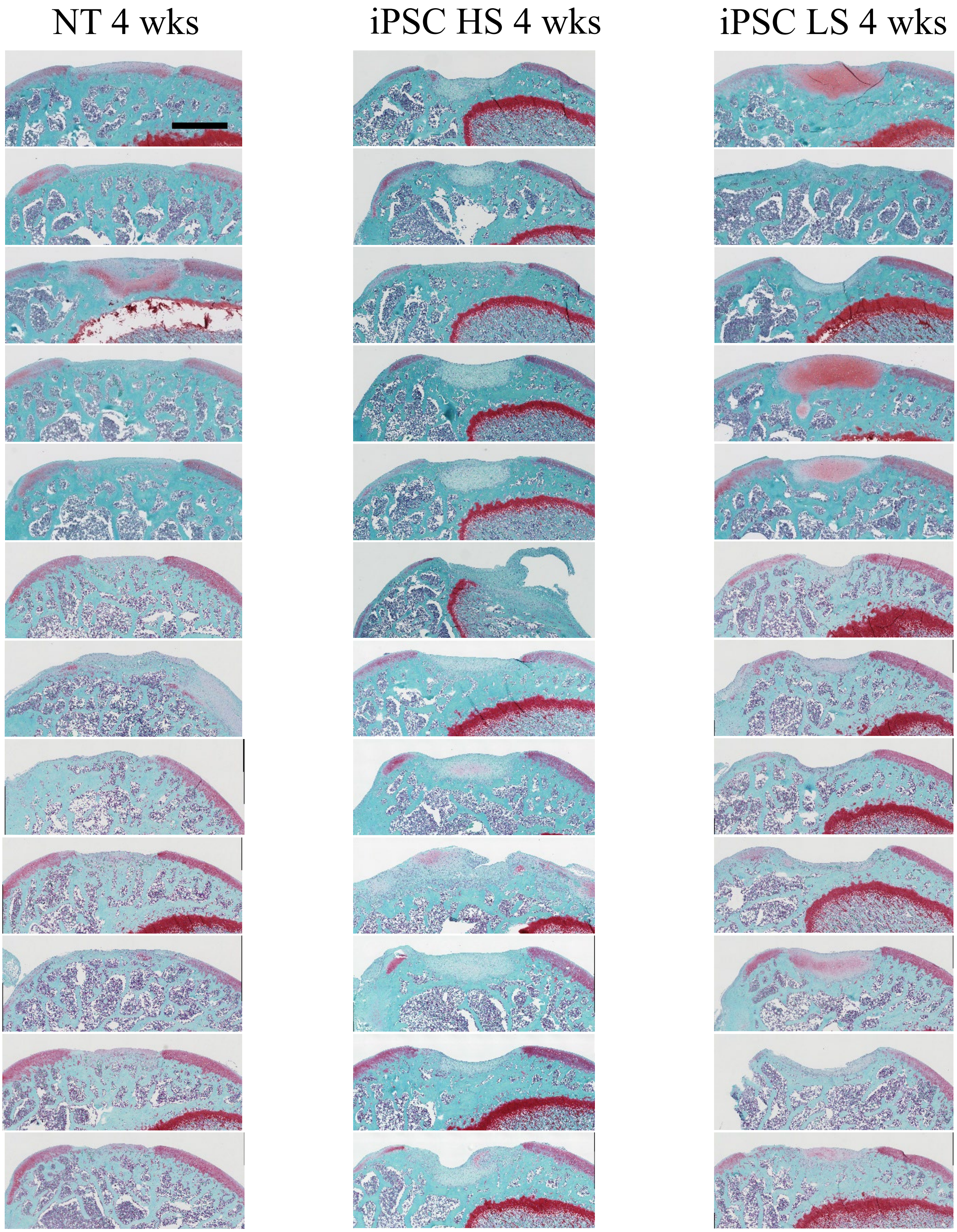


**Supplementary Figure 19. Histological evaluation at 4 weeks after transplantation of iPSC sheets in a xenogeneic orthotopic transplantation model using athymic nude rats.** Safranin O staining results for all animals. n = 12 per group. Scale bar = 500 μm.

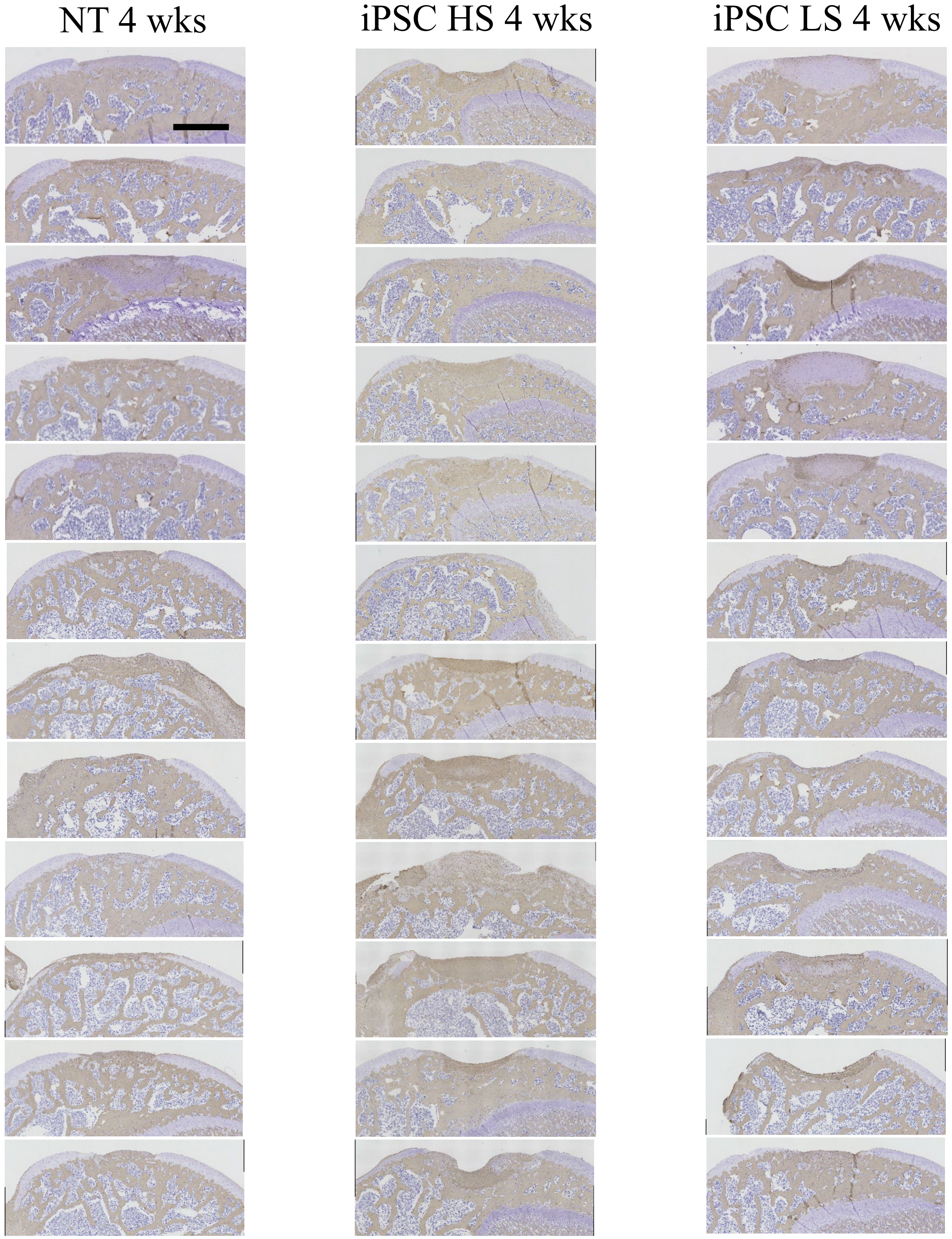


**Supplementary Figure 20. Histological evaluation at 4 weeks after transplantation of iPSC sheets in a xenogeneic orthotopic transplantation model using athymic nude rats.** Immunohistological staining for collagen type I for all animals. n = 12 per group. Scale bar = 500 μm.

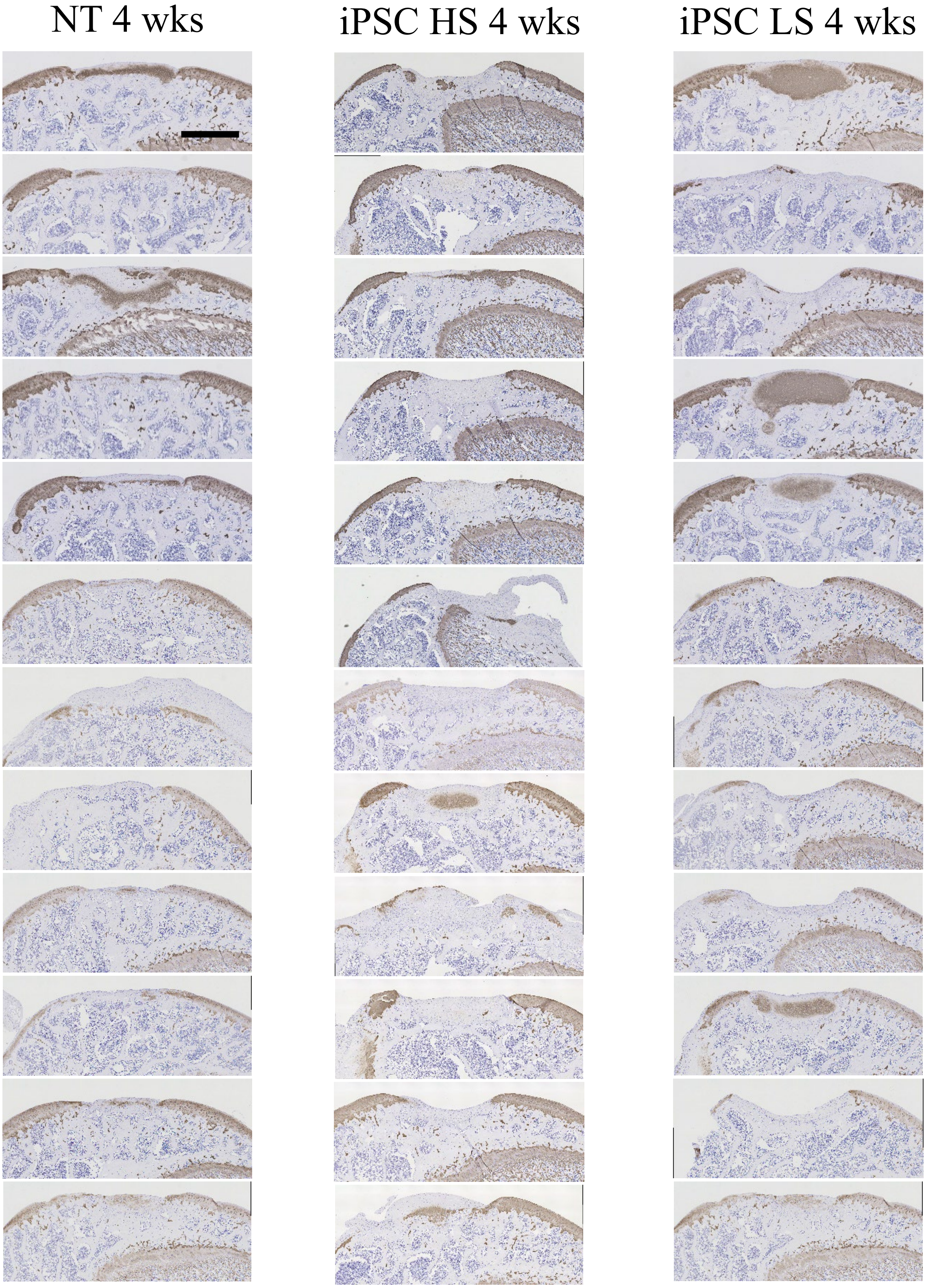


**Supplementary Figure 21. Histological evaluation at 4 weeks after transplantation of iPSC sheets in a xenogeneic orthotopic transplantation model using athymic nude rats.** Immunohistological staining for collagen type II for all animals. n = 12 per group. Scale bar = 500 μm.

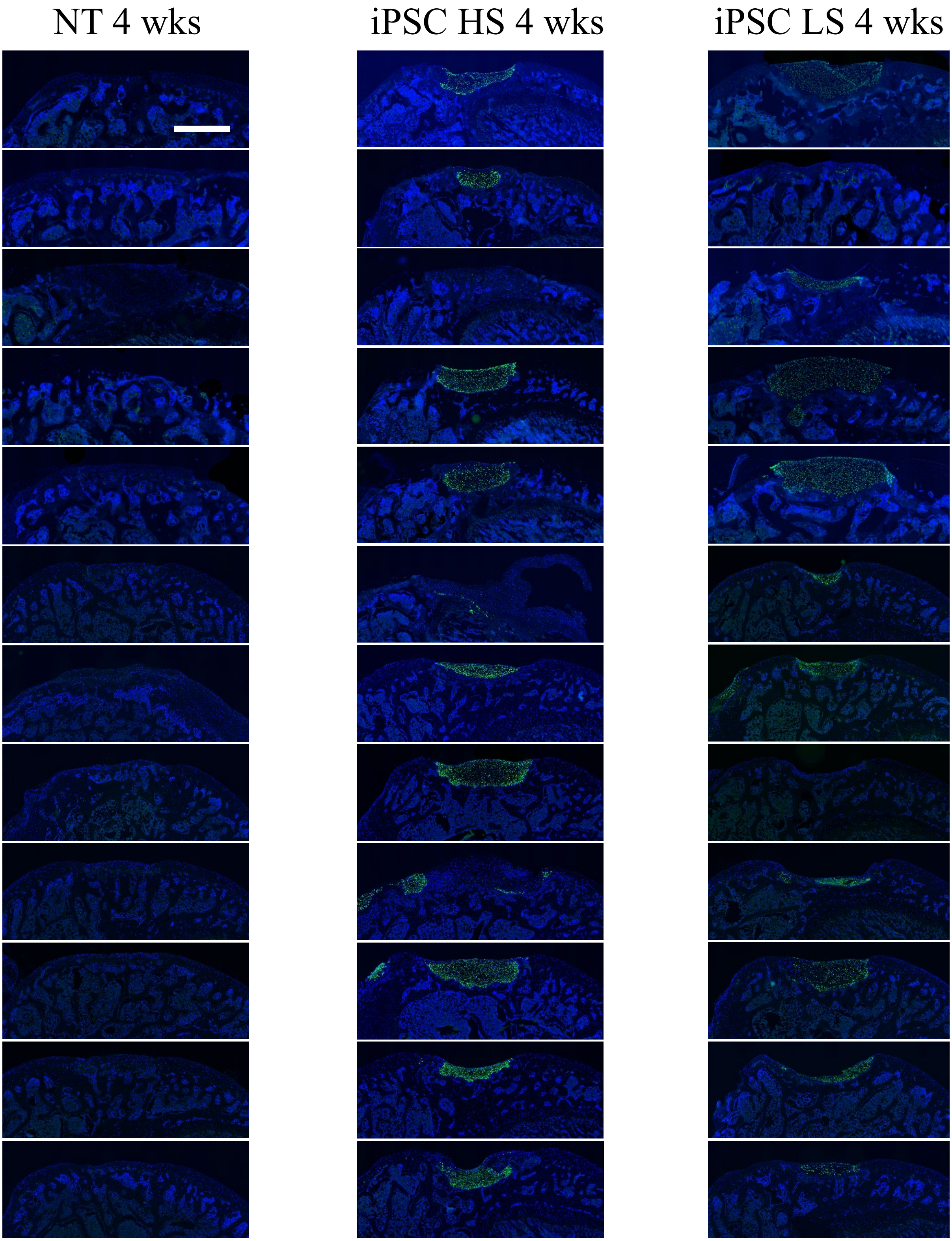


**Supplementary Figure 22. Histological evaluation at 4 weeks after transplantation of iPSC sheets in a xenogeneic orthotopic transplantation model using athymic nude rats.** Immunohistological staining for human-specific vimentin for all animals. n = 12 per group. Scale bar = 500 µm.

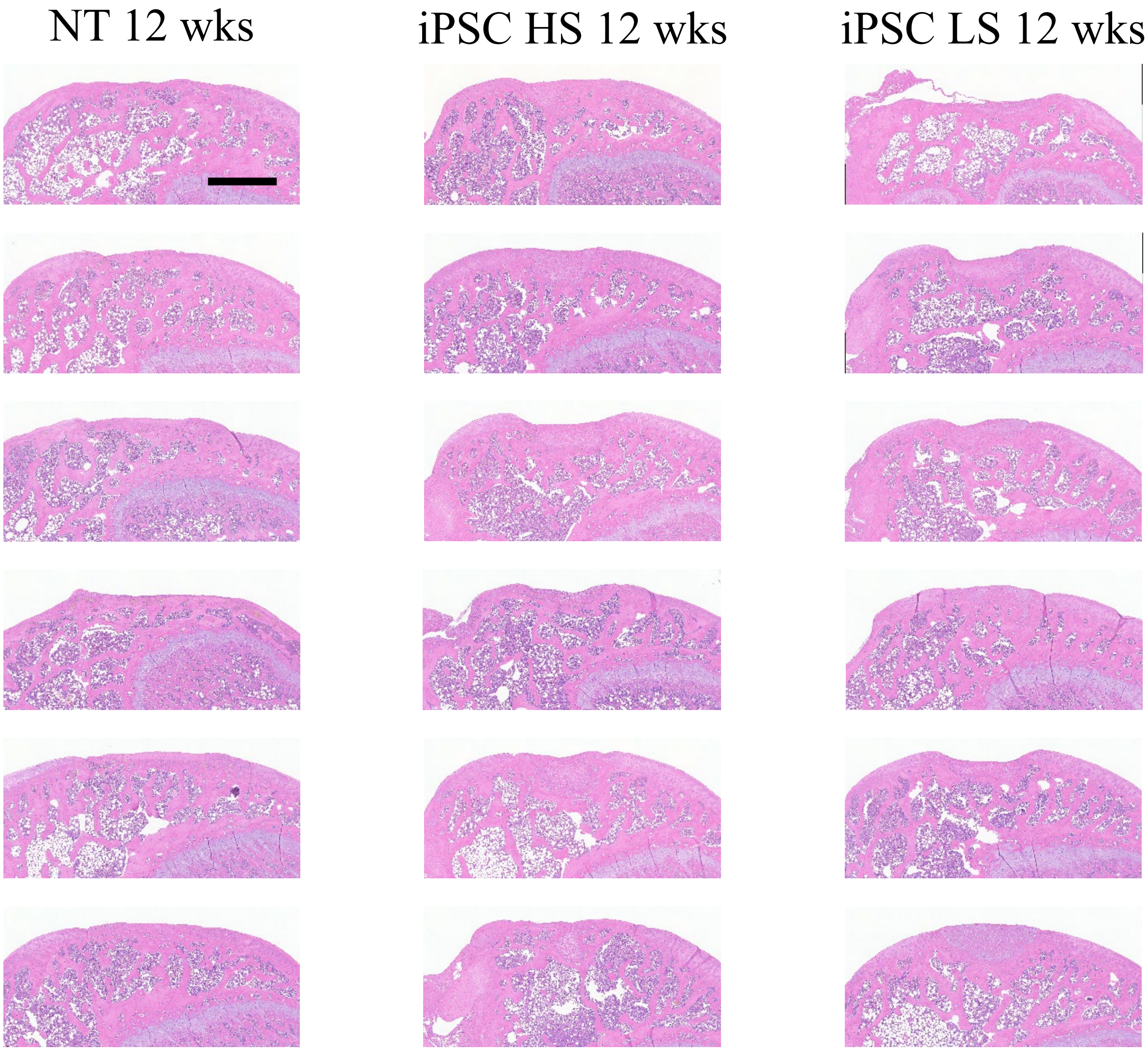


**Supplementary Figure 23. Histological evaluation at 12 weeks after transplantation of iPSC sheets in a xenogeneic orthotopic transplantation model using athymic nude rats.** Hematoxylin and eosin staining results for all animals. n = 6 per group. Scale bar = 500 µm.

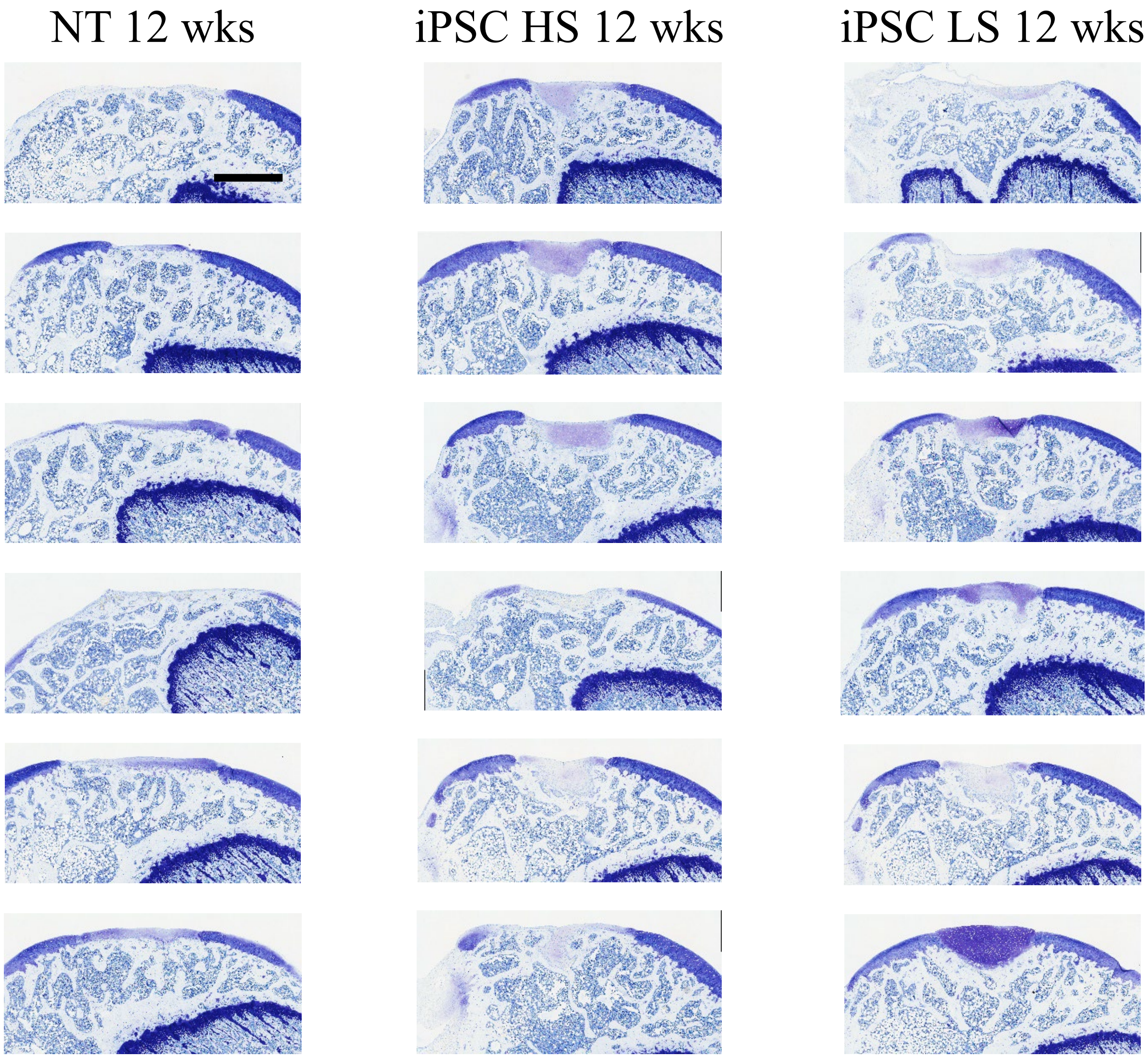


**Supplementary Figure 24. Histological evaluation at 12 weeks after transplantation of iPSC sheets in a xenogeneic orthotopic transplantation model using athymic nude rats.** Toluidine blue staining results for all animals. n = 6 per group. Scale bar = 500 μm.

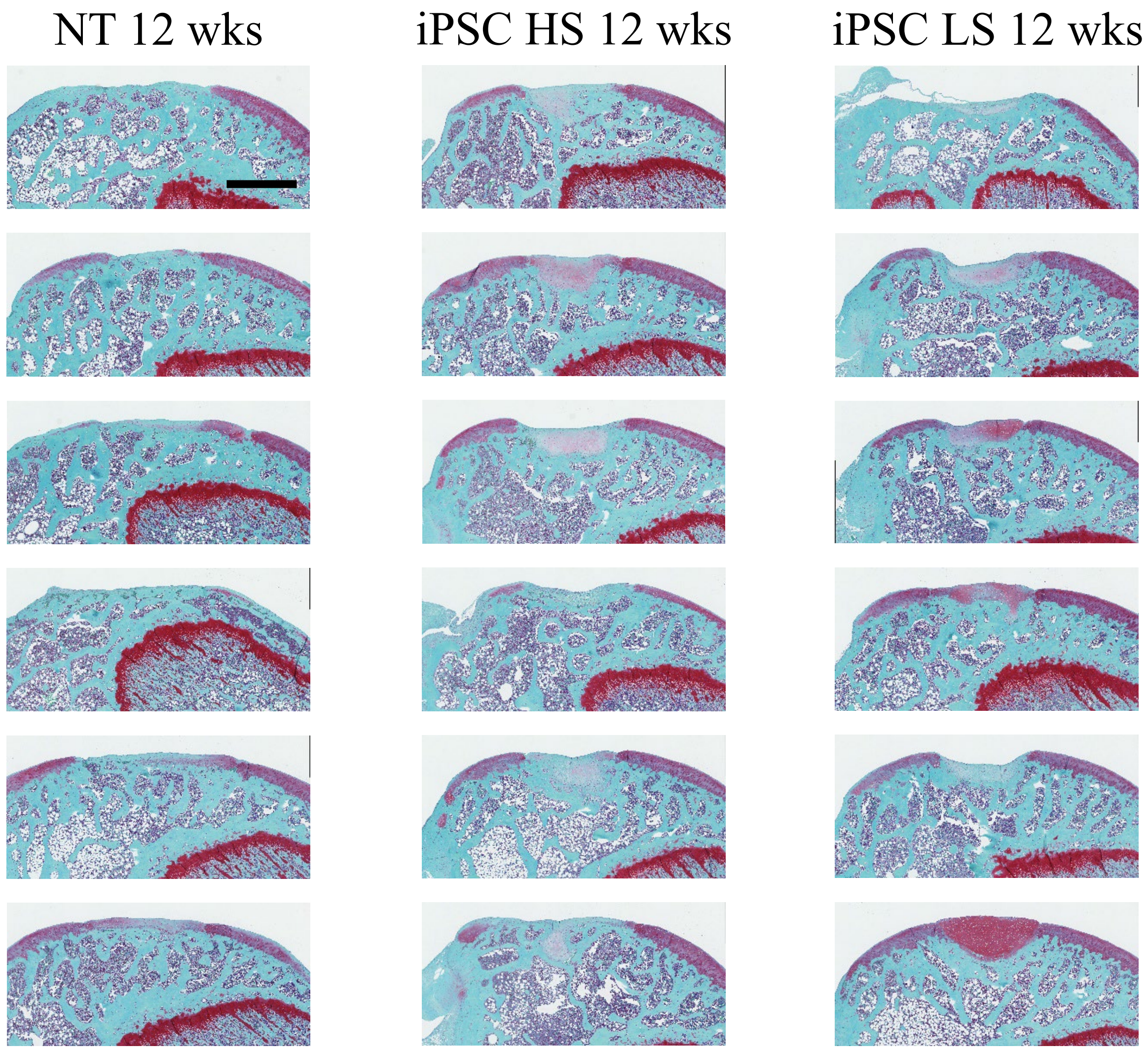


**Supplementary Figure 25. Histological evaluation at 12 weeks after transplantation of iPSC sheets in a xenogeneic orthotopic transplantation model using athymic nude rats.** Safranin O staining results for all animals. n = 6 per group. Scale bar = 500 µm.

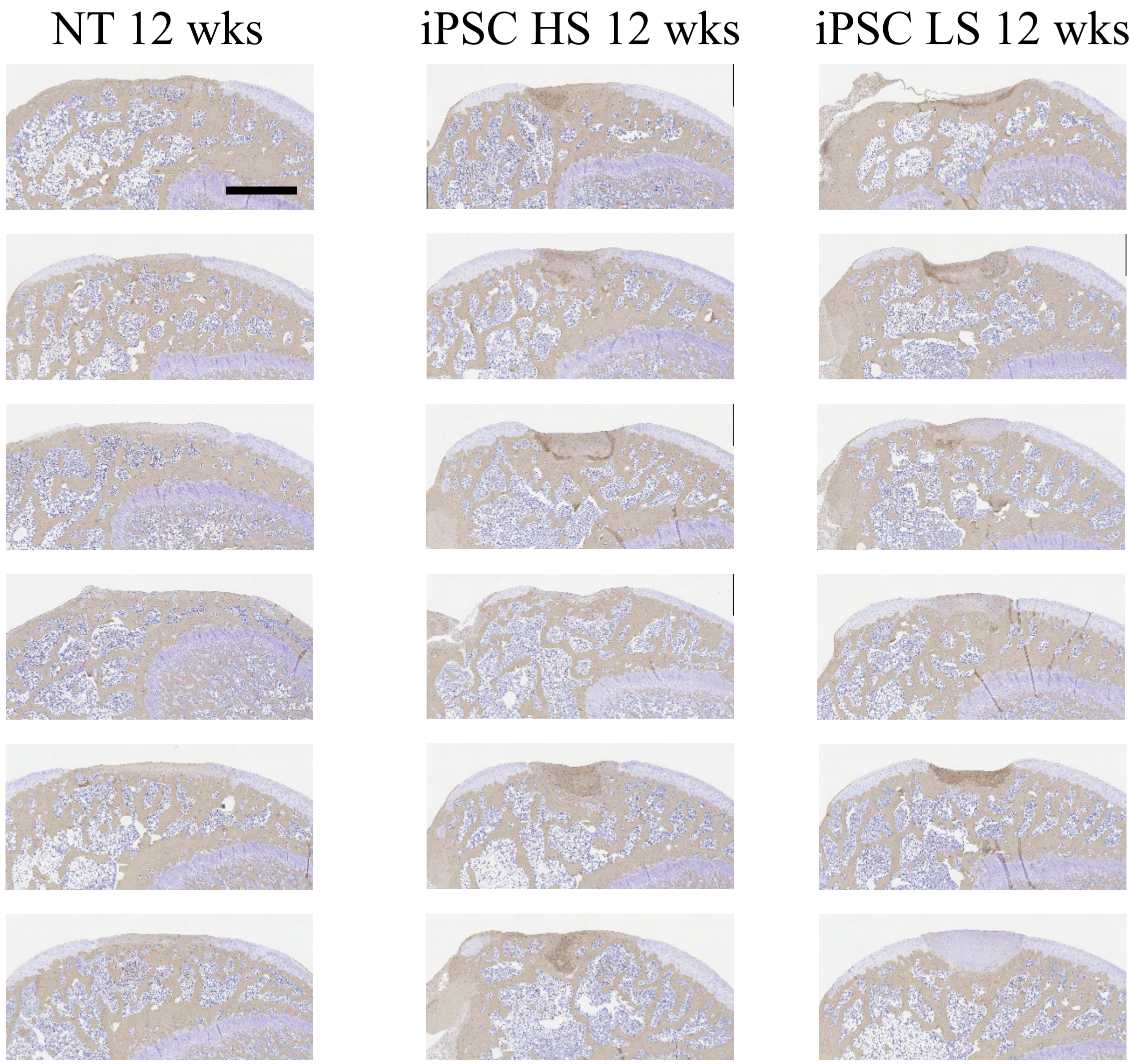


**Supplementary Figure 26. Histological evaluation at 12 weeks after transplantation of iPSC sheets in a xenogeneic orthotopic transplantation model using athymic nude rats.** Immunohistological staining for collagen type I for all animals. n = 6 per group. Scale bar = 500 µm.

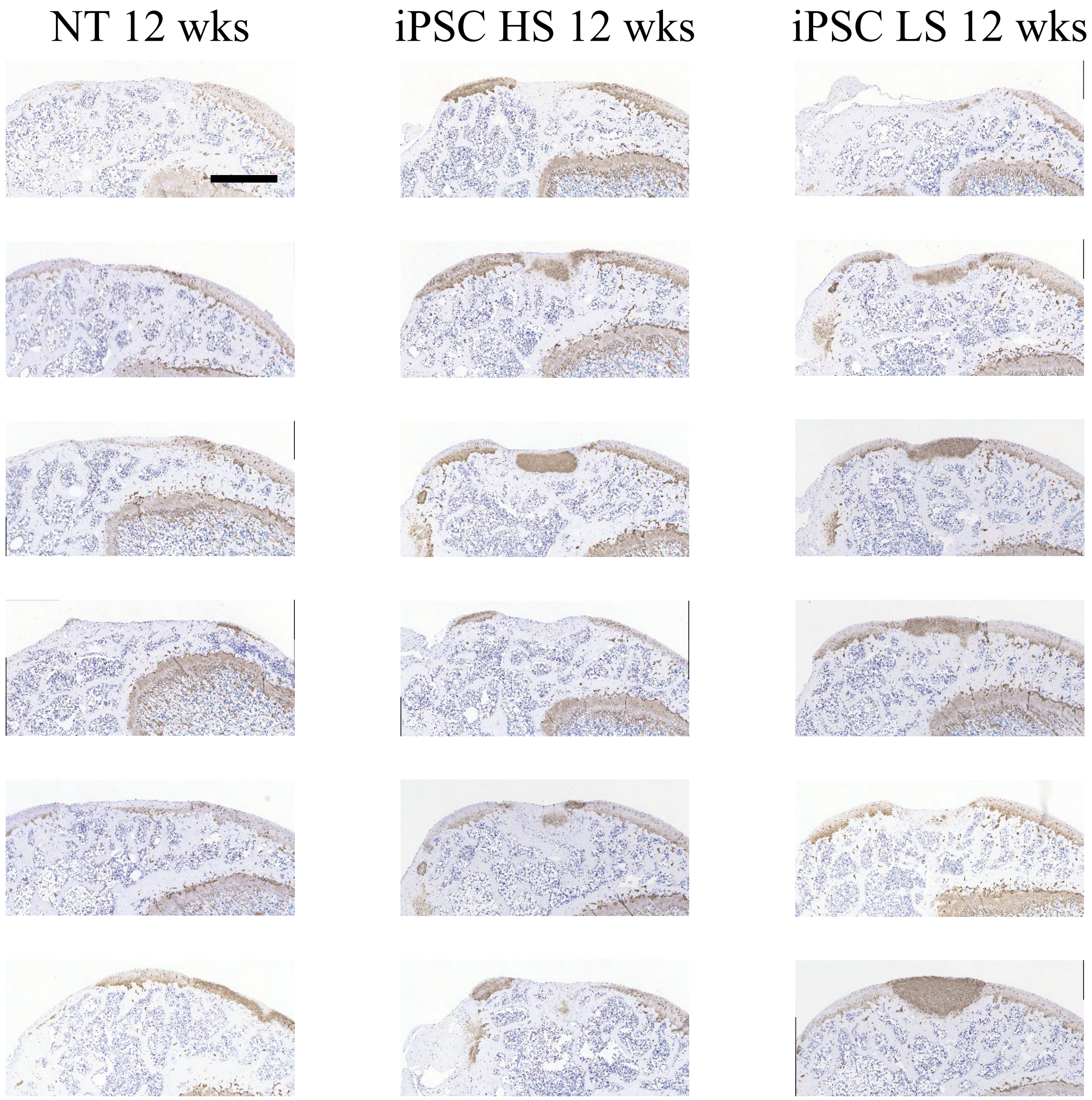


**Supplementary Figure 27. Histological evaluation at 12 weeks after transplantation of iPSC sheets in a xenogeneic orthotopic transplantation model using athymic nude rats.** Immunohistological staining for collagen type II for all animals. n = 6 per group. Scale bar = 500 μm.

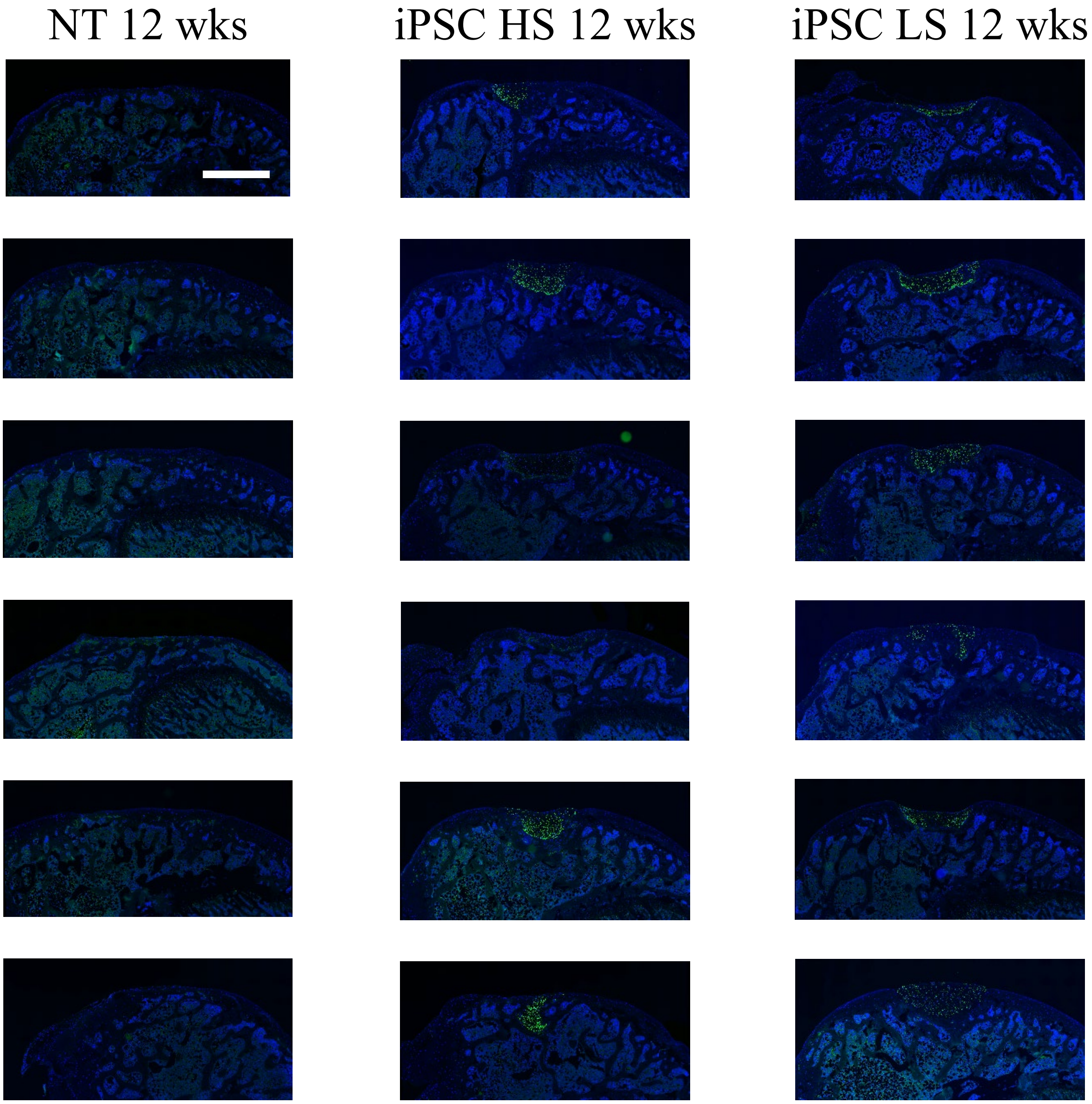


**Supplementary Figure 28. Histological evaluation at 12 weeks after transplantation of iPSC sheets in a xenogeneic orthotopic transplantation model using athymic nude rats.** Immunohistological staining for human-specific vimentin for all animals. n = 6 per group. Scale bar = 500 μm.

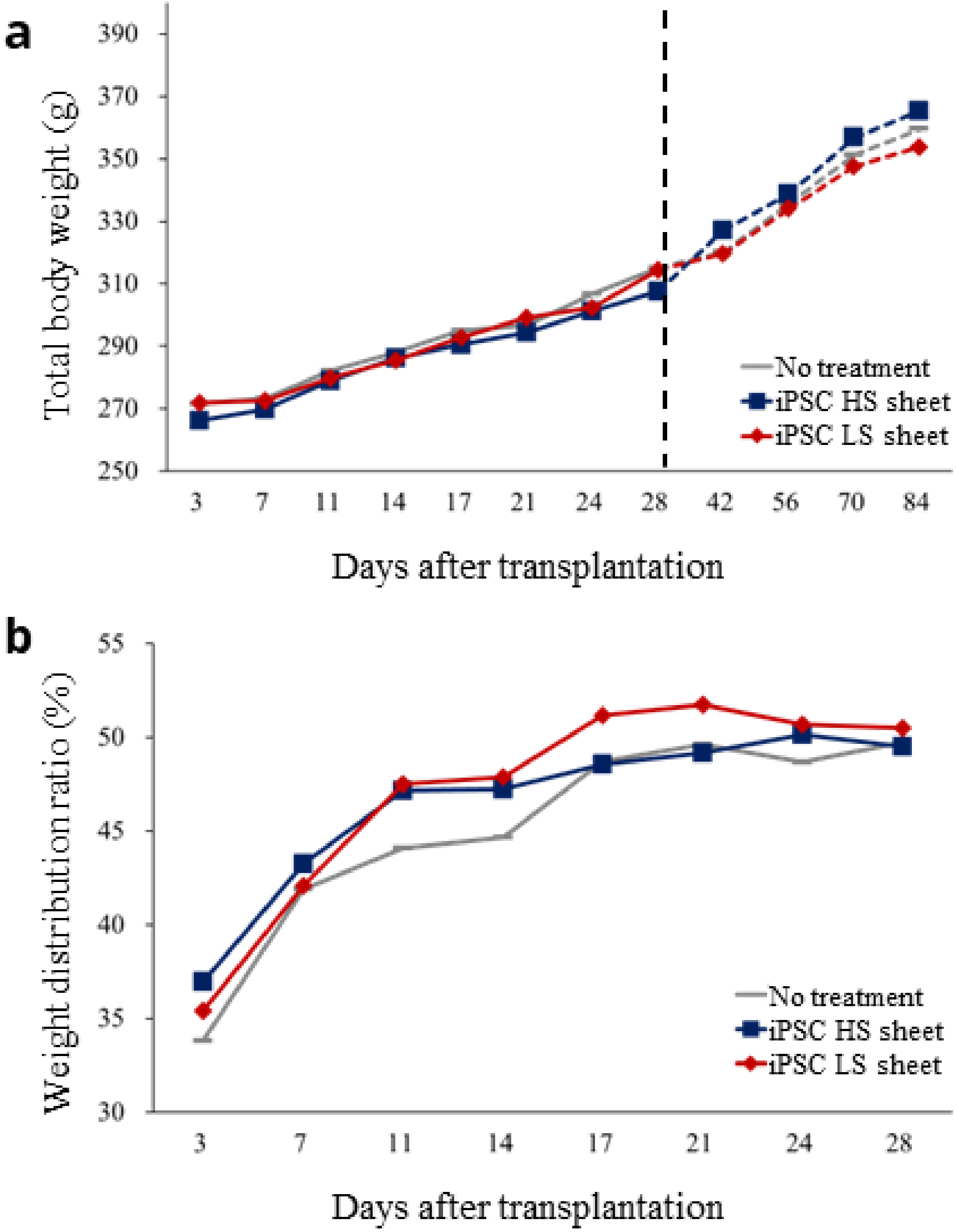


**Supplementary Figure 29. Total body weight and weight distribution ratios for each transplantation group.** (a) No significant differences were detected for total body weight among transplantation groups (n = 18 up to day 28; n = 6 thereafter). (b) No significant differences were detected for the weight distribution ratios among transplantation groups (n = 18), but the iPSC sheet transplantation groups had higher weight distribution ratios at days 11 and 14, suggesting faster recovery from pain.

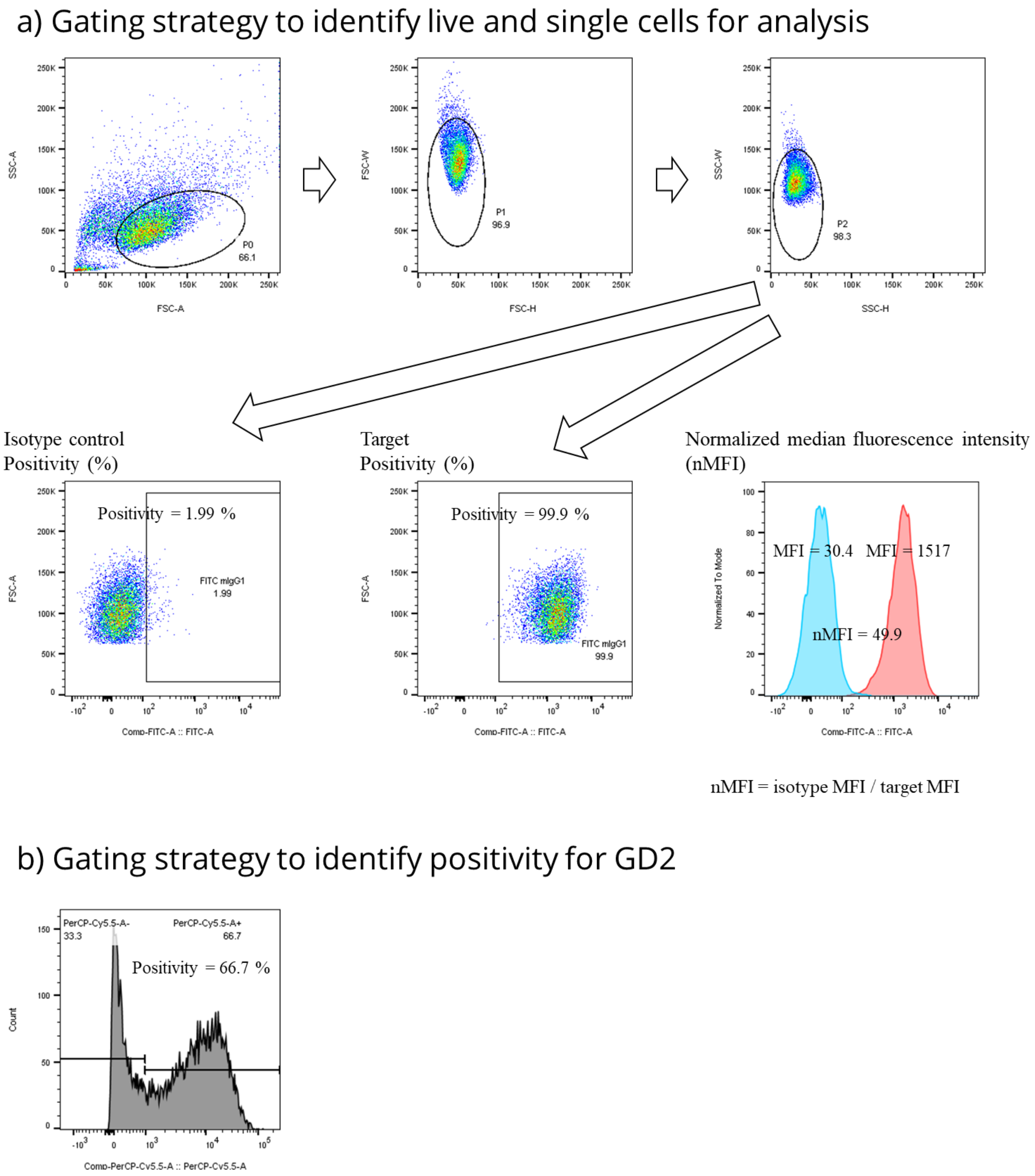


**Supplementary Figure 30. Gating strategies used to identify cell positivity and normalized median fluorescent intensities.** (A) Positivity was determined by setting the gate for the corresponding isotype control to <2% positivity. Normalized median fluorescence intensity (nMFI) was calculated from the isotype MFIs and target MFIs for each target and its corresponding isotype control. Examples of positivity and nMFI calculations are given for CD73. (B) Gating strategy to identify positivity for GD2 with bimodal expression is shown.
