## Supplementary Tables for "Enhancing the hyaline cartilage regenerative potential of induced pluripotent stem cell-derived chondrocyte cell sheets"

**Supplementary Table 1.** Results from the preliminary screening test for cell surface markers using BD's Human Lyoplate Screening Panel

| | | Positivity (%) | | nMFI | |
|---|---|---|---|---|---|
| **#** | **Target** | **HS** | **LS** | **HS** | **LS** |
| 1 | CD1a | 1.2 | 2.25 | 0.91 | 1.00 |
| 2 | CD1b | 1.89 | 2.8 | 0.91 | 1.08 |
| 3 | CD1d | 1.37 | 2.51 | 0.91 | 1.00 |
| 4 | CD2 | 1.52 | 2.1 | 0.91 | 1.00 |
| 5 | CD3 | 1.52 | 1.92 | 0.91 | 1.00 |
| 6 | CD4 | 1.26 | 2.75 | 0.91 | 1.08 |
| 7 | CD4v4 | 1.25 | 3.9 | 0.91 | 1.17 |
| 8 | CD5 | 1.02 | 2.15 | 0.91 | 1.00 |
| 9 | CD6 | 1.66 | 2.06 | 0.91 | 1.00 |
| 10 | CD7 | 1.5 | 3.45 | 0.91 | 1.08 |
| 11 | CD8a | 1.51 | 2.59 | 0.91 | 1.00 |
| 12 | CD8b | 1.37 | 1.5 | 0.91 | 1.00 |
| 13 | CD9 | 94.5 | 99.3 | 57.03 | 168.38 |
| 14 | CD10 | 36.6 | 80.6 | 3.91 | 13.57 |
| 15 | CD11a | 1.72 | 1.66 | 1.00 | 0.92 |
| 16 | CD11b | 1.29 | 1.74 | 0.91 | 1.00 |
| 17 | CD11c | 1.43 | 2.84 | 0.91 | 1.08 |
| 18 | CD13 | 98.4 | 94.9 | 97.73 | 96.69 |
| 19 | CD14 | 1.61 | 1.77 | 0.91 | 1.00 |

| | | | | | |
|---|---|---|---|---|---|
| 20 | CD15 | 0.94 | 1.91 | 0.82 | 0.92 |
| 21 | CD15s | 0.89 | 2.02 | 0.82 | 0.92 |
| 22 | CD16 | 1.19 | 2.27 | 0.80 | 1.00 |
| 23 | CD18 | 5.11 | 11.2 | 1.60 | 2.17 |
| 24 | CD19 | 1.41 | 1.9 | 0.91 | 1.00 |
| 25 | CD20 | 1.18 | 2.17 | 0.82 | 1.00 |
| 26 | CD21 | 1.14 | 2 | 0.91 | 0.92 |
| 27 | CD22 | 1.28 | 2.05 | 0.91 | 1.00 |
| 28 | CD23 | 1.15 | 1.75 | 0.80 | 0.92 |
| 29 | CD24 | 3.56 | 7.29 | 1.00 | 1.25 |
| 30 | CD25 | 1.32 | 2.37 | 0.91 | 1.08 |
| 31 | CD26 | 5.19 | 70.8 | 1.20 | 10.06 |
| 32 | CD27 | 1.35 | 2.6 | 0.80 | 1.08 |
| 33 | CD28 | 1.53 | 1.9 | 0.91 | 1.08 |
| 34 | CD29 | 96.7 | 99.3 | 33.28 | 53.44 |
| 35 | CD30 | 1.6 | 2.48 | 0.91 | 1.08 |
| 36 | CD31 | 1.64 | 1.89 | 0.91 | 0.92 |
| 37 | CD32 | 1.44 | 1.74 | 0.82 | 0.92 |
| 38 | CD33 | 1.47 | 1.81 | 0.91 | 0.92 |
| 39 | CD34 | 3.5 | 8.72 | 1.60 | 1.92 |
| 40 | CD35 | 1.75 | 2.73 | 1.00 | 1.08 |
| 41 | CD36 | 1.24 | 2.11 | 0.91 | 1.00 |
| 42 | CD37 | 0.96 | 2.63 | 0.80 | 1.08 |
| 43 | CD38 | 1.58 | 4.66 | 0.91 | 1.08 |

| | | | | | |
|---|---|---|---|---|---|
| 44 | CD39 | 1.32 | 2.62 | 0.82 | 1.00 |
| 45 | CD40 | 2.81 | 2.9 | 1.10 | 1.08 |
| 46 | CD41a | 1.67 | 1.94 | 0.91 | 1.00 |
| 47 | CD41b | 1.46 | 3.23 | 1.00 | 1.08 |
| 48 | CD42a | 2.17 | 2.71 | 1.00 | 1.17 |
| 49 | CD42b | 1.39 | 2.23 | 0.91 | 1.00 |
| 50 | CD43 | 1.78 | 2.07 | 0.91 | 1.00 |
| 51 | CD44 | 100 | 99.9 | 1101.21 | 791.17 |
| 52 | CD45 | 1.31 | 1.85 | 0.91 | 1.08 |
| 53 | CD45RA | 1.34 | 1.65 | 0.82 | 0.92 |
| 54 | CD45RB | 1.06 | 2.12 | 0.80 | 1.00 |
| 55 | CD45RO | 1.71 | 1.91 | 0.91 | 1.00 |
| 56 | CD46 | 99.7 | 99.8 | 41.56 | 40.26 |
| 57 | CD47 | 99.9 | 99.9 | 62.19 | 94.94 |
| 58 | CD48 | 1.05 | 1.74 | 0.82 | 1.00 |
| 59 | CD49a | 49.4 | 43.1 | 5.02 | 3.92 |
| 60 | CD49b | 44.9 | 88.9 | 4.62 | 12.99 |
| 61 | CD49c | 99.7 | 98.3 | 148.05 | 39.35 |
| 62 | CD49d | 25.3 | 13.1 | 2.80 | 1.92 |
| 63 | CD49e | 99.7 | 99.9 | 95.00 | 82.34 |
| 64 | CD50 | 1.59 | 3.34 | 1.09 | 1.25 |
| 65 | CD51/61 | 80.2 | 56.1 | 11.56 | 5.25 |
| 66 | CD53 | 1.04 | 1.8 | 0.80 | 0.92 |
| 67 | CD54 | 79.8 | 74.3 | 17.16 | 12.14 |

| | | | | | |
|---|---|---|---|---|---|
| 68 | CD55 | 98.3 | 98.3 | 57.03 | 44.81 |
| 69 | CD56 | 69.6 | 90.1 | 8.91 | 18.38 |
| 70 | CD57 | 7.51 | 7.11 | 1.37 | 1.46 |
| 71 | CD58 | 99.9 | 99.9 | 56.25 | 83.12 |
| 72 | CD59 | 100 | 100 | 1187.97 | 820.78 |
| 73 | CD61 | 78.9 | 44 | 10.47 | 4.17 |
| 74 | CD62E | 1.18 | 2.19 | 0.91 | 1.00 |
| 75 | CD62L | 1.26 | 2.01 | 0.91 | 1.00 |
| 76 | CD62P | 1.27 | 1.75 | 0.91 | 1.00 |
| 77 | CD63 | 98.1 | 96.1 | 40.23 | 22.86 |
| 78 | CD64 | 1.49 | 1.87 | 1.00 | 1.00 |
| 79 | CD66 (a, c, d, e) | 1.57 | 13.7 | 1.10 | 1.67 |
| 80 | CD66b | 1.61 | 1.97 | 1.00 | 1.08 |
| 81 | CD66f | 1.58 | 2.37 | 0.91 | 1.08 |
| 82 | CD69 | 1.17 | 2.05 | 0.91 | 1.00 |
| 83 | CD70 | 2.14 | 3.79 | 1.10 | 1.17 |
| 84 | CD71 | 54.2 | 96 | 5.82 | 19.81 |
| 85 | CD72 | 1.92 | 1.84 | 1.00 | 1.00 |
| 86 | CD73 | 99.9 | 99.1 | 236.09 | 171.04 |
| 87 | CD74 | 1.6 | 2.11 | 1.10 | 1.17 |
| 88 | CD75 | 1.25 | 1.17 | 0.73 | 0.84 |
| 89 | CD77 | 1.45 | 1.29 | 0.91 | 0.84 |
| 90 | CD79b | 5.18 | 1.8 | 1.30 | 1.00 |
| 91 | CD80 | 2.73 | 2.5 | 1.00 | 1.00 |

| | | | | | |
|---|---|---|---|---|---|
| 92 | CD81 | 99.9 | 100 | 271.56 | 327.21 |
| 93 | CD83 | 1.48 | 1.99 | 0.91 | 1.00 |
| 94 | CD84 | 1.37 | 2.24 | 0.91 | 1.00 |
| 95 | CD85 | 1.08 | 2.31 | 0.82 | 1.00 |
| 96 | CD86 | 1.36 | 2.11 | 0.91 | 1.00 |
| 97 | CD87 | 7.27 | 2.38 | 1.70 | 1.08 |
| 98 | CD88 | 1.02 | 1.45 | 0.80 | 0.83 |
| 99 | CD89 | 1.3 | 1.12 | 0.80 | 0.83 |
| 100 | CD90 | 99.9 | 100 | 328.28 | 500.91 |
| 101 | CD91 | 90.5 | 96.9 | 18.13 | 19.81 |
| 102 | CDw93 | 1.85 | 2.02 | 0.91 | 1.00 |
| 103 | CD94 | 1.36 | 2.1 | 0.91 | 1.00 |
| 104 | CD95 | 85.7 | 99.4 | 11.48 | 62.47 |
| 105 | CD97 | 1.5 | 1.58 | 0.91 | 0.92 |
| 106 | CD98 | 99.8 | 99.9 | 184.30 | 125.19 |
| 107 | CD99 | 100 | 99.9 | 245.16 | 272.27 |
| 108 | CD99R | 96.2 | 99.6 | 25.96 | 55.75 |
| 109 | CD100 | 1.14 | 1.38 | 0.80 | 0.83 |
| 110 | CD102 | 4.22 | 5.2 | 1.20 | 1.17 |
| 111 | CD103 | 1.31 | 1.32 | 1.00 | 0.92 |
| 112 | CD105 | 99.9 | 99.9 | 58.98 | 69.16 |
| 113 | CD106 | 30.3 | 27.9 | 2.51 | 2.08 |
| 114 | CD107a | 47.8 | 21.4 | 4.91 | 2.58 |
| 115 | CD107b | 33.2 | 8.92 | 3.71 | 1.83 |

| | | | | | |
|---|---|---|---|---|---|
| 116 | CD108 | 32.9 | 21.7 | 3.51 | 2.50 |
| 117 | CD109 | 97.8 | 99.8 | 26.25 | 94.35 |
| 118 | CD112 | 4.9 | 28.6 | 2.01 | 3.50 |
| 119 | CD114 | 1.38 | 1.66 | 0.91 | 1.00 |
| 120 | CD116 | 1.1 | 4.11 | 0.82 | 1.38 |
| 121 | CD117 | 1.25 | 1.68 | 0.80 | 0.92 |
| 122 | CD118 | 1.18 | 1.32 | 0.91 | 0.83 |
| 123 | CD119 | 26 | 41.6 | 3.51 | 4.17 |
| 124 | CD120a | 10.9 | 5.3 | 1.70 | 1.58 |
| 125 | CD121a | 2.05 | 7.6 | 1.10 | 1.75 |
| 126 | CD121b | 1.16 | 1.54 | 0.91 | 0.92 |
| 127 | CD122 | 1.14 | 1.8 | 0.80 | 0.92 |
| 128 | CD123 | 1.15 | 1.41 | 0.80 | 0.92 |
| 129 | CD124 | 1.54 | 1.37 | 1.00 | 1.00 |
| 130 | CD126 | 32 | 21.1 | 3.41 | 2.42 |
| 131 | CD127 | 1.21 | 1.59 | 0.91 | 1.00 |
| 132 | CD128b | 1.57 | 1.52 | 1.00 | 0.92 |
| 133 | CD130 | 9.44 | 15.8 | 2.30 | 2.58 |
| 134 | CD134 | 1.42 | 1.55 | 1.00 | 0.92 |
| 135 | CD135 | 1.07 | 1.48 | 0.80 | 0.92 |
| 136 | CD137 | 0.96 | 2.5 | 0.91 | 1.00 |
| 137 | CD137 Ligand | 1.29 | 1.55 | 0.80 | 0.92 |
| 138 | CD138 | 39.3 | 45 | 3.71 | 4.08 |
| 139 | CD140a | 77.6 | 95.9 | 11.48 | 27.99 |

| | | | | | |
|---|---|---|---|---|---|
| 140 | CD140b | 95.6 | 98.6 | 48.83 | 69.87 |
| 141 | CD141 | 68.4 | 78.3 | 9.61 | 11.30 |
| 142 | CD142 | 70.6 | 37.2 | 10.08 | 3.50 |
| 143 | CD144 | 0.98 | 1.21 | 0.80 | 0.92 |
| 144 | CD146 | 99.8 | 98.6 | 83.75 | 38.77 |
| 145 | CD147 | 99.8 | 100 | 356.72 | 363.90 |
| 146 | CD150 | 1.34 | 1.44 | 0.80 | 0.92 |
| 147 | CD151 | 99.7 | 92.5 | 58.75 | 35.71 |
| 148 | CD152 | 1.28 | 1.2 | 0.91 | 0.92 |
| 149 | CD153 | 1.64 | 1.07 | 0.80 | 0.92 |
| 150 | CD154 | 1.34 | 1.31 | 0.91 | 0.83 |
| 151 | CD158a | 1.1 | 1.37 | 0.82 | 0.84 |
| 152 | CD158b | 1.28 | 1.7 | 0.82 | 1.00 |
| 153 | CD161 | 1.19 | 1.48 | 0.91 | 1.00 |
| 154 | CD162 | 1.47 | 2.46 | 0.91 | 1.17 |
| 155 | CD163 | 1.25 | 1.6 | 0.80 | 0.83 |
| 156 | CD164 | 99.7 | 97.6 | 102.89 | 34.87 |
| 157 | CD165 | 99.3 | 99.5 | 26.25 | 26.36 |
| 158 | CD166 | 99.7 | 99.8 | 92.11 | 88.70 |
| 159 | CD171 | 2.25 | 1.28 | 0.91 | 0.92 |
| 160 | CD172b | 1.25 | 1.7 | 0.91 | 0.92 |
| 161 | CD177 | 1.43 | 1.37 | 0.91 | 0.92 |
| 162 | CD178 | 1.04 | 1.7 | 0.91 | 0.83 |
| 163 | CD180 | 1.37 | 1.62 | 0.91 | 1.00 |

| | | | | | |
|---|---|---|---|---|---|
| 164 | CD181 | 1.79 | 4.44 | 0.91 | 1.42 |
| 165 | CD183 | 2.11 | 1.81 | 0.91 | 1.00 |
| 166 | CD184 | 1.46 | 1.23 | 1.00 | 0.83 |
| 167 | CD193 | 1.61 | 1.58 | 0.91 | 0.92 |
| 168 | CD195 | 1.8 | 1.35 | 0.91 | 0.92 |
| 169 | CD196 | 1.36 | 1.35 | 0.80 | 0.92 |
| 170 | CD197 | 1.85 | 3.61 | 0.82 | 1.08 |
| 171 | CD200 | 2.02 | 1.82 | 0.80 | 0.83 |
| 172 | CD205 | 1.21 | 1.62 | 0.82 | 0.92 |
| 173 | CD206 | 1.24 | 1.33 | 1.00 | 1.00 |
| 174 | CD209 | 2.34 | 2.84 | 0.91 | 1.08 |
| 175 | CD220 | 10.3 | 1.75 | 2.41 | 1.00 |
| 176 | CD221 | 70.1 | 21 | 7.23 | 3.08 |
| 177 | CD226 | 1.46 | 1.43 | 0.91 | 0.92 |
| 178 | CD227 | 64.8 | 97.2 | 7.43 | 34.68 |
| 179 | CD229 | 1.11 | 1.51 | 0.80 | 0.83 |
| 180 | CD231 | 1.82 | 1.26 | 1.00 | 0.92 |
| 181 | CD235a | 1.5 | 1.46 | 0.91 | 0.92 |
| 182 | CD243 | 2.22 | 2.43 | 0.91 | 1.17 |
| 183 | CD244 | 0.9 | 0.86 | 0.91 | 0.83 |
| 184 | CD255 | 1.4 | 1.55 | 1.00 | 0.83 |
| 185 | CD268 | 1.47 | 0.98 | 0.91 | 0.83 |
| 186 | CD271 | 1.35 | 12.9 | 0.91 | 1.58 |
| 187 | CD273 | 34.6 | 12.4 | 3.71 | 1.75 |

| | | | | | |
|---|---|---|---|---|---|
| 188 | CD274 | 42 | 22.2 | 4.41 | 2.67 |
| 189 | CD275 | 1.56 | 1.98 | 0.91 | 1.00 |
| 190 | CD278 | 1.11 | 1.9 | 0.80 | 1.08 |
| 191 | CD279 | 1.53 | 2.1 | 0.91 | 1.00 |
| 192 | CD282 | 1.82 | 3.32 | 1.00 | 1.25 |
| 193 | CD305 | 1.51 | 2.05 | 1.00 | 1.25 |
| 194 | CD309 | 0.91 | 1.99 | 0.91 | 1.00 |
| 195 | CD314 | 1.16 | 1.53 | 0.91 | 0.92 |
| 196 | CD321 | 2.94 | 3.98 | 1.10 | 1.17 |
| 197 | CDw327 | 1.28 | 2.05 | 0.80 | 1.08 |
| 198 | CDw328 | 0.99 | 2.32 | 0.91 | 1.08 |
| 199 | CD329 | 1.39 | 2.75 | 0.91 | 1.08 |
| 200 | CD335 | 0.94 | 2.75 | 0.91 | 1.25 |
| 201 | CD336 | 1 | 2.1 | 0.91 | 1.08 |
| 202 | CD337 | 1.31 | 2.11 | 0.91 | 1.00 |
| 203 | CD338 | 1.18 | 2.36 | 0.82 | 1.08 |
| 204 | CD340 | 86.4 | 97.9 | 10.08 | 18.12 |
| 205 | αβTCR | 1.79 | 1.56 | 0.82 | 0.92 |
| 206 | β2-Microglobulin | 88.3 | 44.5 | 14.68 | 3.92 |
| 207 | BLTR-1 | 1.96 | 2.01 | 1.10 | 1.00 |
| 208 | CLIP | 1.62 | 1.84 | 1.00 | 1.00 |
| 209 | CMRF-44 | 1.78 | 1.86 | 1.00 | 1.00 |
| 210 | CMRF-56 | 2.56 | 2.71 | 1.00 | 1.00 |
| 211 | EGF receptor | 91.5 | 97 | 14.96 | 23.38 |

| | | | | | |
|---|---|---|---|---|---|
| 212 | fMLP receptor | 1.67 | 2.23 | 1.00 | 1.00 |
| 213 | γσTCR | 1.79 | 2.49 | 1.10 | 1.08 |
| 214 | HPC | 2.74 | 7.18 | 1.10 | 1.33 |
| 215 | HLA-ABC | 99.3 | 98 | 47.89 | 24.22 |
| 216 | HLA-A2 | 1.24 | 1.76 | 0.82 | 0.92 |
| 217 | HLA-DQ | 79.4 | 74.4 | 15.63 | 7.86 |
| 218 | HLA-DR | 1.63 | 1.84 | 0.91 | 1.00 |
| 219 | HLA-DR, -DP, -DQ | 1.84 | 1.85 | 0.91 | 1.00 |
| 220 | Invariant NK T | 1.12 | 1.64 | 0.91 | 0.92 |
| 221 | Disialoganglioside GD2 | 56.8 | 90.9 | 7.02 | 77.47 |
| 222 | MIC A/B | 24.5 | 11.3 | 2.80 | 1.67 |
| 223 | NKB1 | 1.49 | 1.78 | 1.00 | 1.00 |
| 224 | SSEA-1 | 1.7 | 1.66 | 0.91 | 1.00 |
| 225 | SSEA-4 | 51.1 | 87.4 | 5.87 | 25.00 |
| 226 | TRA-1-60 | 1.35 | 1.39 | 0.91 | 0.92 |
| 227 | TRA-1-81 | 1.54 | 1.51 | 0.91 | 0.92 |
| 228 | Vβ 23 | 2.26 | 2.21 | 1.00 | 1.00 |
| 229 | Vβ 8 | 1.49 | 2.18 | 0.91 | 1.00 |
| 230 | CD326 | 3.47 | 3.19 | 1.10 | 1.17 |
| 231 | mIgM | 1.94 | 1.81 | 1.94 | 1.81 |
| 232 | mIgG1 | 1.81 | 1.92 | 1.81 | 1.92 |
| 233 | mIgG2a | 1.61 | 1.72 | 1.61 | 1.72 |
| 234 | mIgG2b | 1.91 | 1.82 | 1.91 | 1.82 |

| | | | | | |
|---|---|---|---|---|---|
| 235 | mIgG3 | 1.51 | 1.95 | 1.51 | 1.95 |
| 236 | CD49f | 82.4 | 70.2 | 18.71 | 7.08 |
| 237 | CD104 | 1.77 | 4.16 | 1.00 | 1.18 |
| 238 | CD120b | 2.23 | 1.86 | 0.89 | 1.00 |
| 239 | CD132 | 2.14 | 2.32 | 1.00 | 1.09 |
| 240 | CD201 | 33.9 | 85 | 4.23 | 14.33 |
| 241 | CD210 | 1.63 | 1.81 | 1.00 | 0.92 |
| 242 | CD212 | 1.31 | 1.84 | 1.00 | 0.92 |
| 243 | CD267 | 1.42 | 2.05 | 0.89 | 0.92 |
| 244 | CD294 | 1.58 | 4.43 | 0.89 | 1.08 |
| 245 | SSEA-3 | 1.77 | 2.35 | 0.89 | 1.20 |
| 246 | CLA | 2.17 | 2.85 | 1.00 | 1.20 |
| 247 | Integrin β7 | 1.31 | 2.24 | 1.00 | 1.00 |
| 248 | rIgM | 1.98 | 1.77 | 1.98 | 1.77 |
| 249 | rIgG1 | 1.78 | 1.78 | 1.78 | 1.78 |
| 250 | rIgG2a | 1.55 | 1.86 | 1.55 | 1.86 |
| 251 | rIgG2b | 1.87 | 1.95 | 1.87 | 1.95 |

nMFI, normalized median fluorescence intensity; HS, high-serum; LS, low-serum

**Supplementary Table 2.** Scores for each item of the ICRS histological scores

| | Item | NT | HS | LS |
|---|---|---|---|---|
| After 4 weeks | Ti | 2.9 ± 0.6 | 2.9 ± 0.2 | 3.3 ± 0.7 |
| | Matx | 2.1 ± 0.7 | 1.9 ± 0.5 | 2.5 ± 1.1 |
| | Stru | 2.8 ± 0.4 | 3.2 ± 0.4 | 3.2 ± 0.6 |
| | Clus | 1.8 ± 0.4 | 2.0 ± 0.1 | 2.1 ± 0.4 |
| | Tide | 2.7 ± 0.7 | 3.3 ± 0.7 | 3.8 ± 0.6 |
| | Bform | 2.6 ± 0.3 | 2.5 ± 0.1 | 2.7 ± 0.3 |
| | SurfH | 2.5 ± 0.4 | 2.6 ± 0.6 | 2.7 ± 0.5 |
| | FilH | 2.4 ± 1.0 | 2.9 ± 0.6 | 3.2 ± 1.3 |
| | Latl | 2.4 ± 0.6 | 2.4 ± 0.3 | 2.6 ± 0.4 |
| | Basl | 2.1 ± 0.5 | 2.3 ± 0.5 | 3.0 ± 0.6 |
| | InfH | 5.0 ± 0.0 | 5.0 ± 0.0 | 5.0 ± 0.0 |
| | Hgtot | 29.3 ± 3.7 | 31.0 ± 2.8 | 34.0 ± 5.2 |
| After 12 weeks | Ti | 2.6 ± 0.7 | 3.4 ± 0.6 | 3.5 ± 0.5 |
| | Matx | 1.8 ± 0.8 | 2.8 ± 0.6 | 2.8 ± 1.1 |
| | Stru | 2.7 ± 0.6 | 3.3 ± 0.3 | 3.3 ± 0.4 |
| | Clus | 1.8 ± 0.3 | 2.0 ± 0.0 | 2.1 ± 0.2 |
| | Tide | 2.9 ± 0.7 | 3.4 ± 0.7 | 3.8 ± 0.4 |
| | Bform | 2.5 ± 0.0 | 2.5 ± 0.0 | 2.6 ± 0.2 |
| | SurfH | 2.1 ± 0.4 | 2.9 ± 0.2 | 2.8 ± 0.3 |
| | FilH | 2.9 ± 1.1 | 3.7 ± 0.9 | 3.8 ± 0.7 |
| | Latl | 1.8 ± 0.8 | 2.9 ± 0.2 | 2.8 ± 0.4 |

| | | | | |
|---|---|---|---|---|
| | Basl | 2.4 ± 0.7 | 2.8 ± 0.4 | 2.9 ± 0.4 |
| | InfH | 5.0 ± 0.0 | 5.0 ± 0.0 | 5.0 ± 0.0 |
| | Hgtot | 28.5 ± 5.5 | 34.7 ± 3.2 | 35.4 ± 4.1 |

NT, no-treatment; HS, high-serum; LS, low-serum; Ti, tissue morphology; Matx, matrix staining; Stru, structural integrity; Clus, cluster formation; Tide, tidemark opening; Bform, bone formation; SurfH, histological appraisal of surface architecture; FilH, histological appraisal of the degree of defect filling; Latl, lateral integration of defect-filling tissue; Basl, basal integration of defect-filling tissue; InfH, histological signs of inflammation; Hgtot, histological total score

**Supplementary Table 3.** List of primary and secondary antibodies used for immunohistochemistry of PDC and iPSC sheets and histological sections from the animal study

| Antibody | Dilution | Company | Catalog number |
|---|---|---|---|
| Aggrecan | 1:100 | R&D Systems | AF1220 |
| Fibronectin | 1:500 | Thermo Fisher | MA5-11981 |
| Type I collagen | 1:100 | Southern Biotech | 1310-01 |
| Type II collagen | 1:100 | Kyowa Pharma Chemical | F-57 |
| Human-specific vimentin | 1:100 | Abcam | Ab16700 |
| Alexa Fluor 488-conjugated goat anti-mouse IgG | 1:200 | Thermo Fisher | A32723 |
| Alexa Fluor 488-conjugated goat anti-rabbit IgG | 1:200 | Thermo Fisher | A32731 |

**Supplementary Table 4.** List of antibodies used for flow cytometry in single stain analysis

| Specificity | Fluorochrome | Clone | Company |
|---|---|---|---|
| CD9 | PE | M-L13 | BD |
| CD10 | PE | HI10a | BD |
| CD13 | PE | WM15 | BD |
| CD14 | APC | M5E2 | BD |
| CD19 | PE | HIB19 | BD |
| CD26 | FITC | M-A261 | BD |
| CD29 | PE | MAR4 | BD |
| CD31 | FITC | 5.6E | Beckman Coulter |
| CD34 | PE | 581 | BD |
| CD44 | FITC | G44-26 | BD |
| CD45 | FITC | J33 | Beckman Coulter |
| CD49a | PE | SR84 | BD |
| CD49b | FITC | AK-7 | BD |
| CD49d | PE | 9F10 | BD |
| CD49f | PE | GoH3 | BD |
| CD56 | APC | B159 | BD |
| CD71 | FITC | M-A712 | BD |
| CD73 | FITC | AD2 | BD |
| CD81 | APC | JS-81 | BD |
| CD90 | APC | 5E10 | BD |
| CD91 | PE | A2MR-α2 | BD |

| | | | |
|---|---|---|---|
| CD95 | FITC | DX2 | BD |
| CD99 | FITC | TÜ12 | BD |
| CD105 | PE | 266 | BD |
| CD106 | FITC | 51-10C9 | BD |
| CD107a | FITC | H4A3 | BD |
| CD107b | FITC | H4B4 | BD |
| CD108 | PE | KS-2 | BD |
| CD120a | APC | W15099A | BioLegend |
| CD130 | BB700 | AM64 | BD |
| CD140a | PE | αR1 | BD |
| CD140b | PE | 28D4 | BD |
| CD146 | FITC | P1H12 | BD |
| CD164 | PE | N6B6 | BD |
| CD165 | BB700 | SN2 | BD |
| CD166 | PE | 3A6 | BD |
| CD201 | PE | RCR-252 | BD |
| CD205 | PE | MG38 | BD |
| CD227 | FITC | HMPV | BD |
| β2-Microglobulin | PE | TÜ99 | BD |
| EGFR | PE | EGFR.1 | BD |
| GD2 | BB700 | 14.G2a | BD |
| HLA-ABC | APC | G46-2.6 | BD |
| HLA-DR | PE | G46-6 | BD |
| HLA DR,DP,DQ | FITC | Tu39 | BD |

| | | | |
|---|---|---|---|
| SSEA-4 | PE | MC813-70 | BD |
| STRO-1 | FITC | unknown | Santa Cruz Biotechnology |
| mIgG1 | FITC | 679.1Mc7 | Beckman Coulter |
| mIgG2a | FITC | G155-178 | BD |
| mIgG2b | FITC | MG2b-57 | BioLegend |
| mIgM | FITC | G155-228 | BD |
| mIgG1 | PE | 679.1Mc7 | Beckman Coulter |
| mIgG2a | PE | G155-178 | BD |
| mIgG2b | PE | 27-35 | BD |
| mIgG3 | PE | A112-3 | BD |
| Rat IgG1 | PE | MRG1-58 | BioLegend |
| Rat IgG2a | PE | MRG2a-83 | BioLegend |
| mIgM | PE | G155-228 | BD |
| mIgG1 | APC | 679.1Mc7 | Beckman Coulter |
| mIgG2a | APC | G155-178 | BD |
| mIgG1 | BB700 | X40 | BD |
| mIgG2a | BB700 | G155-178 | BD |

**Supplementary Table 5.** List of primers used for RT-PCR

| Target | Forward Primer | Reverse Primer |
|---|---|---|
| ACAN | AGGAGACAGAGGGACACGTC | TCCACTGGTAGTCTTGGGCAT |
| CHI3L1 | GCAACACTGACTATGCTGTGG | GAGTGAAGCTCCTCCCGAAG |
| CHI3L2 | CCCTTATCACTGGCCACAAC | CCACCTTCTCTGATGGCATT |
| COL10A1 | ATGCTGCCACAAATACCCTTT | GGTAGTGGGCCTTTTATGCCT |
| COL1A1 | GTCGAGGGCCAAGACGAAG | CAGATCACGTCATCGCACAAC |
| COL2A1 | GTGGAGCAGCAAGAGCAA | TGTTGGGAGCCAGATTGT |
| COMP | GATCACGTTCCTGAAAAACA | GCTCTCCGTCTGGATGCAG |
| DKK1 | AGTACTGCGCTAGTCCCACC | TCCTCAATTTCTCCTCGGAA |
| ESM1 | AAGGCTGCTGATGTAGTTC | GCTATTTATGGAAGTGTATGTGTTT |
| GREM1 | CGTGTGAAGCAGTGTCGTTG | CTCATGCACACGAACTACGC |
| MIA | CACAGAGCCTCGCCTTTGCC | TGACCCATGCCCACCATCAC |
| MMP13 | ACTGAGAGGCTCCGAGAAATG | GAACCCCGCATCTTGGCTT |
| MMP3 | ATGATGAACAATGGACAAAGGA | GAGTGAAAGAGACCCAGGGA |
| RPL13A | CTCAAGGTGTTTGACGGCATCC | TACTTCCAGCCAACCTCGTGAG |
| RUNX2 | ACCATGGTGGAGATCATCG | CGCCATGACAGTAACCACAG |
| SOX9 | AACGCCGAGCTCAGCAAGA | CCGCGGCTGGTACTTGTAATC |
| TGFB1 | AGCGACTCGCCAGAGTGGTTA | GCAGTGTGTTATCCCTGCTGTCA |

**Supplementary Table 6.** ICRS histological grading system

| Item | Score | Description |
|---|---|---|
| Ti: tissue morphology | 4 | Mostly hyaline cartilage |
| | 3 | Mostly fibrocartilage |
| | 2 | Mostly noncartilage |
| | 1 | Exclusively noncartilage |
| Matx: matrix staining | 4 | Strong |
| | 3 | Moderate |
| | 2 | Slight |
| | 1 | None |
| Stru: structural integrity | 5 | Normal, similar to healthy mature cartilage |
| | 4 | Beginning of columnar organization of chondrocytes |
| | 3 | No organization of chondrocytes |
| | 2 | Cysts or disruptions |
| | 1 | Severe disintegration |
| Clus: cluster formation | 3 | No clusters |
| | 2 | <25% of the cells clustered |
| | 1 | 25–100% of the cells clustered |
| Tide: tidemark opening | 5 | Complete intactness of the calcified cartilage layer |
| | 4 | 76–90% of the calcified cartilage layer intact |
| | 3 | 50–75% of the calcified cartilage layer intact |
| | 2 | 25–49% of the calcified cartilage layer intact |
| | 1 | <25% of the calcified cartilage layer intact |

| | | |
|---|---|---|
| Bform: bone formation | 3 | Strong |
| | 2 | Slight |
| | 1 | No formation |
| SurfH: histological appraisal of surface architecture | 4 | Normal |
| | 3 | Slight fibrillation or irregularity |
| | 2 | Moderate fibrillation or irregularity |
| | 1 | Severe fibrillation or disruption |
| FilH: histological appraisal of the degree of defect filling | 5 | 91–110% |
| | 4 | 76–90% |
| | 3 | 51–75% |
| | 2 | 25–50% |
| | 1 | <25% |
| Latl: lateral integration of defect-filling tissue | 3 | Bonded at both sides |
| | 2 | Bonded at one end/partially both ends |
| | 1 | Not bonded |
| Basl: basal integration of defect-filling tissue | 4 | 91–100% |
| | 3 | 71–90% |
| | 2 | 50–70% |
| | 1 | <50% |
| InfH: histological signs of inflammation | 5 | No inflammation |
| | 3 | Slight inflammation |
| | 1 | Strong inflammation |
| Hgtot: histological total score | 45 | |
